# Time elapsed between Zika and dengue virus infections affects antibody and T cell responses

**DOI:** 10.1101/621094

**Authors:** Erick X. Pérez-Guzmán, Petraleigh Pantoja, Crisanta Serrano-Collazo, Mariah A. Hassert, Alexandra Ortiz-Rosa, Idia V. Rodríguez, Luis Giavedoni, Vida Hodara, Laura Parodi, Lorna Cruz, Teresa Arana, Laura J. White, Melween I. Martínez, Daniela Weiskopf, James D. Brien, Aravinda de Silva, Amelia K. Pinto, Carlos A. Sariol

## Abstract

The role of Zika virus (ZIKV) immunity on subsequent dengue virus (DENV) infections is relevant to anticipate the dynamics of forthcoming DENV epidemics in areas with previous ZIKV exposure. We study the effect of ZIKV infection with various strains on subsequent DENV immune response after 10 and 2 months of ZIKV infection in rhesus macaques. Our results show that a subsequent DENV infection in animals with early- and middle-convalescent periods to ZIKV do not promote an increase in DENV viremia nor pro-inflammatory status. Previous ZIKV exposure increases the magnitude of the antibody and T cell responses against DENV, and different time intervals between infections alter the magnitude and durability of such responses—more after longer ZIKV pre-exposure. Collectively, we find no evidence of a detrimental effect of ZIKV immunity in a subsequent DENV infection. This supports the implementation of ZIKV vaccines that could also boost immunity against future DENV epidemics.

---

Zika virus (ZIKV) is a re-emerging mosquito-borne *Flavivirus* that has captivated the attention of the scientific community by its explosive spread in The Americas^1^, and severe neurological sequelae following infection ^2–4^. ZIKV established itself in tropical and sub-tropical regions that are endemic to other closely-related flaviviruses such as dengue virus (DENV). Both viruses belong to the Flaviviridae family and are transmitted by *Aedes spp.* mosquitoes. DENV is a global public health threat, having two-thirds of world’s population at risk of infection, causing ∼390 million infections annually^5,6^. DENV exists as four genetically similar but antigenically different serotypes (DENV1-4)^7^. Exposure to one DENV serotype confers long-lived immunity against a homotypic secondary infection. However, secondary infection with a heterologous serotype of DENV is the major risk factor to induce severe DENV disease ^8–10^.

Due to the antigenic similarities between DENV and ZIKV, concerns have been raised regarding the impact of DENV-ZIKV cross-reactive immunity on the development of severe clinical manifestations^11,12^. It has been demonstrated that DENV-immune sera from humans can enhance ZIKV infection *in vitro*^13,14^, and *in vivo* in immune-deficient mouse models^15^. However, recent results from our group and others have shown that previous flavivirus exposure— including DENV—may have no detrimental impact on ZIKV infection *in vivo* in non-human primates (NHP)^16,17^ and humans^18^. Moreover, these studies and others suggest that previous DENV immunity may play a protective role during ZIKV infection involving humoral and cellular responses^19–23^. On the other hand, little is known about the opposite scenario, the role of a previous ZIKV exposure on subsequent DENV infection, which is relevant to anticipate the dynamics of forthcoming DENV epidemics.

The recent ZIKV epidemic in the Americas resulted in the development of a herd immunity that may have an impact in subsequent infections with other actively circulating flaviviruses such as DENV. Thus, human sub-populations such as newborns, international travelers from non-flavivirus endemic areas or DENV-naïve subjects could be exposed to a ZIKV infection prior to DENV—since DENV declined in the Americas during ZIKV epidemic^24^. After the epidemic, herd immunity reduced ZIKV transmission and DENV will re-emerge and potentially infect these DENV-naïve ZIKV-immune sub-populations in The Americas or potentially in other geographic areas newly at risk^25,26^. An epidemiological study based on active DENV surveillance in Salvador, Brazil, suggests that the reduction of DENV cases after the ZIKV epidemic is due to protection from cross-reactive immune responses between these viruses^27^. Prospective experimental studies are needed to confirm this hypothesis. NHPs provide advantages such as an immune response comparable to humans, and the normalization of age, sex, injection route, viral inoculum and timing of infection^28^. Although clinical manifestations by flaviviral infections are limited in NHPs^29^, they have been widely used as an advanced animal model for the study of DENV and ZIKV immune response, pathogenesis, and vaccine development^16,17,28, 30–33^.

ZIKV antibodies (Abs) are capable of enhancing DENV infection *in vitro*^34^. Characterization of the specificity of DENV and ZIKV cross-reactive response revealed that ZIKV monoclonal Abs and maternally acquired ZIKV Abs can increase DENV severity and viral burden in immune-deficient mouse models^35,36^. *George et al.*, showed that an early convalescence to ZIKV induced a significant higher peak of DENV viremia and a pro-inflammatory profile compared to ZIKV-naïve status in rhesus macaques^37^. A recent NHPs study showed that clinical and laboratory parameters of ZIKV-immune animals were not associated with an enhancement of DENV-2 infection. However, a higher peak of DENV-2 plasma RNAemia in ZIKV-immune animals was observed compared to DENV-2 serum RNAemia loads in control animals, but the use of different sample types may account for these differences^38^. Despite these findings, further studies are needed to dissect the complementary role of the innate, humoral and cellular immune response to mechanistically explain these findings. Particularly, there is no evidence of the modulation and functionality of the T cell response in the ZIKV-DENV scenario. Available studies rely upon pathogenesis and antibody studies, but there is no documented evidence as to whether cell-mediated immunity (CMI)— specifically the functional response of T cells—is modulated in a subsequent DENV infection by the presence of ZIKV immune memory.

Shorter time interval between DENV infections result in a subclinical secondary infection, while symptomatic secondary infections and severe DENV cases have been related with longer periods between infections^39–42^. These findings suggest that high titers of cross-reactive Abs play a time-dependent protective role between heterotypic DENV infections. Despite this evidence from DENV sequential infections, it remains poorly understood if the same applies to the time interval between ZIKV-DENV sequential infections. So currently, the role of multiple convalescent periods to ZIKV in the outcome of DENV and other flavivirus infections is in the forefront of discussions based on the limited studies available in experimental models and a lack of characterized human prospective cohorts of this scenario yet^27,43–45^.

To address these knowledge gaps, the objective of our study is to investigate the immune modulatory role of an early- and middle-convalescence after ZIKV infection on the outcome of a subsequent DENV infection in rhesus macaques. To test this, NHP cohorts who were ZIKV immune for 10 months (middle-convalescence), 2 months (early-convalescence) or naïve for ZIKV were exposed to DENV. The 2 months cohort was selected for direct comparison with previous work in NHP^37^, while the 10 months cohort was selected based on availability and to test a longer period of convalescence to ZIKV, where the cross-reactive Abs are known to wane ^46^. In each of these groups we assess DENV pathogenesis, the elicited Ab response, and characterized the CMI. Based on our knowledge, this is the first characterization of CMI with this scenario in NHPs—taking into account the synergistic effect between the Ab and cell-mediated responses. This study provides evidence that the presence of ZIKV immune memory contributes to increase the magnitude of the immune response—more efficient after longer ZIKV pre-exposure—against a DENV infection, without promoting enhancement of DENV viremia nor inducing higher levels of pro-inflammatory cytokines.

## Results

### DENV challenge and clinical status of rhesus macaques

The experimental design includes three cohorts of rhesus macaques (*Macaca mulatta*), within the age range considered as young adults (Supplementary Fig. 1k), that were challenged with DENV-2 (NGC-44 strain), monitored and bled for three months (Fig. 1). Two cohorts were previously exposed to ZIKV: cohort 1 (ZIKVPF-10mo) was comprised of 4 animals that had been exposed to ZIKV H/PF/2013 strain 10 months before DENV-2 challenge (mid-convalescence), and cohort 2 (ZIKVPR-2mo) comprised of 6 animals that had been exposed to ZIKV PRVABC59 strain two months before DENV-2 challenge (early-convalescence). Both ZIKV strains used for previous exposure of these groups are >99.99% comparable in amino acid identity (Supplementary Table 1). An additional cohort 3 (Naïve) included four animals naïve to ZIKV/DENV as a control group. After DENV challenge all macaques were extensively monitored and sample collection was performed at various timepoints up to 90 days post infection (dpi) for serum and PBMCs isolation.

**Figure 1.**
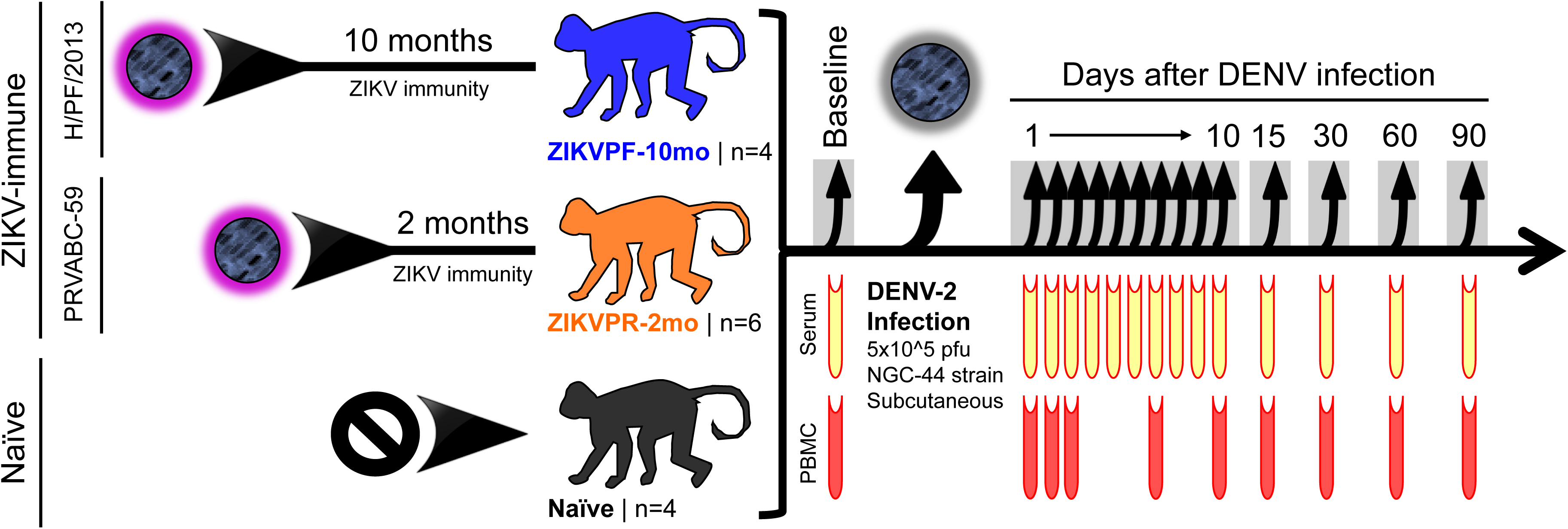
Experimental design for DENV-2 challenge of ZIKV-immune and naïve macaques. 14 young adult male rhesus macaques (*Macaca mulatta*), matched in age and weight, were divided in three cohorts. ZIKVPF-10mo (n=4): composed of four animals (5K6, CB52, 2K2, and 6N1) that were inoculated with 1×10^6^ pfu/500 ul of the ZIKV H/PF/2013 strain subcutaneously 10 months before (middle convalescence) DENV-2 challenge. ZIKVPR-2mo (n=6): composed of six animals (MA067, MA068, BZ34, MA141, MA143, and MA085) that were inoculated with 1×10^6^ pfu/500 ul of the contemporary ZIKV PRVABC59 strain two months before (early convalescence) DENV-2 challenge. Both ZIKV strains used for previous exposure of these groups are >99.99% comparable in amino acid identity (Supplementary Table 1). Naïve (n=4): composed of four ZIKV/DENV naïve animals (MA123, MA023, MA029, and MA062) as a control group. Prior to DENV-2 challenge all animals were subjected to quarantine period. All cohorts challenged subcutaneously (deltoid area) with 5×10^5^ pfu/500 ul of DENV-2 New Guinea 44 strain (NGC44). After DENV-2 challenge all animals were extensively monitored for evidence of disease and clinical status by vital signs such as external temperature (°C), weight (Kg), CBC and CMP panels at the Caribbean Primate Research Center (CPRC). Blood samples were collected at baseline, 1 to 10, 15, 30, 60 and 90 days after DENV infection. In all timepoints the blood samples were used for serum separation (yellow). PBMCs isolation (red) was performed in different tubes with citrate as anticoagulant at baseline, 1, 2, 3, 7, 10, 15, 30, 60, and 90 days after DENV infection.

The clinical status was monitored to determine if the presence of ZIKV immunity affected the clinical outcome of DENV infection. Vital signs such as weight (kg), and temperature (°C) were monitored. Also, complete blood cell counts (CBC), and comprehensive metabolic panel (CMP) were performed before (baseline: day 0) and after DENV infection at multiple timepoints (CBC: 0, 7, 15 dpi; CMP: 0, 7, 15, 30 dpi). Neither symptomatic manifestations nor significant differences in weight or temperature were observed in any of the animals after DENV infection up to 90 dpi (Supplementary Fig. 1a-b). Likewise, no significant differences between groups were detected in CBC parameters: white blood cells (WBC), lymphocytes (LYM), neutrophils (NEU), monocytes (MON), and platelets (PLT) after DENV infection compared to basal levels of each group (Supplementary Fig. 1c-g). CMP levels showed no differences in alkaline phosphatase and aspartate transaminase (AST) (Supplementary Fig. 1h-i). Although within the normal range, levels of alanine transaminase (ALT) were significantly higher in the ZIKVPR-2mo group compared to its baseline at 7 dpi (*p*=0.0379, Two-way Anova Dunnett test), but at 15 and 30 dpi values returned to baseline levels (Supplementary Fig. 1j). Overall, except for the isolated increase of ALT at 7 dpi in ZIKVPR-2mo, the clinical profile suggests that the presence of ZIKV-immunity did not significantly influence the clinical outcome of DENV infection.

### DENV RNAemia not enhanced by previous ZIKV immunity

RNAemia levels in NHPs serum were quantified by qRT-PCR at baseline, 1 to 10, and 15 dpi to determine if the presence of early- (ZIKVPR-2mo) or mid-convalescence (ZIKVPF-10mo) to ZIKV alters DENV RNAemia kinetics. No significant differences between groups were observed in detected levels of DENV genome copies per ml of serum overtime (Fig. 2a). We noted that in the ZIKVPF-10mo group 3 out of 4 animals were able to keep the RNAemia level below 10^3^ genome copies the next day after DENV infection. This group started an early clearance of the RNAemia at 7 dpi, with only 1 out of 4 animals having detectable levels by days 8 and 9 pi. For ZIKVPR-2mo and naïve animals, the clearance of detectable RNAemia started at 8 dpi, in 4 out of 6 and 1 out of 4 of the animals, respectively. Naïve animals had the most delayed clearance of RNAemia with at least half of the animals with detectable levels of viral RNA until day 9 pi. RNAemia was completely resolved in all animals by 10 dpi. In summary, ZIKVPF-10mo had 7.25, ZIKVPR-2mo 7.5, and naïve animals 8 mean days of detectable RNAemia after DENV infection (Fig. 2b). In addition, the area under the curve (AUC) was calculated but no statistically significance differences were observed in the RNAemia peak among groups (Supplementary Fig. 2). However, the AUC trend to be lower in both ZIKV-immune groups. In terms of the kinetics, a delay in the peak RNAemia set point was observed in both ZIKV-immune groups (switch from day 2 to days 5 and 6) followed by higher, but non-significant, levels compared to the naïve group, and a subsequent early RNAemia clearance in both ZIKV-immune groups. Together these results show that, although no statistically significant differences among groups were observed, previous immunity to ZIKV is not associated with an increase in DENV RNAemia; even more, a mid-convalescence to ZIKV tended to develop a shorter viremic period.

**Figure 2.**
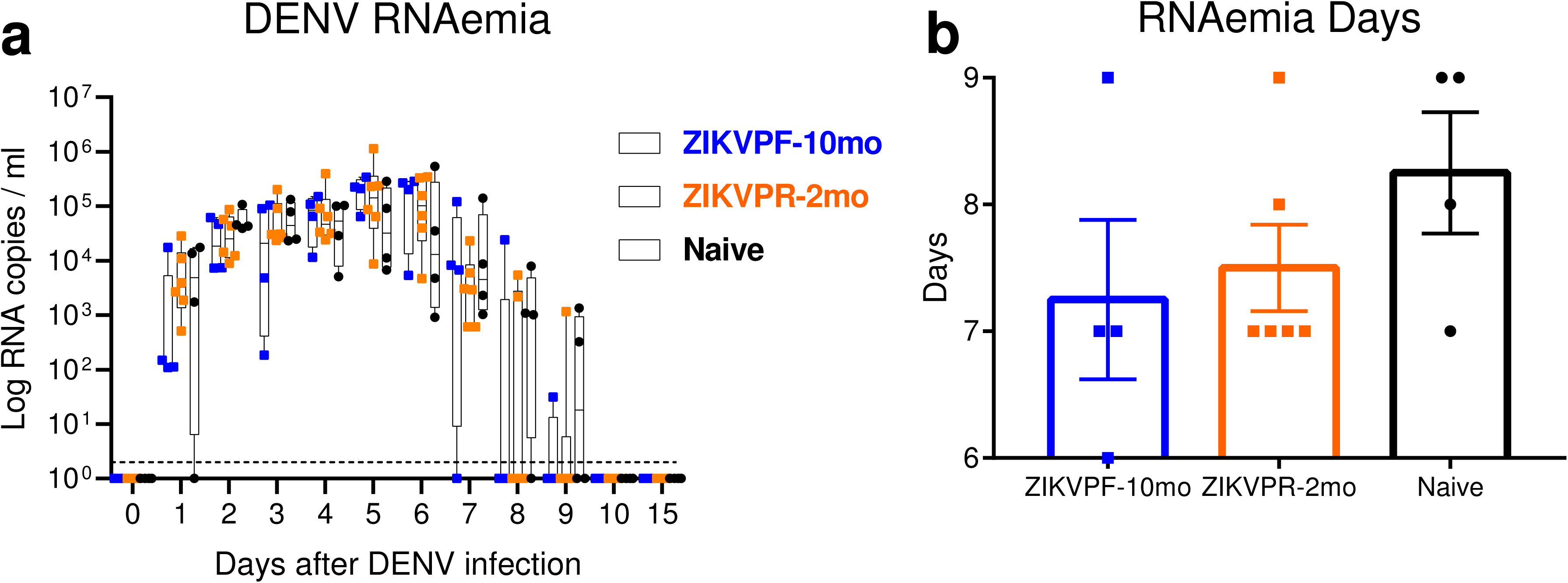
Previous ZIKV immunity does not contribute to an increase of DENV RNAemia. (**a**) DENV-2 RNA kinetics in ZIKV-immune and naïve animals at baseline, day 1 to day 10, and day 15 after DENV infection. RNA genome copies (Log10) per ml of serum were measured by qRT-PCR. Symbols represent individual animals per cohort: blue squares (ZIKVPF-10mo), orange squares (ZIKVPR-2mo) and black circles (Naïve). Box and whiskers show the distribution of log-transformed values per group per timepoint. Boxes include the mean value per group while whiskers depict the minimum and maximum values for each group. Cutted line mark the limit of detection (20 genomes copies). Statistically significant differences between groups were determined using Two-Way Anova adjusted for Tukey’s multiple comparisons test including 12 families, and 3 comparisons per family. (**b**) Total days that DENV-2 RNAemia was detected for each animal within cohorts. Bars represent mean days per cohort. Source data are provided as a Source Data file.

### Pro-inflammatory cytokines not exacerbated by ZIKV immunity

To determine if the characterized cytokine profile of an acute DENV infection was modulated by ZIKV immunity we assessed the serum concentration (pg/ml) of 8 cytokines/chemokines by Luminex multiplex at baseline, 1, 2, 3, 5, 10, 15 and 30 dpi. The naïve group showed significant higher levels of Type I interferon alpha (IFN-α) and pro-inflammatory cytokines such as Interleukin-6 (IL-6), and monokine induced by IFN-gamma (MIG/CXCL9) (Fig. 3a-c). IFN-α was highest at 5 dpi (Fig. 3a: *p*<0.0001 vs ZIKVPF-10mo and *p*=0.0003 vs ZIKVPR-2mo, Two-way Anova Tukey test). IFN-α has been demonstrated to be involved in the innate anti-viral immunity and elevated levels are associated with higher viral load and antigen availability. IL-6, a multifunctional cytokine involved in immune response regulation and many inflammatory reactions showed the highest levels at 1 dpi in naïve animals (Fig. 3b: *p*=0.0115 vs ZIKVPF-10mo and *p*=0.0185 vs ZIKVPR-2mo, Two-way ANOVA Tukey test). Finally, MIG/CXCL9, which is a potent chemoattractant involved in leucocyte trafficking demonstrated the highest levels at 10 dpi in naïve animals (Fig. 3c: *p*=0.0004 vs ZIKVPR-2mo, Two-way Anova Tukey test). On the other hand, the mid-convalescent ZIKVPF-10mo group showed higher levels of CXCL10 (IP-10) (Fig. 3g) at day 1 (*p*=0.0198 vs ZIKVPR-2mo, Two-way Anova Tukey test), 5 (*p*=0.0487 vs Naïve, Two-way Anova Tukey test) and 10 pi (*p*=0.0009 vs ZIKVPR-2mo, Two-way Anova Tukey test). CXCL10 is a T cell-activating chemokine and chemoattractant for many other immune cells. Also, this group showed higher levels of perforin (Fig. 3h) at day 10 (*p*=0.0024 vs Naïve and *p*=0.0190 vs ZIKVPR-2mo, Two-way Anova Tukey test) and 15 pi (*p*=0.0178 vs Naïve, Two-way Anova Tukey test). Perforin is an effector cytolytic protein released by activated cytotoxic CD8+ T cells and natural killer (NK) cells. No significant differences between groups were observed for other pro-inflammatory citokines such as monocyte chemoattractant protein 1 (MCP-1), macrophage inflammatory protein 1-beta (MIP-1β) and IL-1 receptor antagonist (IL-1RA) (Fig. 3d-f). Collectively, these results demonstrate that the presence of ZIKV immunity does not exacerbate pro-inflammatory status after DENV infection while mid-convalescence immunity to ZIKV stimulated levels of mediators mainly involved in the activation of cell-mediated immune response.

**Figure 3.**
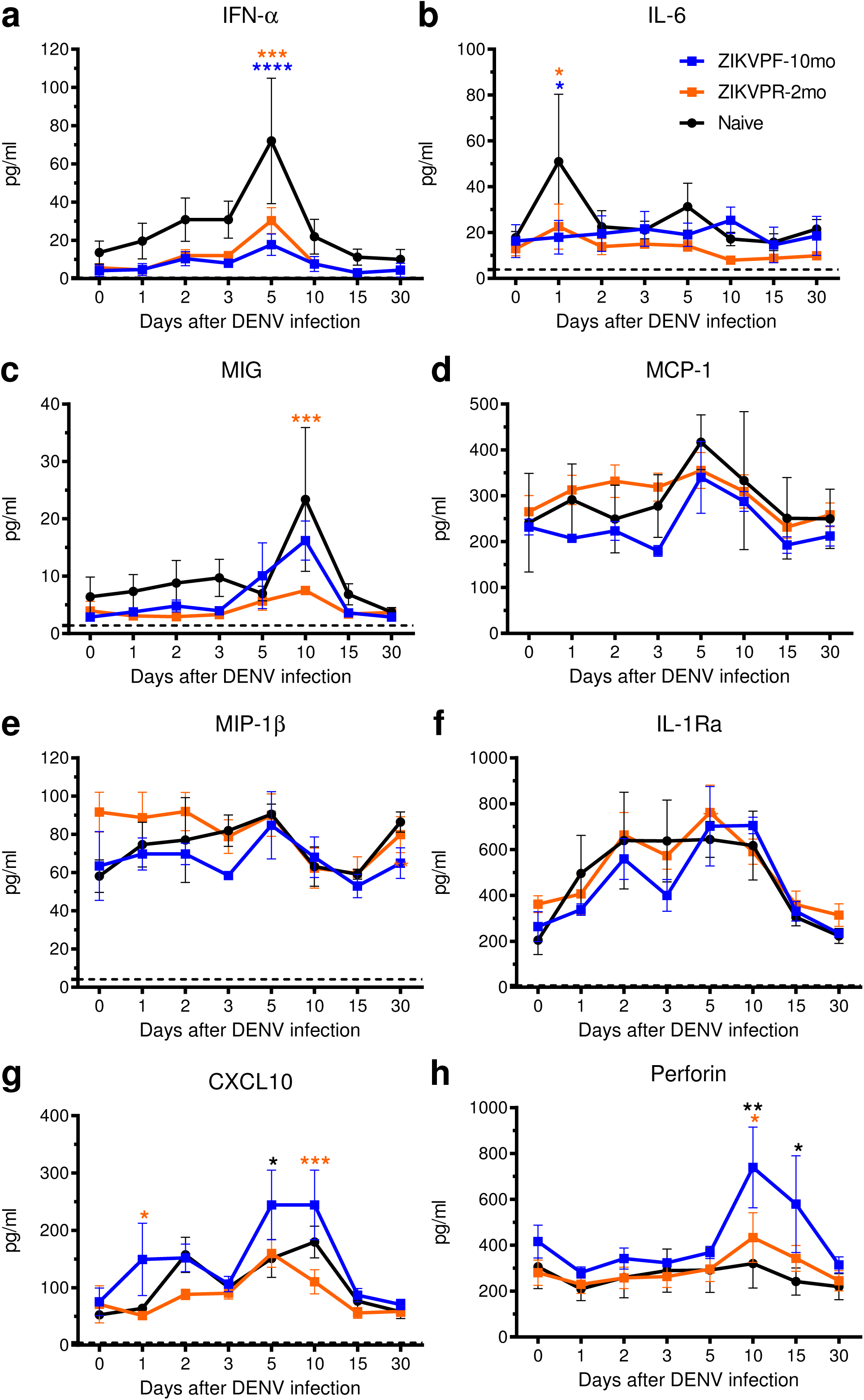
ZIKV immunity does not exacerbate levels of pro-inflammatory cytokines. Cytokines and chemokines expression levels were determined in serum (pg/ml) by multiplex bead assay (Luminex) at baseline, 1, 2, 3, 5, 10, 15 and 30 days after DENV infection. The panel includes: (**a**) interferon alpha (IFN-α), (**b**) interleukin-6 (IL-6), (**c**) monokine induced by IFN-gamma (MIG/CXCL9), (**d**) monocyte chemoattractant protein 1 (MCP-1/CCL2), (**e**) macrophage inflammatory protein 1-beta (MIP-1β/CCL4), (**f**) IL-1 receptor antagonist (IL-1RA), (**g**) C-X-C motif chemokine 10 (CXCL10/IP-10) and (**h**) perforin. Symbols connected with lines represent mean expression levels detected of each cytokine/chemokine per cohort over time: blue squares (ZIKVPF-10mo), orange squares (ZIKVPR-2mo) and black circles (Naïve). Error bars indicate the standard error of the mean (SEM) for each cohort per timepoint. Cutted line mark the limit of detection for each individual cytokine/chemokine. Statistically significant differences between groups were calculated using Two-Way Anova adjusted for Tukey’s multiple comparisons test including 8 families, and 3 comparisons per family. Significant multiplicity adjusted *p* values (* <0.05, ** <0.01, *** <0.001, **** <0.0001) are shown colored representing the cohort against that particular point where is a statistically significant difference between groups. Source data are provided as a Source Data file.

### DENV and ZIKV cross-reactive antibody response

An ELISA-based serological profile was performed to determine the contribution of ZIKV immunity in the cross-reactive Ab response before and after DENV infection. We assessed the levels of DENV IgM and IgG, and cross-reactivity with ZIKV (IgM, IgG, NS1-IgG and EDIII-IgG) at multiple timepoints (Supplementary Fig. 3). Naïve cohort showed a significant higher peak of IgM (Supplementary Fig. 3a) characteristic of a primary DENV infection at 15 and 30 dpi (*p*<0.0001 vs ZIKVPF-10mo and *p*=0.0004 vs ZIKVPR-2mo, *p*=0.0044 vs ZIKVPF-10mo and *p*=0.0179 vs ZIKVPR-2mo, respectively, Two-way Anova Tukey test). This indicates the productive and acute DENV infection, while ZIKV immune groups showed lower levels of IgM resembling a heterotypic secondary infection. Total DENV IgG levels (Supplementary Fig. 3b) of both ZIKV-immune groups were significantly higher compared to naïve since baseline (cross-reactive ZIKV-IgG Abs) and 7, 15, 30, 60 and 90 (the latter for ZIKVPF-10mo only) (ZIKVPF-10mo vs Naïve: *p*=0.0010, *p*<0.0001, *p*<0.0001, *p*<0.0001, *p*<0.0001, *p*=0.0016; ZIKVPF-2mo vs Naïve: *p*=0.0029, *p*=0.0002, *p*<0.0001, *p*<0.0001, *p*=0.0006; Two-way Anova Tukey test). The ZIKVPF-10mo group showed significant higher levels than ZIKVPR-2mo group at 30 and 90 dpi (*p*=0.0242 and *p*=0.0348, Two-way Anova Tukey test). Overall, ZIKVPF-10mo developed higher and long-lasting levels of DENV IgG.

In contrast, ZIKV IgM levels were under or near the limit of detection in all groups over time after DENV infection despite several significant differences between groups (Supplementary Fig. 3c). ZIKV IgG levels (Supplementary Fig. 3d) were high in both ZIKV-immune groups at baseline and 7 dpi compared to naive (*p*<0.0001 vs naïve, Two-way Anova Tukey test), suggesting that although different pre-infecting ZIKV strains, the previous elicited IgG response against both ZIKV strains is comparable. After DENV infection, an increase of ZIKV IgG was shown and remain constantly high at 15, 30, 60 and 90 dpi in both ZIKV-immune groups (*p*<0.0001 vs naïve for all timepoints, Two-way Anova Tukey test), suggesting that DENV has the potential to stimulate ZIKV-binding Ab-producing plasmablasts. In addition, to elucidate the composition of similar ZIKV IgG levels in ZIKV-immune groups, we measured ZIKV-specific NS1 IgG (Supplementary Fig. 3e) and ZIKV-specific EDIII IgG (Supplementary Fig. 3f) levels. Although ZIKVPR-2mo showed significant differences compared to naïve at 30, 60 and 90 dpi (*p*<0.0001, *p*=0.0001, *p*=0.0159; Two-way Anova Tukey test), we observed a significantly higher expansion and long-lasting response of ZIKV NS1-specific Abs in the ZIKVPF-10mo group compared to the ZIKVPR-2mo group at baseline, 60 and 90 dpi (*p*=0.0036, *p*=0.0071, *p*=0.0294; Two-way Anova Tukey test) and also compared to naïve animals at all timepoints (*p*<0.0001, Two-way Anova Tukey test). Moreover, higher magnitude of ZIKV-specific EDIII-IgG levels in the ZIKVPF-10mo group than in the ZIKVPR-2mo group was observed compared to naïve at baseline (ZIKVPF-10mo only), 15, 30 and 60 (ZIKVPF-10mo vs Naïve: *p*=0.0092, *p*<0.0001, *p*<0.0001, *p*=0.0034; ZIKVPR-2mo vs Naïve: *p*=0.0003, *p*=0.0014, *p*=0.0055; Two-way Anova Tukey test), suggesting that a ZIKV mid-convalescence promotes an expansion of higher magnitude of ZIKV EDIII-IgG Abs from ZIKV memory B cells (MBC). However, those higher cross-reacting levels decrease overtime as expected. In summary, a boost of DENV and ZIKV Abs is triggered by the presence of ZIKV immunity and the expansion of specific- and cross-reactive Abs is higher on magnitude and durability when a mid-convalescence immunity to ZIKV is present.

### DENV neutralization is boosted by ZIKV immunity

Neutralizing antibodies (NAbs) are essential to combat DENV and ZIKV infection. The maturation and potency of this response is known to define to a great extent the infection outcome^11,47^. Accordingly, we tested the neutralization capacity of NAbs in serum from ZIKV-immune and naïve animals before and after DENV infection, to determine whether an early- or mid-convalescence to ZIKV affected the NAb response. Plaque Reduction Neutralization Test (PRNT) was performed to elucidate the NAb titers of all groups against all DENV serotypes and both ZIKV pre-infecting strains. Before infection with DENV the naïve groups had no detectable NAb levels (<1:20 PRNT60 titers) against all DENV serotypes, while ZIKV-immune groups showed low cross-NAb titers against DENV-2 and DENV-4 (Fig. 4a). These cross-reactive levels were higher in the ZIKVPF-10mo group than in the ZIKVPR-2mo group for both viruses. The peak of high NAb titers occurred at 30 days after DENV infection for all groups (ZIKVPF-10mo>ZIKVPR-2mo>Naïve) against all DENV serotypes (DENV-2>DENV-4>DENV-3>DENV-1) (Fig. 4b). The ZIKVPF-10mo group neutralized all DENV serotypes with significant higher potency than naïve animals (*p*<0.0001, *p*=0.0337, *p*<0.0001, *p*<0.0001 for DENV1-4; Two-way Anova Tukey test) and the ZIKVPR-2mo group, except for DENV-2, that both ZIKV-immune groups have comparable neutralization magnitude at 30 dpi (*p*=0.0002, *p*=0.7636, *p*=0.0016, *p*=0.0004 for DENV1-4; Two-way Anova Tukey test). However, the neutralization kinetics by sigmoidal response curves suggest higher percent of neutralization against DENV-2 overtime in the group with mid-convalescence to ZIKV (Supplementary Fig. 4). On the other hand, the ZIKVPR-2mo group showed significantly higher potency of the NAb response only against DENV-1 compared to naive animals (*p*=0.0146; Two-way Anova Tukey test) (Fig. 4b).

**Figure 4.**
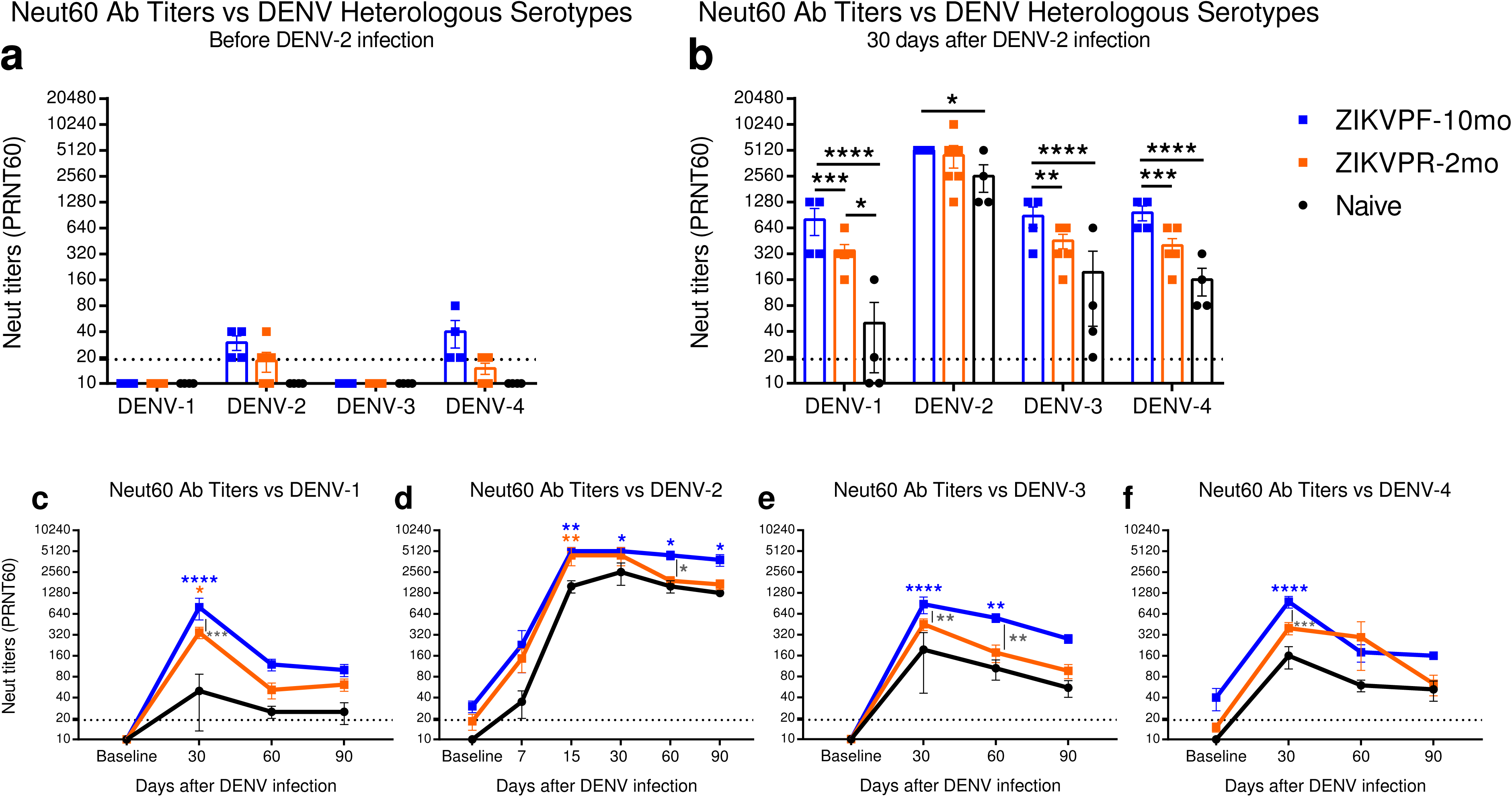
Neutralization of DENV serotypes by ZIKV-immune animals is higher in magnitude. The magnitude of the neutralizing antibody (NAb) response was determined (**a**) before and (**b**) 30 days after DENV infection by Plaque Reduction Neutralization Test (PRNT) against all DENV serotypes. (**c-f**) The durability of the neutralizing response was assessed measuring NAb titers up to 90 dpi against all DENV serotypes. Symbols connected with full lines indicate mean levels of NAb titers detected per cohort over time: blue squares (ZIKVPF-10mo), orange squares (ZIKVPR-2mo) and black circles (Naïve). Error bars represent the standard error of the mean (SEM). PRNT60: NAb titer capable of reduce 60% or more of DENV serotypes plaque-forming units (pfu) compared with the mock (control of virus without serum). A PRNT60 1:20 titer was considered positive, and <1:20 as a negative Neut titer. Dotted line mark <1:20 for negative results. Non-neutralizing titers (<1:20) were assigned with one-half of the limit of detection for graphs visualization (1:10). Statistically significant differences between groups were calculated using Two-Way Anova adjusted for Tukey’s multiple comparisons test including 4 and 6 families for heterologous serotypes and DENV-2, respectively, and 3 comparisons per family. Significant multiplicity adjusted *p* values (* <0.05, ** <0.01, *** <0.001, **** <0.0001) are shown. Blue and orange asterisks represent significant difference between the corresponded ZIKV immune groups and naive group, and gray asterisks indicate a significant difference between ZIKV immune groups. Source data are provided as a Source Data file.

In addition, we tested whether the NAb titers that peak at 30 dpi for all groups remain constant over time (up to 90 dpi) against all DENV serotypes (Fig. 4c-f). In general, the neutralizing response of the ZIKVPF-10mo maintained higher NAb titers up to 90 dpi compared to ZIKVPR-2mo and naïve groups. Significant differences between ZIKVPF-10mo and ZIKVPR-2mo groups were observed against DENV-1,-3 and -4 at day 30 pi (*p*=0.0002, *p*=0.0016, *p*=0.0004; Two-way Anova Tukey test) and at day 60 pi against DENV-2 and DENV-3 (*p*=0.0179, *p*=0.0047; Two-way Anova Tukey test). The neutralizing Ab response of the ZIKVPF-10mo group was even more significantly higher compared to the naïve group at day 15 (only performed for the infecting serotype to monitor early neutralizing activity), day 30, 60 and 90 pi against DENV-2 (*p*=0.0022, *p*=0.0337, *p*=0.0146, *p*=0.0337; Two-way Anova Tukey test); at day 30 pi against DENV-1 (*p<*0.0001, Two-way Anova Tukey test); at day 30 and 60 pi against DENV-3 (*p<*0.0001, Two-way Anova Tukey test); and at day 30 pi against DENV-4 (*p<*0.0001, Two-way Anova Tukey test). In contrast, the ZIKVPR-2mo group showed a neutralizing Ab response with a magnitude and long-lasting levels comparable to the naïve group, except at day 15 and 30 pi against DENV-2 and DENV-1, respectively (*p*=0.0067, *p*=0.0146; Two-way Anova Tukey test). The neutralizing response was long-lasting in the ZIKVPF-10mo group compared to other groups as supported by the data from days 30 and 60 p.i. At day 90 pi, although no significant differences were observed between all groups, the ZIKVPF-10mo group showed a consistent trend to maintain higher NAb titers against all DENV serotypes indicating a higher and long-lasting breadth of cross-neutralization within DENV serocomplex.

Collectively, these results demonstrate that a mid-convalescence to ZIKV provokes a boost of the magnitude and durability of the neutralizing response against all DENV serotypes more effectively than in animals with an early-convalescence to ZIKV, but also higher compared to a *de novo* DENV-specific NAb response of the naïve animals.

### ZIKV neutralization is boosted by DENV infection

Previous exposure to ZIKV strains in ZIKV-immune groups developed high levels of cross-reactive, non-neutralizing, and neutralizing Abs before DENV infection (baseline). To determine if this memory Ab response is strain-specific and if the difference in convalescence period to ZIKV alters the efficacy and modulation after DENV infection, we assessed the NAb levels in ZIKV-immune (ZIKVPF-10mo and ZIKVPR-2mo) and ZIKV-naïve serum with both pre-infecting contemporary Asian-lineage H/PF/2013 and PRVABC59 ZIKV strains at multiple timepoints after DENV infection. At baseline, both ZIKV-immune groups showed high NAb titers against H/PF/2013 strain, which suggest that irrespective of pre-exposure to different ZIKV strains and different convalescent periods the Ab response remains similarly effective (Fig. 5a). As early as day 15 after DENV infection, a potent boost of NAb titers in both ZIKV-immune groups was developed. However, elevated NAb titers were significantly higher in the ZIKVPF-10mo group compared to the ZIKVPR-2mo and naïve groups at day 15 pi (*p*=0.0005, *p*<0.0001; Two-way Anova Tukey test) and day 30 pi (*p*=0.0067, *p*=0.0012; Two-way Anova Tukey test). As expected, this elevated ZIKV cross-reactive NAb levels decreased gradually overtime after 15 dpi in both ZIKV-immune groups. Nevertheless, the ZIKVPF-10mo group retained higher NAb titers until 90 dpi while the titers of the ZIKVPR-2mo group returned to baseline levels. Of note, the NAb titers of the naïve group were considered as negative in all timepoints and failed to neutralize ZIKV throughout DENV infection even at concentrated levels of the antibodies (Fig. 5a). These results are confirmed by the behavior of neutralization kinetics by sigmoidal response curves where the ZIKVPF-10mo group retained elevated magnitude of ZIKV neutralization overtime (Supplementary Fig. 5).

**Figure 5.**
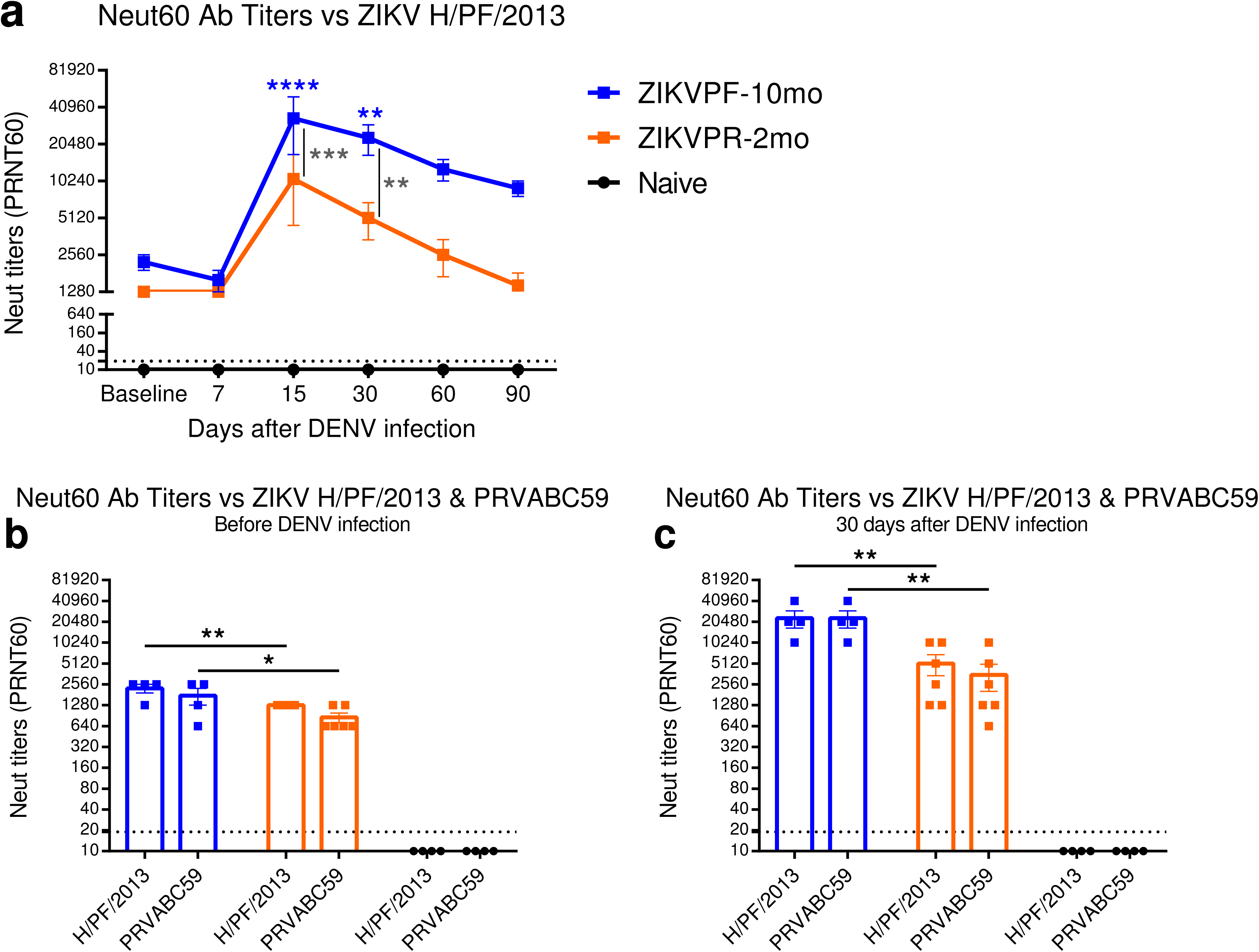
ZIKV neutralization is boosted after DENV infection and is strain independent. (**a**) NAb titers against ZIKV H/PF/2013 were determined by PRNT60 at baseline, 7, 15, 30, 60 and 90 days after DENV infection. Comparison of NAb titers between pre-infecting ZIKV strains was performed (**b**) before and (**c**) after DENV infection. Symbols connected with full lines indicate mean levels of NAb titers detected per cohort over time: blue squares (ZIKVPF-10mo), orange squares (ZIKVPR-2mo) and black circles (Naïve). Error bars represent the standard error of the mean (SEM). PRNT60: NAb titer capable of reduce 60% or more of ZIKV strains plaque-forming units (pfu) compared with the mock (control of virus without serum). A PRNT60 1:20 titer was considered positive, and <1:20 as a negative Neut titer. Dotted line mark <1:20 for negative results. Non-neutralizing titers (<1:20) were assigned with one-half of the limit of detection for graphs visualization (1:10). Statistically significant differences between groups were calculated using Two-Way Anova adjusted for Tukey’s multiple comparisons test including 6 and 2 families for panel a and b-c, respectively, and 3 comparisons per family. Significant multiplicity adjusted *p* values (* <0.05, ** <0.01, *** <0.001, **** <0.0001) are shown. Blue and orange asterisks represent significant difference between the corresponded ZIKV-immune groups and naive group, and gray asterisks indicate a significant difference between ZIKV-immune groups. Source data are provided as a Source Data file.

To determine if the immune memory induced by different ZIKV strains play a role in the modulation of the cross-NAb response triggered by a subsequent DENV infection, NAb titers were measured against both ZIKV strains before and 30 days after DENV infection. The ZIKVPF-10mo group showed significant higher NAb titers against both ZIKV strains compared to the ZIKVPR-2mo group before DENV infection (*p*=0.0093, *p*=0.0141; Two-way Anova Tukey test) (Fig. 5b). Subsequently, DENV infection promote an equally 8-fold increase of NAb titers against both strains in the ZIKVPF-10mo group, significantly higher than the 4-fold increase in the ZIKVPR-2mo group (*p*=0.0025, *p*=0.0011; Two-way Anova Tukey test) (Fig. 5c). To rule out that difference in fitness between both ZIKV strains would bias the magnitude of the NAbs after DENV infection we compared in parallel the NAb titers at 30 and 60 days after ZIKV infection (day 60 correspond to the baseline of the ZIKVPR-2mo group). No significant differences were observed between ZIKV-immune groups in the NAb titers induced by both strains at the same timepoints after ZIKV infection (Supplementary Fig. 6). Altogether, these results demonstrate that DENV infection results in a significant increase in the magnitude and durability of the cross-neutralizing Ab response against ZIKV in animals with a mid-convalescent period from ZIKV infection. The elicited changes in neutralization capacity were likely driven more by the longevity of the immune memory maturation and the associated memory recall of the ZIKV immunity than by a strict dependency of the specific pre-exposed ZIKV strain.

### Immune cell subsets modulated by ZIKV immunity

We performed immunophenotyping by flow cytometry to assess the frequency, early activation and proliferation of multiple immune cell subsets and how these parameters are affected by the presence of pre-existing immunity to ZIKV on a subsequent DENV infection (Supplementary Fig. 7, 8, and 9 for gating strategy; Supplementary Table 2 for Ab panel). As part of the innate immune response, the frequency of dendritic cells (DCs) and natural killer (NK) cells subpopulations were measured. Plasmacytoid DCs (pDCs: Lin^-^HLA-DR^+^CD123^+^) are known to respond to viral infection by production of IFN-α, while myeloid DCs (mDCs: Lin^-^HLA-DR^+^CD11c^+^) interacts with T cells. The frequency of pDCs was not significantly altered by DENV infection in any group compared to baseline levels (Supplementary Fig. 10a). At day 2 pi we detected a significant increase of mDCs in the ZIKVPF-10mo group (*p*=0.0082; Two-way Anova Dunnett test) (Supplementary Fig. 10b). Furthermore, we determined the frequency of NK subpopulations including: NKCD8, NKCD56, NKp30 and NKp46 (Supplementary Fig. 11). In general, no differences were detected between baseline and after DENV infection in all groups for all NK subpopulations and receptor expression with the exception of the ZIKVPR-2mo group that showed a significant increases in the following subpopulations: NKG2A^+^NKp30 and NKp30^+^NKp46^+^ at 7 dpi (*p*=0.0495, *p*=0.0006; Two-way Anova Dunnett test) and NKp46^+^NKp30^+^ at 7 and 10 dpi (*p*=0.0005, *p*=0.0001; Two-way Anova Dunnett test) (Supplementary Fig. 11j, o, s).

We next investigated cell subsets that are part of the bi-phasic (humoral/cellular) adaptive immune response such as B (CD20+CD3-) and T (CD3+CD20-) cells. No differences were detected in total B cells between groups following DENV infection compared to baseline levels (Supplementary Fig. 12a), but ZIKV-immune groups had elevated levels of activated B cells (CD20+CD3-CD69+) since baseline and a trend to increase these levels more than the naïve group overtime (Supplementary Fig. 12b). We detected a significant decrease of proliferating B cells (CD20+CD3-Ki67+) in naïve animals at 7 and 10 dpi (p=0.0031, p=0.0345; Two-way Anova Dunnett test), while ZIKV-immune groups retained their proliferating levels (Supplementary Fig. 12c). Interestingly, the ZIKVPF-10mo group showed a significant increase of B cells that were proliferating and activated simultaneously (CD20+CD3-CD69+Ki67+) as early as in day 1 pi (p=0.0240; Two-way Anova Dunnett test) and maintained higher levels up to 10 dpi (Supplementary Fig. 12d). Together, these phenotyping results of B cells are consistent with the early and boosted production of binding and neutralizing Abs in the ZIKVPF-10mo group compared to naïve animals. The frequency of total T cells (CD3^+^CD20^-^) and CD4^+^/CD8^+^ T cells subsets, was comparable at all timepoints before and after DENV infection in all groups of animals (Supplementary Fig. 13a-c).

Previous studies have demonstrated that DENV and ZIKV specific CD4^+^ and CD8^+^ T cells are enriched in certain memory subsets^23,48^. Thus, we measured whether the early activation of T cell subpopulations, such as effector memory (CD3^+^CD4^+^CD28^-^CD95^+^) and central memory (CD3^+^CD4^+^CD28^+^CD95^+^) T cells (T-EM and T-CM), within each T cell compartment was modulated following DENV infection in presence or absence of convalescence to ZIKV (Fig. 6, and Supplementary Fig. 7 for gating strategy). The ZIKVPF-10mo group showed significant higher frequency of activated CD4^+^ and CD8^+^ T-EM (CD3^+^CD4^+^CD28^-^CD95^+^CD69^+^ and CD3^+^CD8^+^CD28^-^CD95^+^CD69^+^) following DENV infection compared to basal levels (CD4^+^ T-EM at 7 and 10 dpi: *p*=0.0001, *p*=0.0072; CD8^+^ T-EM at 2 and 7 dpi: *p*=0.0291, *p*=0.0001; Two-way Anova Dunnett test) (Fig. 6a, d). Interestingly, the ZIKVPR-2mo group showed a very limited frequency and activation of the CD4^+^ and CD8^+^ T-EM compared to the ZIKVPF-10mo and naïve groups. However, this group with an early convalescent period to ZIKV, contrary to the other two groups, showed a very limited but significant activation of CD8^+^ T-CM (CD3^+^CD8^+^CD28^+^CD95^+^CD69^+^) at day 7 and 10 pi (*p*=0.0007, *p*=0.0147; Two-way Anova Dunnett test) (Fig. 6e). In contrast, naïve animals did not show any significant activation of these memory cell subsets after DENV infection. Collectively, these results suggest that following DENV infection: (i) animals with a mid-convalescence ZIKV immunity have a more dynamic B cell response and are able to rapidly produce more activated effector memory T cells from both T cell compartments; (ii) animals with an early-convalescence to ZIKV induced activation of central memory T cells in the CD8^+^ compartment with a very limited T-EM frequency and activation profile compatible with a contraction phase of the T cells compartments; (iii) and animals without previous exposure to ZIKV exhibited a limited B cell response and minimal modulation of memory T cell subpopulations at early timepoints as the ZIKV-immune groups.

**Figure 6.**
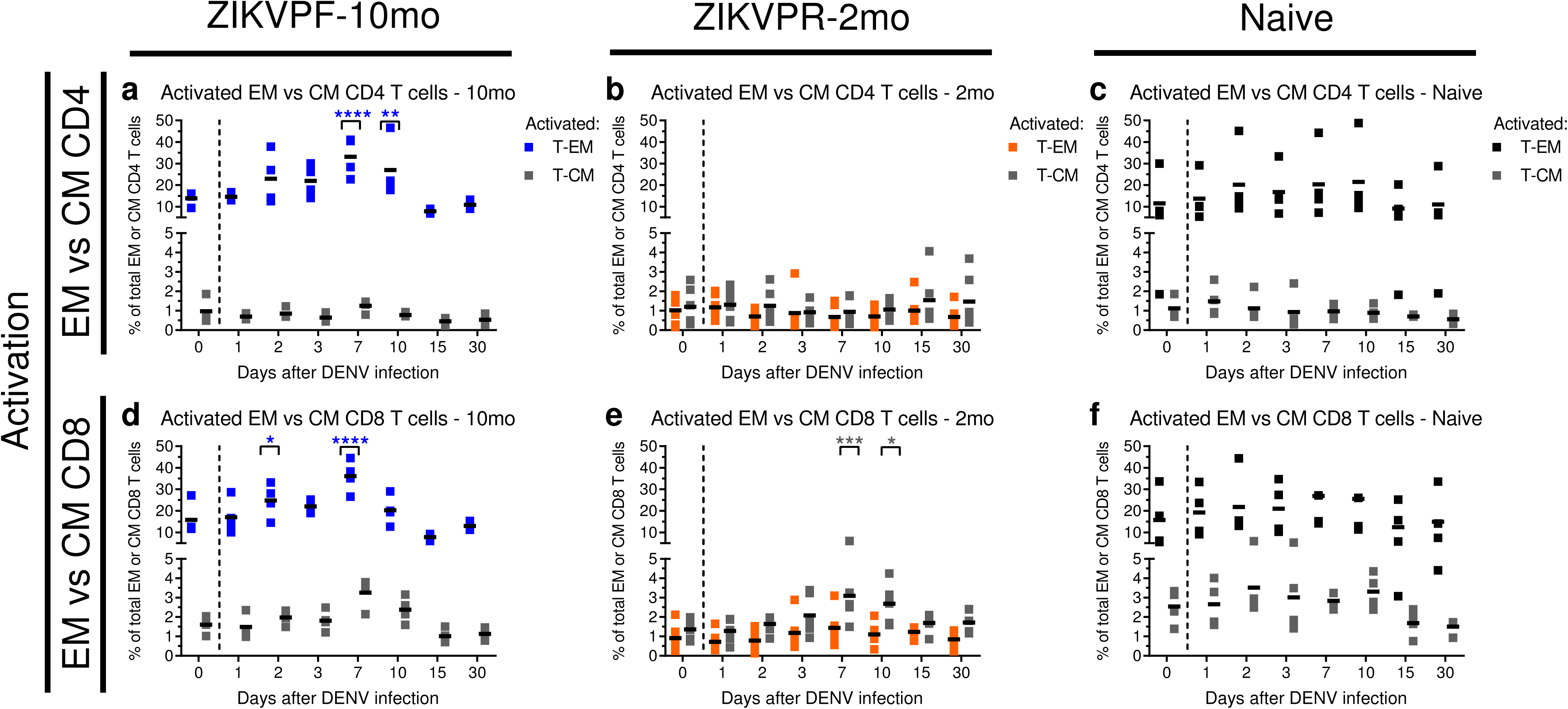
Activation of effector and central memory CD4^+^ and CD8^+^ T cells after DENV infection. Activation (CD69^+^) of effector memory (T-EM: CD3^+^CD4^+^CD28^-^CD95^+^) and central memory (T-CM: CD3^+^CD4^+^CD28^+^CD95^+^) T cells within (**a-c**) CD4^+^ and (**d-f**) CD8^+^ T cell compartments before and after DENV infection. Percent of cells were determined by immunophenotyping using flow cytometry (Supplementary Fig. 7 for gating strategy). Blue, orange and black squares represent T-EM for ZIKVPF-10mo, ZIKVPR-2mo and Naïve, respectively. Gray squares represent T-CM for each group. Short black lines mark mean value for each group per timepoint. Cutted line divide % of T-EM and T-CM cells quantified before and after DENV infection. Statistically significant differences within groups were determined using Two-Way Anova adjusted for Dunnett’s multiple comparisons test (comparison of each group response at each timepoint versus baseline of the same group) including 2 families, and 7 comparisons per family. Significant differences are reported as multiplicity adjusted *p* values (* <0.05, ** <0.01, *** <0.001, **** <0.0001). Asterisks represent significant difference between the corresponded timepoint and baseline within the same group. Source data are provided as a Source Data file.

### T cell functional response is shaped by ZIKV immunity

To further characterize the cross-reactive T cell response, we investigated if different convalescent periods of ZIKV immunity impacted the outcome of the effector role of CD4^+^ and CD8^+^ T cells following DENV infection. PBMCs were isolated and stimulated with peptide pools from DENV and ZIKV envelope (E) proteins and from ZIKV non-structural proteins (ZIKV-NS) (Supplementary Table 4 for peptide sequences). Then, intracellular cytokine staining using flow cytometry analysis (Supplementary Fig. 14 for gating strategy; Supplementary Table 3 for Ab panel) was performed to quantify the production of effector immune markers such as the cytotoxic marker CD107a, IFN-γ, and TNF-α by CD4^+^ and CD8^+^ T cells at baseline, 30, 60, and 90 days after DENV infection (Fig. 7).

**Figure 7.**
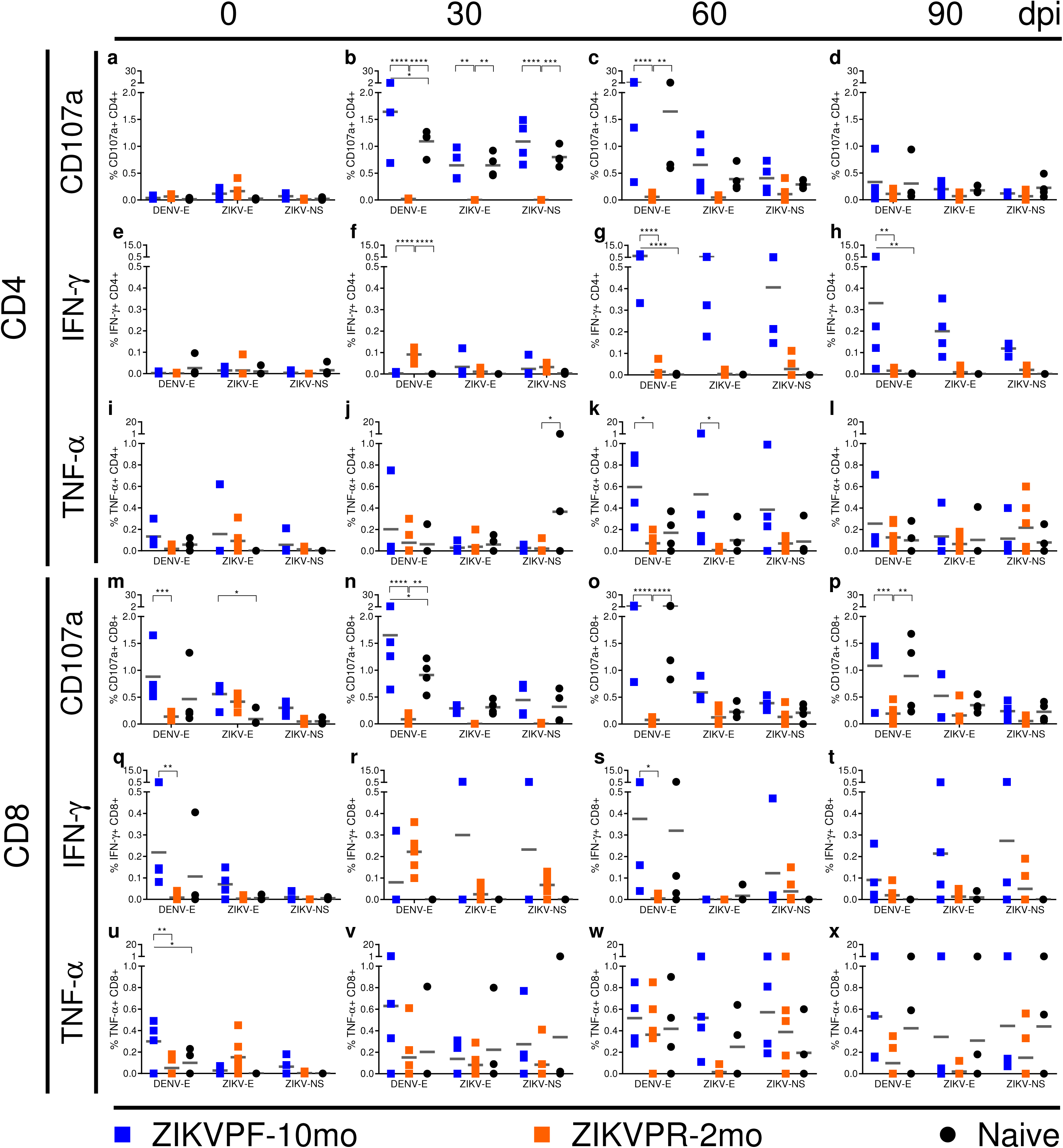
Longevity of ZIKV immunity shapes the T cell functional response. T cell functional effector response was determined by the quantification (%) of (**a-d; m-p**) CD107a-expressing and (**e-h; q-t**) IFN-γ or (**i-l; u-x**) TNF-α producing CD4^+^ and CD8^+^ T cells before (0) and 30, 60 and 90 days after DENV infection. Responses to several peptide pools that encode for DENV and ZIKV envelope (E) proteins or ZIKV non-structural (NS) protein were quantified. After antigenic stimulation intracellular cytokine staining was performed using flow cytometry analysis (Supplementary Fig. 14 for gating strategy). Individual symbols represent each animal per antigenic stimulation over time: blue squares (ZIKVPF-10mo), orange squares (ZIKVPR-2mo) and black circles (Naïve). Short gray lines mark mean value for each group. Statistically significant differences between groups were calculated using Two-Way Anova adjusted for Tukey’s multiple comparisons test including 3 families, and 3 comparisons per family. Significant multiplicity adjusted *p* values (* <0.05, ** <0.01, *** <0.001, **** <0.0001) are shown. Asterisks represent significant difference between indicated groups. Source data are provided as a Source Data file.

To assess the ZIKV-primed specific- or cross-reactive effector T cell response we studied the response against ZIKV or DENV stimuli before DENV infection. In general, before DENV infection, we found that the ZIKV-primed effector T cell response was higher in CD8^+^ (Fig. 7m, q, u) than in CD4^+^ (Fig 7a, e, i) T cells. Of note, significant higher levels of CD107a, INF-γ and TNF-α producing CD8^+^ T cells were found only in the ZIKVPF-10mo group before DENV infection (ZIKVPF-10mo vs ZIKVPR-2mo for CD107a: *p*=0.0002; ZIKVPF-10mo vs Naïve for CD107a: *p*=0.0401; ZIKVPF-10mo vs ZIKVPR-2mo for INF-γ: *p*=0.0020; ZIKVPF-10mo vs ZIKVPR-2mo for TNF-α: *p*=0.0033; ZIKVPF-10mo vs Naïve for TNF-α: *p*=0.0354; Two-way Anova Tukey test) (Fig. 7m, q, u). This basal effector response of CD8^+^ T cells in the ZIKVPF-10mo group is predominated by cross-reactive CD8^+^ T cells against DENV E protein. Very low effector T cell response against ZIKV NS proteins was detected for all groups (ZIKVPF-10mo>ZIKVPR-2mo>Naïve). In summary, results of T cell functional response before DENV infection suggest that a mid-convalescence to ZIKV provoke a higher CD8^+^ T cell effector response capable to cross-react efficiently with DENV E protein.

After DENV infection, we were able to determine the modulation of the ZIKV-primed effector CD4^+^ and CD8^+^ T cell responses of ZIKV-immune groups and the *de novo* response of ZIKV-naïve animals. The ZIKVPF-10mo and naïve groups significantly boosted their CD107a expression in both T cell compartments stimulated mainly by DENV E protein at 30 and up to 90 days after DENV infection (CD4^+^ T cells: ZIKVPF-10mo vs ZIKVPR-2mo: *p*<0.0001 at 30 dpi, *p*<0.0001 at 60 dpi; Naïve vs ZIKVPR-2mo: *p*<0.0001 at 30 dpi, *p*=0.0018 at 60 dpi; ZIKVPF-10mo vs Naïve: *p*=0.0204 at 30 dpi. CD8^+^ T cells: ZIKVPF-10mo vs ZIKVPR-2mo: *p*<0.0001 at 30 dpi, *p*<0.0001 at 60 dpi, *p*=0.0008 at 90 dpi; Naïve vs ZIKVPR-2mo: *p*=0.0039 at 30 dpi, *p*<0.0001 at 60 dpi; *p*=0.0081 at 90 dpi; ZIKVPF-10mo vs Naïve: *p*=0.0194 at 30 dpi; Two-way Anova Tukey test) (Fig. 7b, c, n, o, p). Also, these groups boosted the CD107a cytotoxic signature reacting against ZIKV E and NS proteins by cross-reactive CD4^+^ T cells 30 days after DENV infection (ZIKVPF-10mo vs ZIKVPR-2mo: *p*=0.0025 for ZIKV E, *p*<0.0001 for ZIKV NS; Naïve vs ZIKVPR-2mo: *p*=0.0025 for ZIKV E, *p*=0.0002 for ZIKV NS; Two-way Anova Tukey test) (Fig. 7b).

The ZIKVPF-10mo group showed a remarkable significant increase of the IFN-γ-producing CD4^+^ T cells against DENV E protein since 60 dpi and is maintained up to 90 dpi compared to other groups (ZIKVPF-10mo vs ZIKVPR-2mo at 60 and 90 dpi: *p*<0.0001, *p*=0.0024; ZIKVPF-10mo vs Naïve at 60 and 90 dpi: *p*<0.0001, *p*=0.0037; Two-way Anova Tukey test) (Fig. 7g, h), and was the only group with significant increase in the IFN-γ producing CD8^+^ T cell compartment at 60 dpi (ZIKVPF-10mo vs ZIKVPR-2mo: *p*=0.0253; Two-way Anova Tukey test) (Fig. 7s). On the other hand, the ZIKVPR-2mo group exhibited a significant increase of IFN-γ producing CD4^+^ T cells earlier than other groups at 30 dpi (ZIKVPR-2mo vs ZIKVPF-10mo: *p*<0.0001; ZIKVPR-2mo vs Naïve: Two-way Anova Tukey test) (Fig. 7f). Interestingly, the naïve group showed an increase of cross-reactive TNF-α producing CD4^+^ T cells against ZIKV NS proteins 30 days after DENV infection (Naïve vs ZIKVPR-2mo: *p*=0.0359; Two-way Anova Tukey test) (Fig. 7j). The ZIKVPF-10mo group developed a significant effector T cell response by TNF-α producing CD4^+^ T cells against DENV and ZIKV E proteins at 60 days after DENV infection (ZIKVPF-10mo vs ZIKVPR-2mo against DENV/ZIKV E protein: *p*=0.0163, *p*=0.0172; Two-way Anova Tukey test) (Fig. 7k). Although all groups showed a boosted TNF-α effector response in the CD8^+^ T cell compartment up to 90 days after DENV infection, no significant differences between groups were observed.

Collectively, these results after DENV infection suggest that a mid-convalescence to ZIKV translate in a more complete functional T cell response characterized by: (i) a cytotoxic CD107a^+^ phenotype directed to DENV E protein for both T cell compartments comparable to the DENV-specific *de novo* response of the naïve group, (ii) developed CD107a, IFN-γ and TNF-α producing CD8^+^ T cell effector response that cross-react efficiently with DENV E protein since baseline and is boosted after DENV infection, (iii) and promoted the higher T cell effector response against ZIKV NS proteins. An early-convalescence to ZIKV results in (iv) a very limited cytotoxic activity (limited expression of CD107a marker) which is in line with a very limited activation of the T-EM, and with failed capability to react efficiently against E or NS proteins. The ZIKV-naïve group response was characterized by: (v) production of a DENV-specific *de novo* functional T cell response with similar magnitude between both T cell compartments, (vi) capable to cross-react against ZIKV E and NS proteins, (vii) and able to mount a DENV-specific cytotoxic CD107a^+^ phenotype.

## Discussion

We found that previous ZIKV infection modulates the immune response against subsequent DENV infection without an enhancement of DENV viremia nor pro-inflammatory status. This modulation is influenced by the longevity of ZIKV convalescence—more after longer ZIKV pre-exposure.

The aftermath of the recent ZIKV epidemic has been related to a remarkable decrease in DENV cases in Brazil ^27^, and also in most of Latin American and Caribbean countries (http://www.paho.org/data/index.php/es/temas/indicadores-dengue/dengue-nacional/9-dengue-pais-ano.html?start=2)^24^. Yet, little is known about the role of previous ZIKV immunity in the outcome of a subsequent DENV infection in human populations, and if ZIKV immunity is supporting this epidemiological phenomenon observed post-ZIKV epidemic^27^. To evaluate the hypothesis of a potential ZIKV-DENV cross-protection in humans characterizing the immunological history of prospective cohorts^45^ will be necessary, but human samples for this purpose are scarce yet. Because of this, NHPs are key to provide knowledge and anticipate different immunological scenarios when DENV epidemics re-emerge in human populations with previous immunity to ZIKV.

Animals with pre-existing ZIKV immunity do not show an enhancement of DENV-induced RNAemia, regardless of the period of convalescence from previous ZIKV infection (10 or 2 months) and different pre-infecting ZIKV strains. Previous ZIKV immunity is associated with a trend of less RNAemia days during subsequent DENV infection. This effect is more evident in animals with a ZIKV convalescence period of 10 months. Previous work reported that a period of early-convalescence (56 days) to ZIKV (PRVABC59 strain) in rhesus macaques was associated with a significant increase of DENV-2 RNAemia at day 5 after DENV infection and a pro-inflammatory cytokine profile. However, very similar to our results, it was noteworthy a delay at early timepoints and an early clearance in late timepoints of the DENV-2 RNAemia in ZIKV-immune macaques in comparison to the naïve ones^37^. The lack of significant DENV RNAemia enhancement found in the group with the early-convalescence period in our work, compared to previous results^37^, may be attributable to the different sample types collected (plasma vs serum), or different DENV-2 strains used for the challenge [New Guinea/1944 strain vs Thailand/16681/1964 strain, from Asian II and Asian I Genotype, respectively]. This fact is of relevance because it suggests that the effect of previous ZIKV immunity on a subsequent DENV infection may differ between DENV serotypes or even within genotypes. Another possible explanation is the genetic heterogeneity of rhesus macaques used in these two studies as they are derived from different breeders. The importance of selecting genetic well-characterized macaques have been discussed previously^49^.

Due to limited availability of ZIKV-immune cohorts we used animals infected with two different ZIKV strains for our subsequent challenge with DENV-2. However, extensive revision of the literature up to date reveals a broad consensus that these two contemporary ZIKV strains behave very similar from an antigenic point of view^11,50–52^. Our results are confirmatory of those results showing that both ZIKV strains were neutralized with same efficacy by serum within each ZIKV-convalescent group, explained by the broadly neutralization activity against multiple ZIKV strains irrespective of the infecting strain^51^. However, the magnitude of the neutralization of both strains was statistically higher in animals exposed to DENV 10 months (mid-convalescence) after ZIKV infection compared to the animals with a shorter ZIKV convalescence (2 months). These results suggest that the differences in the neutralization profile between the two ZIKV-immune groups are associated to the longevity of ZIKV convalescence which may be attributable to the maturation of the cross-reactive immune memory elicited by the heterologous DENV infection and no to the antigenic differences or the different replication capabilities in rhesus macaques of those two pre-infecting ZIKV strains^16,53^.

The period of convalescence further had an impact in the maintenance of the neutralization magnitude against ZIKV and DENV overtime. We observed a higher activation of the memory immune response characterized by transiently higher peak levels of serum NAbs against DENVs and ZIKV in ZIKVPF-10mo immune animals compared to ZIKVPR-2mo immune animals challenged with DENV-2. However, unlike heterologous infections with different DENV serotypes, by 90 days after DENV-2 infection, the naïve and ZIKV-immune animals had similar levels of DENV-2 NAbs. On the other hand, ZIKV NAbs decay overtime in ZIKV-immune animals after DENV infection, but animals with longer convalescence retain higher titers until the end of the study. Overall, these results demonstrate that pre-existing ZIKV immunity leads to a transient increase in neutralizing Ab responses in animals challenged with DENV-2 compared to naïve animals. This is in contrast with previous findings were ZIKV-convalescent macaques show a lack of an early and delayed anamnestic response overtime with limited induction of DENV NAbs compared to ZIKV-naïve animals after DENV infection^54^. However, our results show the ability of DENV-2 to activate MBCs stimulated by the previous ZIKV infection, but this activation is modest and short-lived compared to the robust and sustained activation of MBCs on secondary DENV infections ^10,30,46,55^. Is still uncertain why the ZIKVPF-10mo animals have a slightly higher peak of Ab response compared to the ZIKVPR-2mo animals. We speculate this may be caused by modification of MBCs overtime, so that by 10 months the cells are able to better respond to antigen compared to cells at two months. After ZIKV infection in human DENV-naïve subjects, the ZIKV/DENV cross-reactive MBC response increased in magnitude (39% of total MBC proportion) after longer periods of ZIKV convalescence (∼8 months post-ZIKV infection)^56^, similar to the 10 months in the ZIKV mid-convalescent group that exhibited higher DENV cross-neutralization. Based upon studies of human monoclonal Abs, plasmablasts response during secondary DENV infection is mainly of MBC origin, resulting in a mature response characterized by cross-neutralizing Abs *in vitro*^57^. These are seminal contributions to forecast and understand the cross-neutralization capacity of further heterologous DENV epidemics in the context of previous ZIKV-DENV immunity. Interestingly, ZIKV-convalescent animals showed some degree of cross-neutralization against DENV-2 and DENV-4 before DENV infection. This is consistent with our previous results showing that DENV-naïve ZIKV-infected animals also preferentially neutralized DENV-4 followed by DENV-2 after ZIKV infection^16^. Longitudinal data of cross-neutralization of DENV serotypes in DENV-naïve ZIKV-infected human subjects showed low cross-neutralization against all DENV serotypes, but DENV-4 followed by DENV-2 were neutralized more efficiently up to 6 months after ZIKV infection with comparable basal titers reported here^58^. There is no data yet that delineates shared cross-neutralizing epitopes between ZIKV and DENV-2/-4, but it is known that DENV-4 genotypic diversity impact the capacity of its neutralization^59^.

Early studies of T cells associate their contribution towards immunopathogenesis in DENV secondary infections explained by the original antigenic sin^60^, but increasing evidence suggest their protective role during primary and secondary DENV infections^61^. Recently, with the introduction of ZIKV into The Americas, T cells from DENV immunity are being implicated in mediating cross-protection against ZIKV^21–23^. We found that animals with a mid-convalescence to ZIKV developed an early activation of CD4^+^ and CD8^+^ effector memory T cells after DENV infection. This early activation has been observed for the opposite scenario in DENV-immune ZIKV-infected patients^23^. Interestingly, the ZIKV early-convalescent group displays a modest activation (T-CM>T-EM) early after DENV infection. Since this group was infected with ZIKV only two months before DENV it is possible that after viral clearance and development of ZIKV-specific T cell response, the T cell compartments were still under the contraction phase at the time of the DENV challenge. Yellow fever virus (YFV) and vaccinia virus vaccinations in humans demonstrate that T cell contraction start as early as approximately one-month post-vaccination and at least for almost three months is still ongoing^62^. Also, a study shows that re-stimulation using alphavirus replicons during T cell response contraction does not have significant impact modulating the pre-existing T cell response^63^.

The profile of ZIKV-specific CD8^+^ T cells in humans with convalescence to ZIKV is characterized by the production of IFN-γ, and expression of activation and cytotoxic markers^64^. Presence of sustained levels of IFN-γ prior and early after DENV challenge in vaccinees has been associated with protection against viremia and/or severe disease^65,66^. We observed a similar phenotype of the functional response of CD8^+^ T cells prior DENV infection in animals with longer convalescence to ZIKV. Strikingly, this response recognizes more efficiently peptides from DENV E protein than from ZIKV E protein. However, ZIKV-specific CD8^+^ T cells direct 57% of their response against structural proteins, which may suggest these cells can recognize conserved epitopes between ZIKV and DENV structural proteins. Cross-reactivity of T cells between heterologous flavivirus infections is explained by selective immune recall of memory T cells that recognize conserved epitopes between DENV and ZIKV^23^, which also has previously been demonstrated during secondary heterotypic DENV infections^67,68^. In addition, an increased cytotoxic profile as demonstrated by the higher frequency of CD107a-expressing CD4^+^ and CD8^+^ T cells in the ZIKV mid-convalescent group correlates with the synchronously early activation of CD4^+^ and CD8^+^ effector memory T cells and elevated levels of perforin release.

Higher proportion of IFN-γ and TNF-α producing T cells before a secondary heterologous DENV infection has been associated to a subsequent subclinical outcome^69^. Herein, we observed that the ZIKV mid-convalescent group had elevated levels of IFN-γ and TNF-α producing T cells since baseline. In this group, DENV infection stimulated a higher frequency of these cells, but remarkably, also increased highly cross-reactive IFN-γ-producing CD4^+^ T cells directed to DENV E, and ZIKV E/NS proteins. Memory CD4^+^ T cells are required to generate an effective humoral response against ZIKV^70^. Based on this, the higher proportion of DENV-E-reactive IFN-γ-producing CD4^+^ T cells may play a role in the induction of the robust Ab response in the ZIKV mid-convalescent group against ZIKV and all DENV serotypes. On the other hand, we showed that naïve animals with DENV *de novo* response did not cross-neutralized ZIKV at all, which state that although similar, antigenic differences are sufficient to mount predominantly type-specific rather than cross-reactive responses during a primary infection^50,56^.

A lack of ZIKV immunity promoted a more pro-inflammatory profile after DENV infection characterized by significant elevated levels of IL-6 and MIG/CXCL9. Interestingly, higher levels of IFN-α were observed in the ZIKV-naïve animals. This antiviral cytokine is known to be actively produced during acute DENV infection *in vitro* and *in vivo*^71^. Elevated levels have been correlated with severity in DHF patients, and to act as a marker for elevated DENV replication^72,73^. On the other hand, the presence of a longer ZIKV convalescence is associated with increased levels of CXCL10 and perforin. CXCL10 is an immune mediator for T cells proliferation, recruitment of CD4^+^ and CD8^+^ activated T cells and IFN-γ-producing CD8^+^ T cells, required to control DENV infection *in vivo*^74,75^. This correlates with higher proportion and activation of both T cell compartments and subsequent functional T cell response against DENV-E-specific peptides in the group with longer convalescence to ZIKV. Perforin is involved in the cytotoxic degranulation process against virus-infected cells. In DENV infection, perforin is part of the anti-DENV cytotoxic phenotype of CD8^+^ and CD4^+^ T cells^48,76^. Perforin levels were significantly elevated only in the ZIKV mid-convalescent group after DENV infection. Accordingly, this coincides with a significant activation of CD8^+^ and CD4^+^ effector memory T cells, and degranulation functional response of both T cell compartments, suggesting an enhanced perforin-producing cytotoxic role of T cells in presence of longer convalescence to ZIKV. Contrary to our findings, a previously published work found that an approximately two month ZIKV immunity period resulted in an increase of pro-inflammatory cytokines^37^. However, a differential effect due to the use of different sample types (plasma vs serum) between both studies cannot be ruled out.

One limitation of our study is the utilization of low numbers of animals per group. Additional studies with a larger number of animals are warranted. However, fundamental and seminal contributions on ZIKV and ZIKV/DENV interactions have been obtained by using similar limited number of animals per group^16,17,77,78^ . Another limitation is that our study monitored the immune response up to 90 days after DENV infection. Additional longitudinal studies are needed to test the immune response over longer periods of time including subsequent DENV heterotypic challenges to evaluate the efficacy of the memory recall in cross-protection between serotypes. Finally, we cannot comment about the likelihood to increase or decrease susceptibility to develop DHF/DSS in the context of ZIKV immunity since DENV clinical manifestations in NHP models are limited and are characterized to be subclinical infections ^28^.

In summary, dissecting our main findings per previous ZIKV-immune status we found that a ZIKV middle-convalescence: (i) results in shorter DENV viremic period, (ii) lowest pro-inflammatory status with upregulation of cellular immune response mediators, (iii) robust neutralizing antibody response higher in magnitude and durability against ZIKV strains and DENV serotypes, (iv) elevated activated and proliferating B cells, (v) early activation of cross-reactive CD4^+^ and CD8^+^ effector memory T cells, (v) and a major breadth of functional T cell response. For ZIKV early-convalescence we demonstrated: (i) average DENV viremic period and no exacerbation of pro-inflammatory status, (ii) neutralizing antibody response with high magnitude but less durability against ZIKV strains and DENV serotypes compared to the ZIKV middle-convalescent group, (iii) early activation of central memory CD8^+^ T cells, (iv) and very limited activation of effector memory T cells. For the ZIKV-naïve group we demonstrated: (i) longer DENV viremic period and pro-inflammatory status, (ii) a more delayed *de novo* neutralizing antibody response against DENV serotypes and inability to neutralize ZIKV strains, (iii) a limited B cell response, (iv) and an overall *de novo* T cell response lower in magnitude and cross-reactivity compared to ZIKV-immune groups.

This study reinforces the usefulness of NHPs as a suitable model to characterize the immune response elicited by heterologous and consecutive flavivirus infections and to identify differential modulation of the immune response influenced by the time interval between infections. Our findings of highly cross-reactive response against DENV in presence of previous ZIKV immunity with no exacerbation of DENV pathogenesis may contribute to explain the decrease of detected DENV cases after ZIKV epidemic in the Americas. This scenario has been suggested recently using a fewer number of animals^38^. Furthermore, our data show a positive scenario that supports the implementation of ZIKV vaccine programs, since it suggests that a vaccine-acquired ZIKV-immunity may not worsen DENV pathogenesis and may ameliorate immune response against a subsequent infection with DENV. Similarly, the implementation of DENV vaccines is also supported in the context of previous ZIKV immunity, since ZIKV convalescence may boost the vaccine-acquired anamnestic immune response to DENV without predisposing to an enhanced pathogenesis. However, the selection of the vaccine schedule may be critical to induce the optimal immune response when more than one doses are planned.

## Methods

### Cell Lines

Aedes *albopictus* cells, clone C6/36 (ATCC CRL-1660), whole mosquito larva cells, were maintained in Dulbecco Minimum Essential Medium (DMEM) (GIBCO, Life Technologies) supplemented with 10% fetal bovine serum (FBS) (Gibco) and 1% Penicillin/Streptomycin (P/S) (Gibco). C6/36 were used to produce previous ZIKV and DENV viral stocks with high titers in 150-175 cm^2^ cell culture flasks (Eppendorf), and incubated at 33°C and 5% CO_2_. Vero cells, clone 81 (ATCC CCL-81), African green monkey kidney epithelial cells, were maintained with DMEM supplemented with 10% FBS and 1% of P/S, HEPES, L-glutamine and non-essential amino acids (NEAA) in 75 cm^2^ cell culture flasks, and incubated at 37°C and 5% CO_2_. Vero-81 cells were used for the cells monolayer in viral titrations by plaque assays and plaque reduction neutralization test (PRNT) in flat-bottom 24-well plates (Eppendorf).

### Viral Stocks

The DENV-2 New Guinea 44 (NGC) strain (kindly provided by Steve Whitehead, NIH/NIAID, Bethesda, Maryland, USA), known to replicate well in rhesus macaques, was used for the challenge in order to obtain comparative results with previous published studies from our group on DENV and ZIKV challenge studies^16,33^. We have standardized the assays to quantify this virus by Plaque assay, as described in our previous work^16^. The titer of DENV-2 for the challenge was 5×10^7^ pfu/ml. In addition, ZIKV H/PF/2013 strain (kindly provided by CDC-Dengue Branch, San Juan, Puerto Rico), ZIKV PRVABC59 (ATCC VR-1843), DENV-1 Western Pacific 74, DENV-3 Sleman 73, and DENV-4 Dominique strains (kindly provided by Steve Whitehead from NIH/NIAID, Bethesda, Maryland, USA) were propagated in C6/36 cells, titrated and used for Plaque Reduction Neutralization Test (PRNT) assays.

### Viral Titration Plaque Assay

DENV titrations by plaque assay were performed seeding Vero-81 (∼8.5×10^4^ cells /well) in flat bottom 24-well cell culture well plates (Eppendorf) in supplemented DMEM the day before. Viral dilutions (10-fold) were made in diluent media [Opti-MEM (Invitrogen) with 2% FBS (Gibco) and 1% P/S (Gibco)]. Prior to inoculation, growth medium was removed and cells were inoculated with 100 ul/well of each dilution in triplicates. Plates were incubated for 1 hr, 37°C, 5% CO_2_ and rocking. After incubation, virus dilutions were overlaid with 1 ml of Opti-MEM [1% Carboxymethylcellulose (Sigma), 2% FBS, 1% of NEAA (Gibco) and P/S (Gibco)]. After 3 to 5 days of incubation (days vary between DENV serotypes), overlay was removed and cells were washed twice with phosphate buffered saline (PBS), fixed in 80% methanol (Sigma) in PBS, and incubated at room temperature (RT) for 15 minutes. Plates were blocked with 5% Non-fat dry milk (Denia) in PBS for 10 minutes. Blocking buffer was discarded and 200 ul/well of primary antibodies mix [anti-E protein monoclonal antibody (mAb) 4G2 and anti-prM protein mAb 2H2 (kindly provided by Aravinda de Silva and Ralph Baric, University of North Carolina Chapel Hill, North Carolina, USA), both diluted 1:250 in blocking buffer] were added and incubated for 1 hr, 37°C, 5% CO_2_ and rocking. Plates were washed twice with PBS and incubated in same conditions with horseradish peroxidase (HRP)-conjugated goat anti-mouse secondary antibody (Sigma), diluted 1:1000 in blocking buffer. Plates were washed twice with PBS and 150 ul/well of TrueBlue HRP substrate (KPL) were added and plates were incubated from 1-10 minutes at RT until plaque-forming units (pfu) were produced and visible. Then 200 ul/well of distilled water were added to stop the substrate reaction, plates get dry and pfu were counted to calculate viral titers.

### Macaques and Viral Challenge

From 2008 to 2015, the Caribbean Primate Research Center (CPRC) funded a large DENV research program. Multiple studies made available several cohorts of rhesus macaques (*Macaca mulatta*) infected with different DENV serotypes in distinct timelines and also naïve cohorts were available as well. After our laboratories prioritized ZIKV research since 2016, DENV pre-exposed and naïve cohorts were infected with ZIKV and pre-exposed animals became available for this study. All animals were housed within the Animal Resources Center facilities at the University of Puerto Rico-Medical Sciences Campus (UPR-MSC), San Juan, Puerto Rico. All the procedures were performed under the approval of the Institutional Animal Care and Use Committee (IACUC) of UPR-MSC and in a facility accredited by the Association for Assessment and Accreditation of Laboratory Animal Care (AAALAC file # 000593; Animal Welfare Assurance number A3421; protocol number, 7890116). Procedures involving animals were conducted in accordance with USDA Animal Welfare Regulations, *the Guide for the Care and use of Laboratory Animals* and institutional policies to ensure minimal suffering of animals during procedures. All invasive procedures were conducted using anesthesia by intramuscular injection of ketamine at 10-20 mg kg^-1^ of body weight. Rhesus macaques from the CPRC are very well genetically characterized from a common stock introduced in 1938 at Cayo Santiago, an islet located in the southeast of Puerto Rico. These macaques with Indian genetic background are part of the purest colony used in the United States for comparative medicine and biomedical research^49^

The experimental design was based on 14 young adult male rhesus macaques divided in three cohorts. Cohort 1 (ZIKVPF-10mo): composed of four animals (5K6, CB52, 2K2, and 6N1) that were inoculated with 1×10^6^ pfu/500 ul of the ZIKV H/PF/2013 strain subcutaneously^16^ 10 months before DENV-2 challenge. Cohort 2 (ZIKVPR-2mo): composed of 6 animals (MA067, MA068, BZ34, MA141, MA143, and MA085) that were inoculated with 1×10^6^ pfu/500 ul of the ZIKV PRVABC59 strain two months before DENV-2 challenge. Both ZIKV strains used for previous exposure of these groups are >99.99% comparable in amino acid identity (Supplementary Table 1). Cohort 3 (Naïve): composed of four ZIKV/DENV naïve animals (MA123, MA023, MA029, and MA062) as a control group. Cohort 1 and 3 were challenged on the same day while cohort 2 was challenged 3 months later with the same stock of DENV-2. However, all samples were frozen and analyzed together, except for the immunophenotyping analysis.

The ages of all animals are within the age range for young adults rhesus macaques https://www.nc3rs.org.uk/macaques/macaques/life-history-and-diet/ (ZIKVPF-10mo: 6.8, 6.8, 5.8, and 5.9; ZIKVPR-2mo: 6.4, 6.5, 5.2, 4.3, 5.6, and 5.5; Naïve: 4.8, 6.6, 6.8, and 5.7). Prior to DENV-2 challenge all animals were subjected to quarantine period. All cohorts were bled for baseline and challenged subcutaneously (deltoid area) with 5×10^5^ pfu/500 ul of DENV-2 New Guinea 44 strain. After DENV-2 challenge all animals were extensively monitored by trained and certified veterinary staff for evidence of disease and clinical status: external temperature (°C) with an infrared device (EXTECH Instruments, Waltham, MA), weight (Kg), CBC and CMP. All animals were bled once daily from day 1 to day 10 and after that on days 15, 30, 60 and 90 post-infection (dpi). In all timepoints the blood samples were used for serum separation (Baseline, 7, 30, 60, 90 dpi only). PBMCs were collected at same time points using CPT tubes (BD-Biosciences, San Jose, CA) containing citrate. Additional heparin samples were obtained for immunophenotyping by flow cytometry using fresh whole blood. Fig. 1 shows the experimental design and samples collection timeline.

### DENV RNAemia

DENV viral RNA extraction was performed from acute serum samples (Baseline, 1-10, and 15 dpi) using QIAamp Viral RNA mini kit (Qiagen Inc, Valencia, CA, USA) according to the manufacturer’s instructions. RNAemia levels were measured by a One-Step qRT-PCR detection kit (Oasig, Primerdesign Ltd., UK) and using DENV RT primer/probe Mix kit (Genesig, Primerdesign Ltd., UK) according to the manufacturer’s protocol (catalog No. oasig-onestep). Primers are designed to target the 3’ untranslated region (3’ UTR) of all four DENV serotypes and have 100% homology with over 95% of reference sequences contained in the NCBI database. Assays were performed in an iCycler IQ5 Real-Time Detection System with Optical System Software version 2.1 (Bio-Rad, Hercules, CA, USA). Limit of detection (LOD) was 20 copies per ml. Furthermore, in order to correlate RNAemia levels with DENV pathogenesis we monitored the clinical status for injury and/or clinical manifestations. Complete Blood Counts (CBC) were performed for all animals in several timepoints (Baseline, 7, and 15 dpi) to determine the absolute number (106 cells/ml) and percent (%) of lymphocytes (LYM), monocytes (MON), white blood cells (WBC), neutrophils (NEU) and platelets (PLT). Also, Comprehensive Metabolic Panel (CMP) were evaluated in several timepoints (Baseline, 7, 15 and 30 dpi) to measure concentration (U/L) of alkaline phosphatase and liver enzymes alanine aminotransferase (ALT) and aspartate aminotransferase (AST).

### ELISA

Seroreactivity to DENV and cross-reactivity to ZIKV was measured at different timepoints before and after DENV-2 challenge. DENV-IgM (Focus Diagnostics, Cypress, CA, USA) was quantified at baseline, 5, 10, 15 and 30 dpi. DENV-IgG was quantified at baseline, 7, 15, 30, 60 and 90 dpi (Focus Diagnostics, Cypress, CA, USA). To determine the modulation of serological profile against ZIKV we assessed: levels of anti-ZIKV IgM (InBios, Seattle, WA, USA) at baseline, 5, 10, 15 and 30 dpi; anti-ZIKV IgG (XPressBio, Frederick, MD, USA) at baseline, 7, 15, 30, 60 and 90 dpi; anti-ZIKV NS1-IgG (Alpha Diagnostics, San Antonio, TX, USA) at baseline, 30, 60 and 90 dpi (including additional timepoints prior baseline for both ZIKV-immune groups); and anti-ZIKV EDIII-IgG (Alpha Diagnostics International, San Antonio, TX, USA). All ELISA-based assays were performed following the manufacturers’ instructions. This serological characterization allows us to assess the dynamics of DENV and ZIKV cross-reactivity but without discerning between cross-reactive binding Abs and cross- or type-specific neutralizing Abs.

### Plaque Reduction Neutralization Test (PRNT)

Selected serum samples (baseline, 30, 60 and 90 dpi) were challenged to neutralized ZIKV (H/PF/2013, PRVABC59), DENV-1 Western Pacific 74, DENV-2 NGC 44, DENV-3 Sleman 73, and DENV-4 Dominique strains. For the infecting serotype (DENV-2) and ZIKV the NAbs were measured in early timepoints as well (7 and 15 dpi). For the PRNT, serum samples were inactivated, diluted (2-fold), mixed with a constant inoculum of virus (volume necessary to produce ∼35 pfu/well) and then incubated for 1 hr at 37°C and 5% CO_2_. After incubation, virus-serum mix dilutions were added to Vero-81 cells monolayer in flat bottom 24-well plates seeded the day before for 1 hr at 37°C and 5% CO_2_, finally overlay medium was added and incubated by several days (serotype dependent). Results were reported as PRNT60 titers, NAb titer capable of reduce 60% or more of DENV serotypes or ZIKV strains pfu compared with the mock (control of virus without serum). A PRNT60 1:20 titer was considered a positive Neut titer, and <1:20 as a negative Neut titer. Non-neutralizing titers (<1:20) were assigned with one-half of the limit of detection for graphs visualization.

### Multiplex Cytokine Profile

A total of 8 cytokines/chemokines were measured (pg /ml^-1^) by Luminex at baseline, 1, 2, 3, 5, 10, 15 and 30 dpi, including: interferon alpha (IFN-α), interleukin-6 (IL-6), monokine induced by IFN-gamma (MIG/CXCL9), monocyte chemoattractant protein 1 (MCP-1/CCL2), macrophage inflammatory protein 1-beta (MIP-1β/CCL4), IL-1 receptor antagonist (IL-1RA), C-X-C motif chemokine 10 (CXCL10/IP-10) and perforin. The multiplex assay was conducted as previously described^16,79^.

### Immunophenotyping

Flow cytometry (MACSQuant Analyzer 10, Miltenyi Biotec) analysis was performed to determine the frequency, activation and proliferation of cell populations of the innate and adaptive immune response based on the phenotyping strategy of a previous study^16^ (Supplementary Fig. 7, 8, and 9 for gating strategy; Supplementary Table 2 for Ab panel). Phenotypic characterization of macaque PBMCs from fresh whole blood samples was performed by 8-multicolor flow cytometry using fluorochrome conjugated Abs at several timepoints (Baseline, 1, 2, 3, 7, 10 dpi; and 15 and 30 dpi for B/T cell panel only). Single cells (singlets) were selected by their FSC area (FSC-A) and height (FSC-H) patterns. Lymphocytes (LYM) were gated based on their characteristic forward and side scatter pattern (FSC, SSC). T cells were selected gating on the CD3^+^ population. CD4^+^ and CD8^+^ T cells were defined as CD3^+^CD4^+^ and CD3^+^CD8^+^, respectively. Naive (N; CD28^+^CD95^-^), effector memory (EM; CD28^-^ CD95^+^) and central memory (CM; CD28^+^CD95^+^) T cell subpopulations were determined within CD4^+^ and CD8^+^ T cells. B cells were defined as CD20^+^CD3^-^. The activation of B and T cell memory subpopulations (EM and CM) was assessed by the presence of the early activation marker CD69. Proliferation of total and activated B cells was quantified by the expression of the intracellular marker Ki67. Natural killer (NK) cells were defined as CD3^-^CD20^-^CD14^-^ and analyzed by the double positive expression of the following NK cell markers: CD8, CD56, NKG2A, NKp30, and NKp46 (Supplementary Fig. 9 for gating strategy). Dendritic cells (DC) were separated in two populations within the Lineage-DR+ (HLA-DR^+^ CD3^-^ CD14^-^ CD16^-^ CD20^-^ CD8^-^ NKG2A^-^) by the expression of CD123 (plasmacytoid, pDC) or CD11c (myeloid, mDCs) (Supplementary Fig. 8 for gating strategy). Then, DC percentages were calculated from total PBMCs (total events of the DC subpopulation divided by total PBMCs and multiplied by 100). The phenotyping assays were optimized and performed as previously published^16,33,80^.

### T Cell Functional Response

Intracellular cytokine staining of macaques PBMCs was performed by multicolor flow cytometry using methods previously described (Supplementary Fig. 14 for gating strategy; Supplementary Table 3 for Ab panel)^16,80^. Functional effector response of CD4^+^ and CD8^+^ T cells was measured before and after DENV infection. Antigen-specific CD4^+^ and CD8^+^ T cell effector responses were measured at baseline to determine basal levels in presence (ZIKVPF-10mo, ZIKVPR-2mo) or absence (Naïve) of previous immunity to ZIKV. Also, 30, 60 and 90 dpi were assessed to determine how this pre-existing functional response is modulated after DENV infection and if is maintained over time. For peptide pools stimulation, PBMCs were stimulated for 6 hr at 37°C and 5% CO_2._ The peptides used for DENV-E, ZIKV-E and ZIKV-NS were 15-mers overlapped by 10 amino acids at 1.25 ug/ml^-1^, 2.5 ug/ml^-1^, 475 ng/ml^-1^ per peptide, respectively (Supplementary Table 4 for peptide sequences). The stimulation with peptides was performed in presence of brefeldin A at 10 ug/ml^-1^. After stimulation, the cells were stained for the following markers: CD3, CD4, CD8, CD20 (excluded), CD107a (functional cytotoxicity). Levels of IFN-γ and TNF-α also were measured in gated lymphocytes cell populations. Samples were measured and data was collected on a LSRII (BD).

### Statistical Analysis

Statistical analyses were performed using GraphPad Prism 7.0 software (GraphPad Software, San Diego, CA, USA). The statistical significance between the means of all groups were determined using Two-way ANOVA Multiple Comparison Tukey Test, and to compare each mean against the baseline mean within same group Two-way ANOVA Multiple Comparison Dunnett Test was performed. Total number of families and comparisons per family used for adjustments are depicted in each figure legend. Significant multiplicity adjusted *p* values (* <0.05, ** <0.01, *** <0.001, **** <0.0001) show statistically significant difference between groups (Tukey Test) or timepoints within a group (Dunnett Test).

### Data Availability

All relevant data is in main figures and supplementary information, any additional details are available from authors upon request. The RAW data from all main and supplementary figures is provided as a Source Data File

## Acknowledgments

We thank all the staff of the Caribbean Primate Research Center and Animal Resources Center for their continuous support with the sample collection, schedule, and monitoring of the animals. Authors recognize the support provided by Dr. Elmer Rodriguez reviewing the statistics. This work was supported by the following grants: 2 P40 OD012217 and 2U42OD021458-15 to C.A.S. and M.I.M., K22AI104794 to J.D.B., P51OD011133 (L.G.), HHSN272201400045C to D.W., and R25GM061838 to E.X.P.-G.

## Author Contributions

C.A.S. and E.X.P.-G. developed the experimental design. I.V.R. supervised and performed sample collection and animals monitoring. E.X.P.-G., P.P., C.S.-C., M.A.H., A.O.-R., V.H., L.P., L.C., and T.A. performed the experiments. E.X.P.-G., C.A.S., V.H., M.A.H., L.J.W., A.d.S., and D.W. analyzed the data. E.X.P.-G. and C.A.S. drafted the manuscript. C.A.S., E.X.P.-G., D.W., A.K.P., J.D.B., M.A.H., L.G., L.J.W., and A.d.S. revised the manuscript.

## Additional Information

**Supplementary Information** available at:

