## Supplementary Figures and Tables for "Time elapsed between Zika and dengue virus infections affects antibody and T cell responses"

Pérez-Guzmán *et al.*

### Supplementary Information

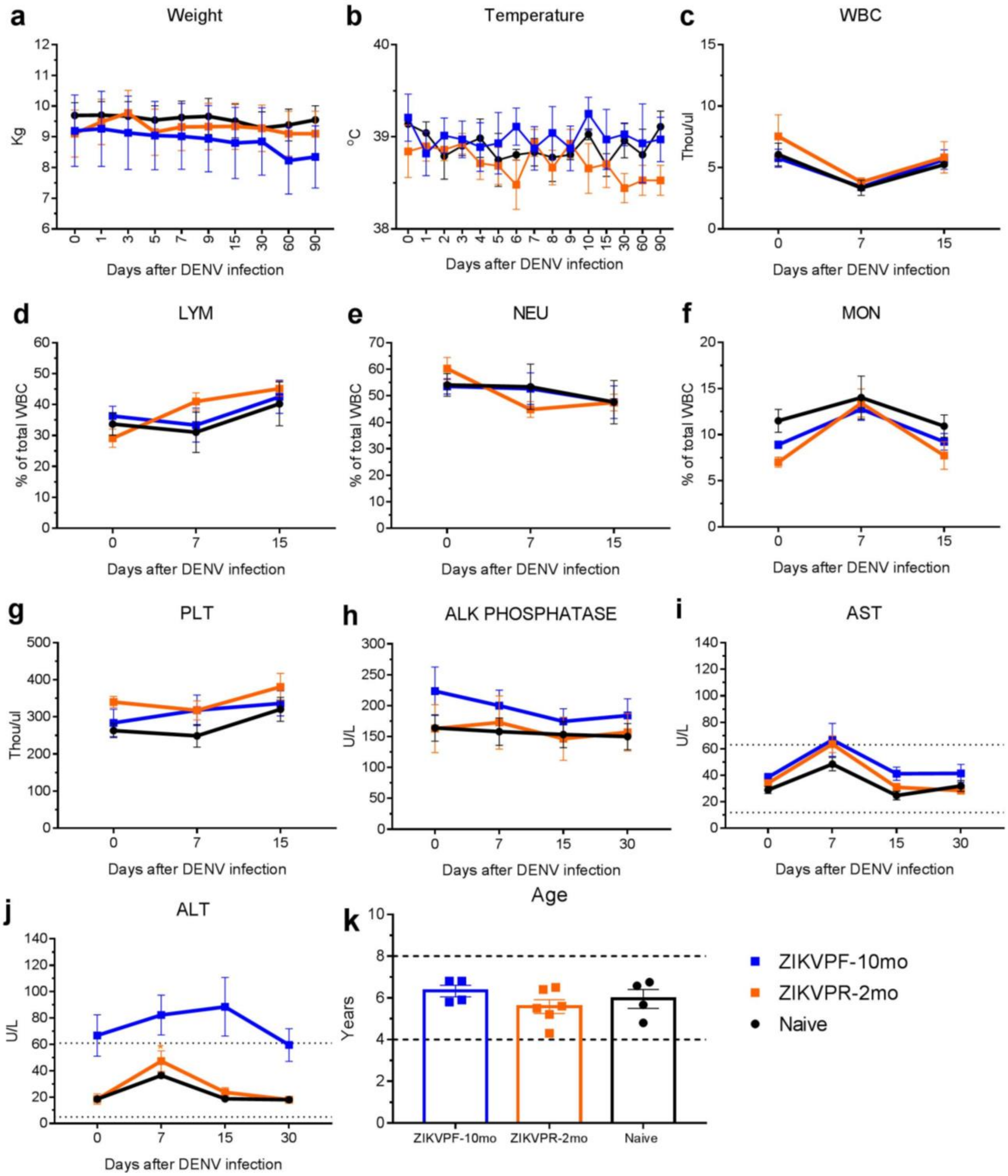


**Supplementary Figure 1 | Clinical status and vital signs kinetics in ZIKV-immune and naïve macaques.** (**a**) Weight (kg) was measured at baseline, 1, 3, 5, 7, 9, 15, 30, 60 and 90 dpi. (**b**) Temperature (°C) was monitored with an infrared device at baseline, 1-10, 15, 30, 60 and 90 dpi. Complete blood cell counts (CBC) parameters (thou/ul and/or % of total WBC) such as (**c**) white blood cells (WBC), (**d**) lymphocytes (LYM), (**e**) neutrophils (NEU), (**f**) monocytes (MON), and (**g**) platelets (PLT) were screened at baseline, 7, and 15 dpi. Comprehensive metabolic panel (CMP)

was performed to assess levels (U/L) of (**h**) alkaline phosphatase (ALK PHOSPHATASE) and liver enzymes (**i**) aspartate transaminase (AST), and (**j**) alanine transaminase (ALT) at baseline, 7, 15 and 30 dpi. Normal range of AST and ALT are depicted for reference. (**k**) Age of rhesus macaques are depicted including the range of young adults for reference. Symbols represent mean level detected for each parameter per cohort per timepoint: blue squares (ZIKVPF-10mo), orange squares (ZIKVPR-2mo) and black circles (Naïve). Lines connect mean values detected over time. Error bars indicate the standard error of the mean (SEM) for each cohort per timepoint. Statistically significant differences between groups were determined using Two-Way Anova adjusted for Tukey's multiple comparisons test including 10, 15, 3, 4, and 3 families for panel a, b, c-g, h-j, and k, respectively, and 3 comparisons per family. For differences in ALT levels Two-Way Anova Dunnett's multiple comparisons test (comparison of each group response at each timepoint versus baseline of the same group) was performed including 3 families, and 3 comparisons per family due to divergence of non-specific levels between cohorts at baseline. Statistically differences are reported as multiplicity adjusted *p* values (* <0.05).

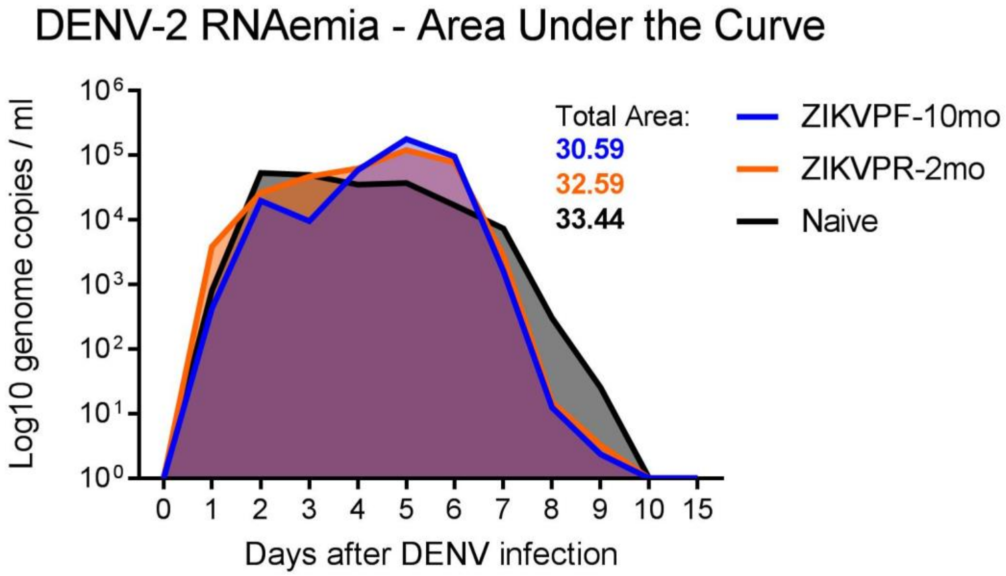


**Supplementary Figure 2 | Previous ZIKV immunity modulates DENV RNAemia kinetics and is associated with a lower area under the curve.** The area under the curve (AUC) was calculated using log-transformed values of DENV-2 RNAemia in ZIKV-immune and naïve animals. The total area by group is depicted on the graph as light blue, light orange and gray for ZIKVPF-10mo, ZIKVPR-2mo, and Naïve, respectively. Lines mark the mean value of genome copies per group per timepoint. A value of 1 was assigned to all samples below the LOD in order to calculate the means.

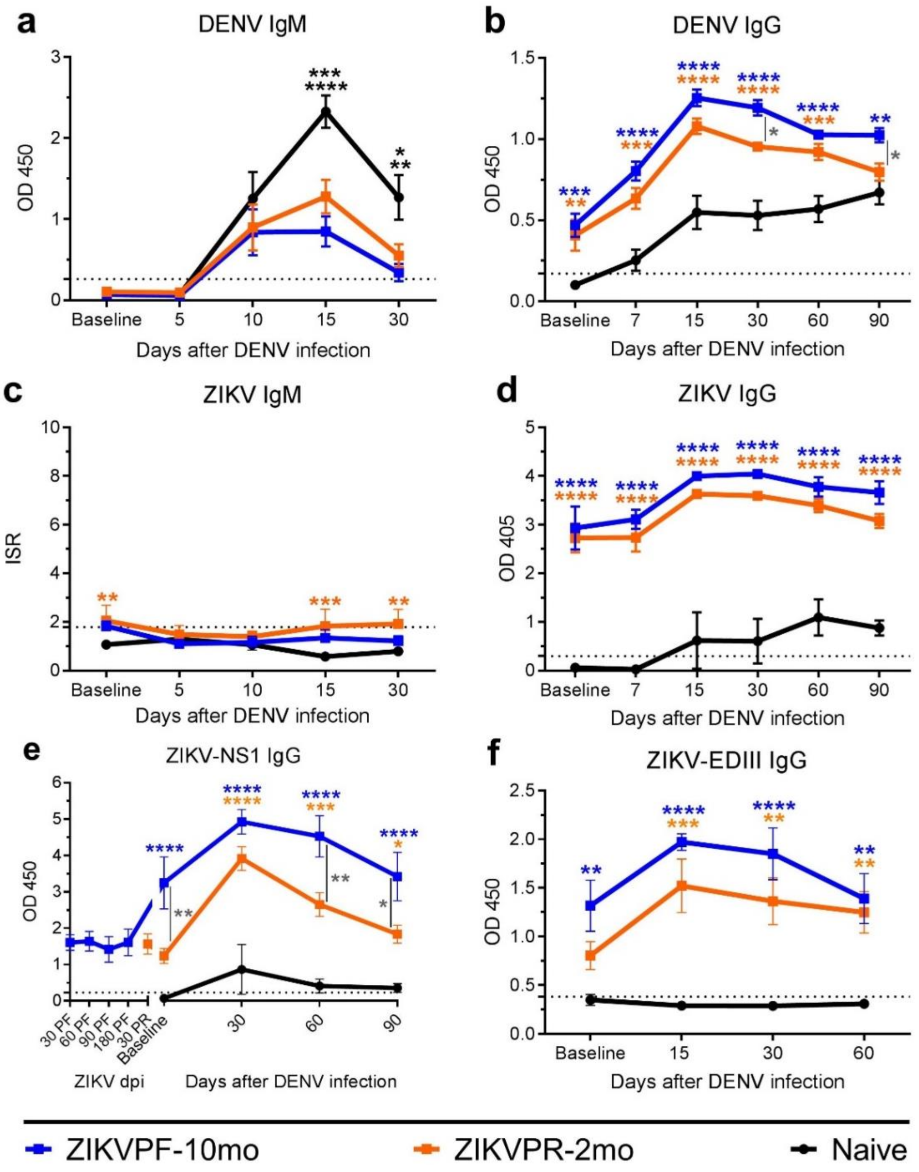


**Supplementary Figure 3 | Serological cross-reactivity is boosted by ZIKV immunity.** Levels of DENV (**a**) IgM and (**b**) IgG, and ZIKV (**c**) IgM, (**d**) IgG, (**e**) NS1-IgG and (**f**) EDIII-IgG were measured by ELISA at multiple timepoints before and after DENV infection. Symbols connected with full lines represent mean levels of Abs detected per cohort over time: blue squares (ZIKVPF-10mo), orange squares (ZIKVPR-2mo) and black circles (Naïve). Panel e includes additional timepoints before DENV infection for ZIKV-immune groups: 30, 60, 90 and 180 days after ZIKV (H/PF/2013) infection for the ZIKVPF-10mo group, and 30 days after ZIKV (PRVABC59) infection for the ZIKVPR-2mo group. Error bars indicate the standard error of the mean (SEM) and dotted line mark the limit of detection for each individual ELISA. Results were read at OD 450, 405 or using ISR (Immune Status Ratio) following manufacturer's instructions. Statistically significant differences between groups were calculated using Two-Way Anova adjusted for Tukey's multiple comparisons test including 5, 6, 9, and 4 families, and 3 comparisons per family. Significant multiplicity adjusted *p* values (* <0.05, ** <0.01, *** <0.001, **** <0.0001) are shown. Blue and orange asterisks represent significant difference between the corresponded ZIKV immune groups and naive group, and gray asterisks indicate a significant difference between ZIKV immune groups.

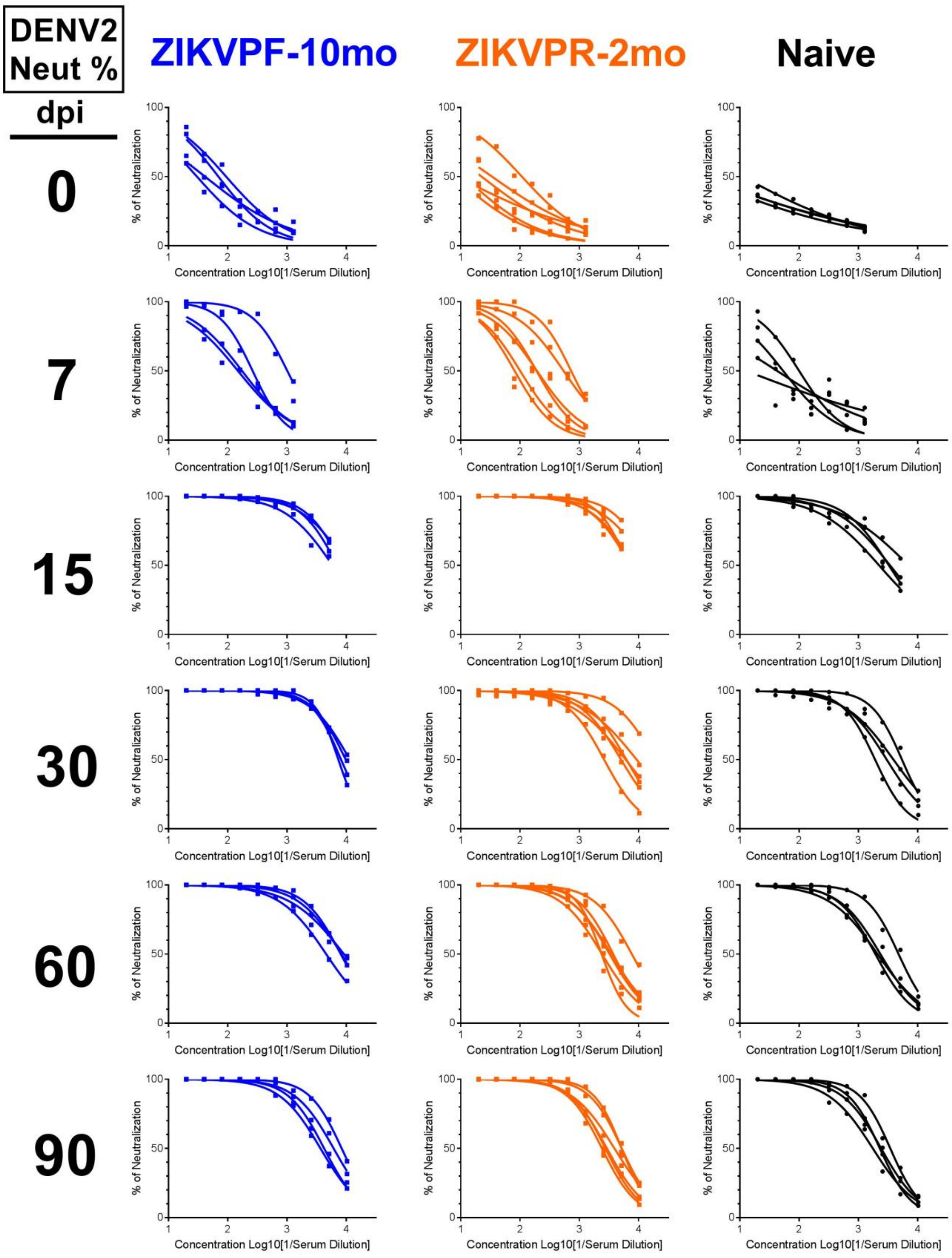


**Supplementary Figure 4 | Neutralization kinetics against DENV-2.** Percentage of DENV-2 neutralization of each animal per group calculated by the transformation of PRNT60 Neut 2-fold titers into Log10 (1/serum dilution), and sigmoidal-dose response curves were generated. Each column of panels represent the % of DENV-2 neutralization for each group (ZIKVPF-10mo: blue squares/curves; ZIKVPR-2mo: orange squares/curves; Naïve: black circles/curves) and each row of panels represent a timepoint before and after DENV infection (baseline, 7, 15, 30, 60, 90 dpi).

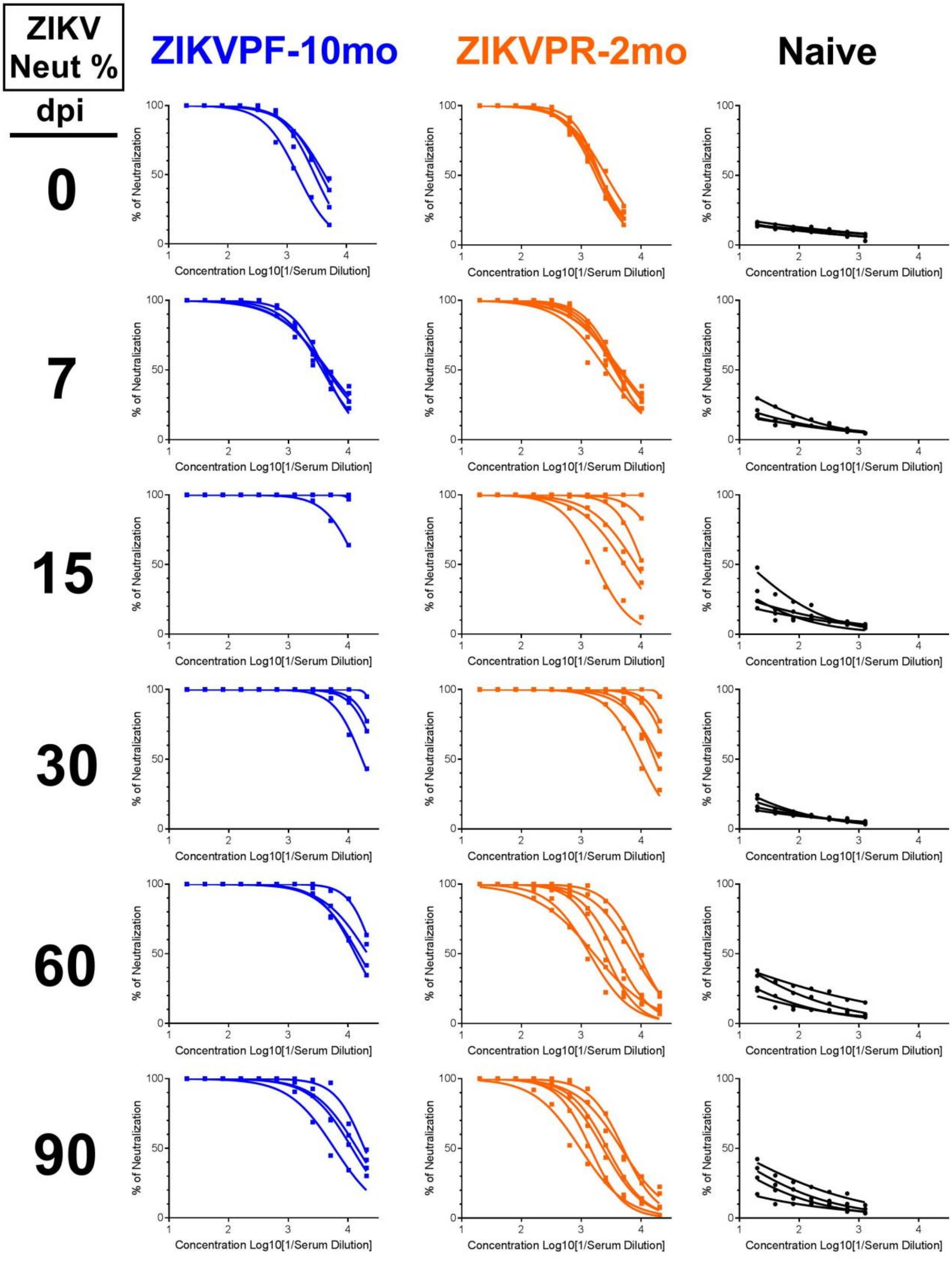


**Supplementary Figure 5 | Neutralization kinetics against ZIKV.** Percentage of ZIKV (H/PF/2013) neutralization of each animal per group calculated by the transformation of PRNT60

Neut 2-fold titers into Log10 (1/serum dilution), and sigmoidal-dose response curves were generated. Each column of panels represent the % of ZIKV neutralization for each group (ZIKVPF-10mo: blue squares/curves; ZIKVPR-2mo: orange squares/curves; Naïve: black circles/curves) and each row of panels represent a timepoint before and after DENV infection (baseline, 7, 15, 30, 60, 90 dpi).

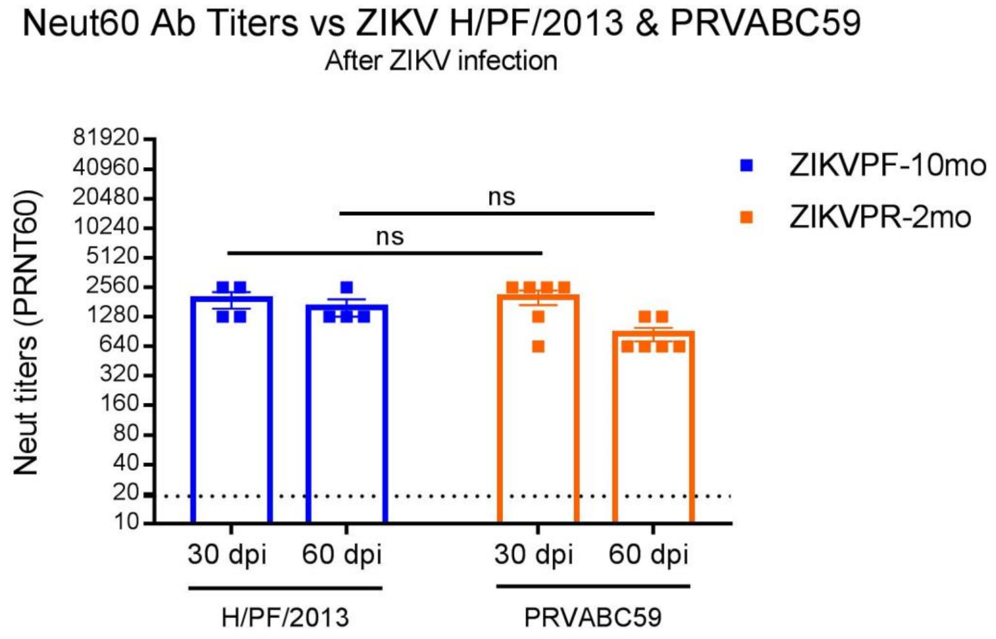


**Supplementary Figure 6 | Similar neutralizing titers induced by two different ZIKV strains.** NAb titers against H/PF/2013 and PRVABC59 ZIKV strains for ZIKVPF-10mo and ZIKVPR-2mo groups, respectively, were determined by PRNT60 at 30 and 60 after ZIKV infection. Symbols indicate levels of NAb titers detected per animal: blue squares (ZIKVPF-10mo), and orange squares (ZIKVPR-2mo). Error bars represent the standard error of the mean (SEM). PRNT60: NAb titer capable of reduce 60% or more of ZIKV strains plaque-forming units (pfu) compared with the mock (control of virus without serum). A PRNT60 1:20 titer was considered positive, and <1:20 as a negative Neut titer. Dotted line mark <1:20 for negative results. Statistically significant differences (ns: not significant) between two groups were calculated using Two-Way Anova corrected for Sidak's multiple comparisons test including 1 family, and 2 comparisons within the family.

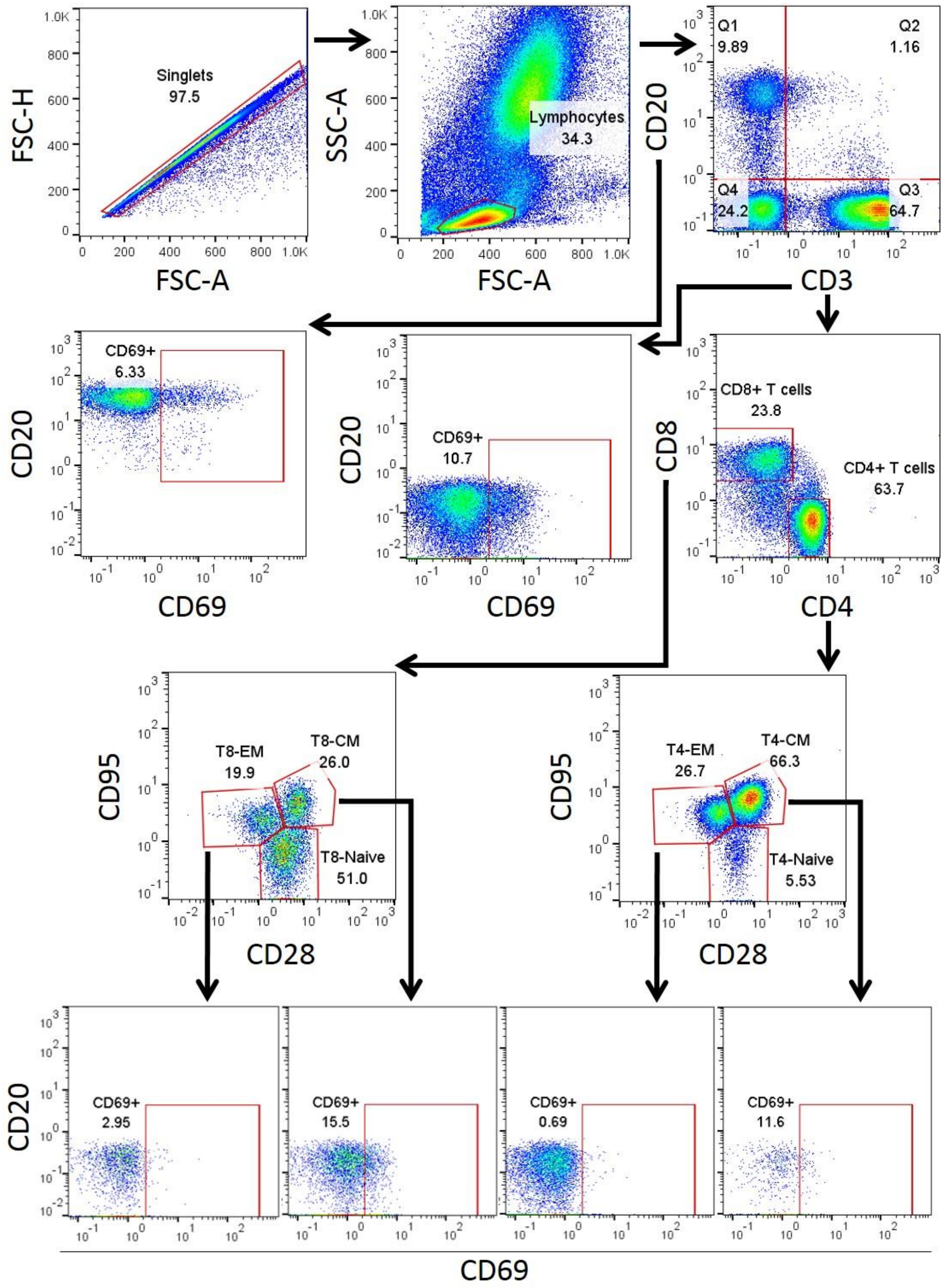


**Supplementary Figure 7 | Gating strategy for immunophenotyping and activation of B cells, and memory T cell subpopulations.** Single cells (singlets) were selected by their FSC area

(FSC-A) and height (FSC-H) patterns. Lymphocytes (LYM) were gated based on their characteristic forward and side scatter pattern (FSC, SSC). T cells were selected gating on the $CD3^+$ population. $CD4^+$ and $CD8^+$ T cells were defined as $CD3^+CD4^+$ and $CD3^+CD8^+$, respectively. Naive (N; $CD28^+CD95^-$), effector memory (EM; $CD28^-CD95^+$) and central memory (CM; $CD28^+CD95^+$) T cell subpopulations were determined within $CD4^+$ and $CD8^+$ T cells. B cells were defined as $CD20^+CD3^-$. The activation of B and T cell memory subpopulations (EM and CM) was assessed by the presence of the early activation marker CD69.

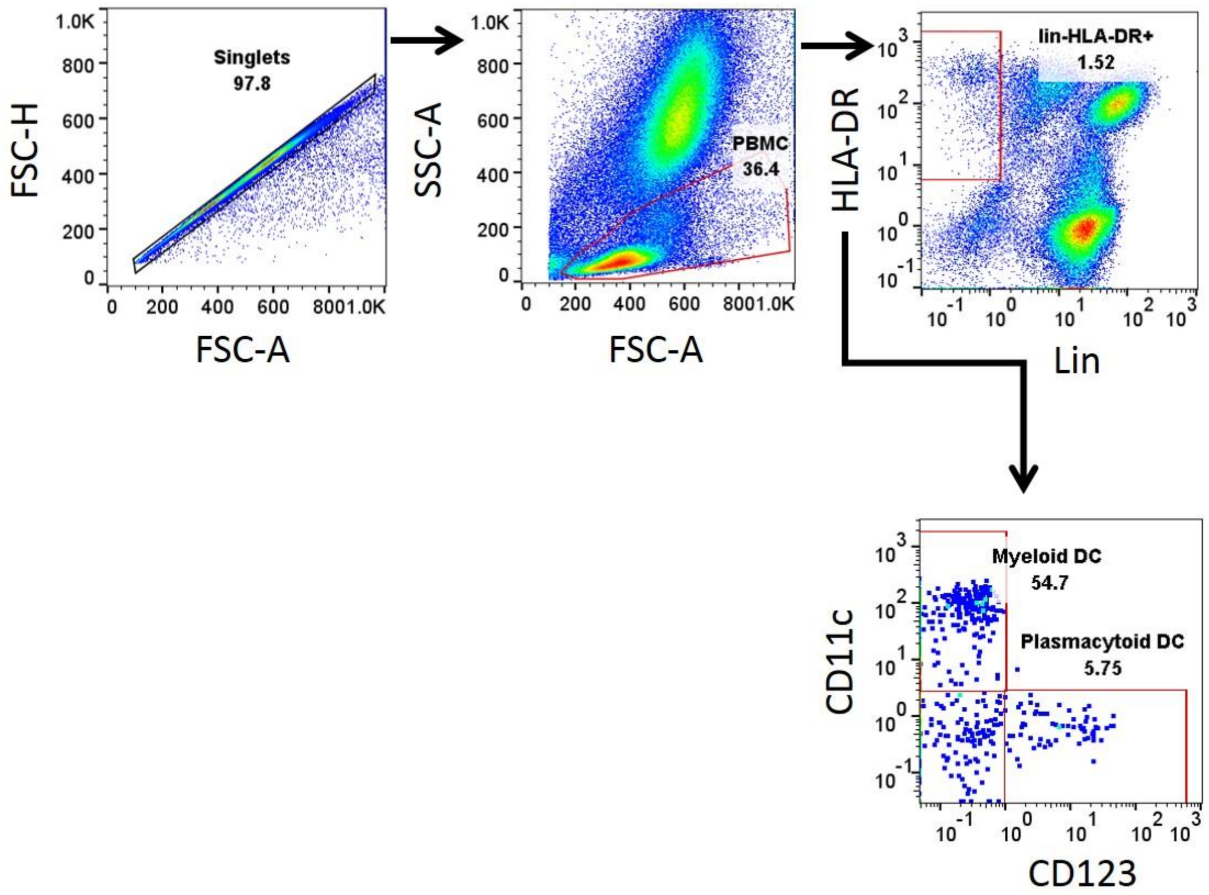


**Supplementary Figure 8 | Gating strategy for immunophenotyping of plasmacytoid and myeloid dendritic cells.** Single cells (singlets) were selected by their FSC area (FSC-A) and height (FSC-H) patterns. Lymphocytes (LYM) were gated based on their characteristic forward and side scatter pattern (FSC, SSC). Dendritic cells (DC) were separated in two populations within the Lineage-DR+ ($HLA\text{-}DR^{+}$ $CD3^{-}$ $CD14^{-}$ $CD16^{-}$ $CD20^{-}$ $CD8^{-}$ $NKG2A^{-}$) by the expression of CD123 (plasmacytoid, pDC) or CD11c (myeloid, mDCs).

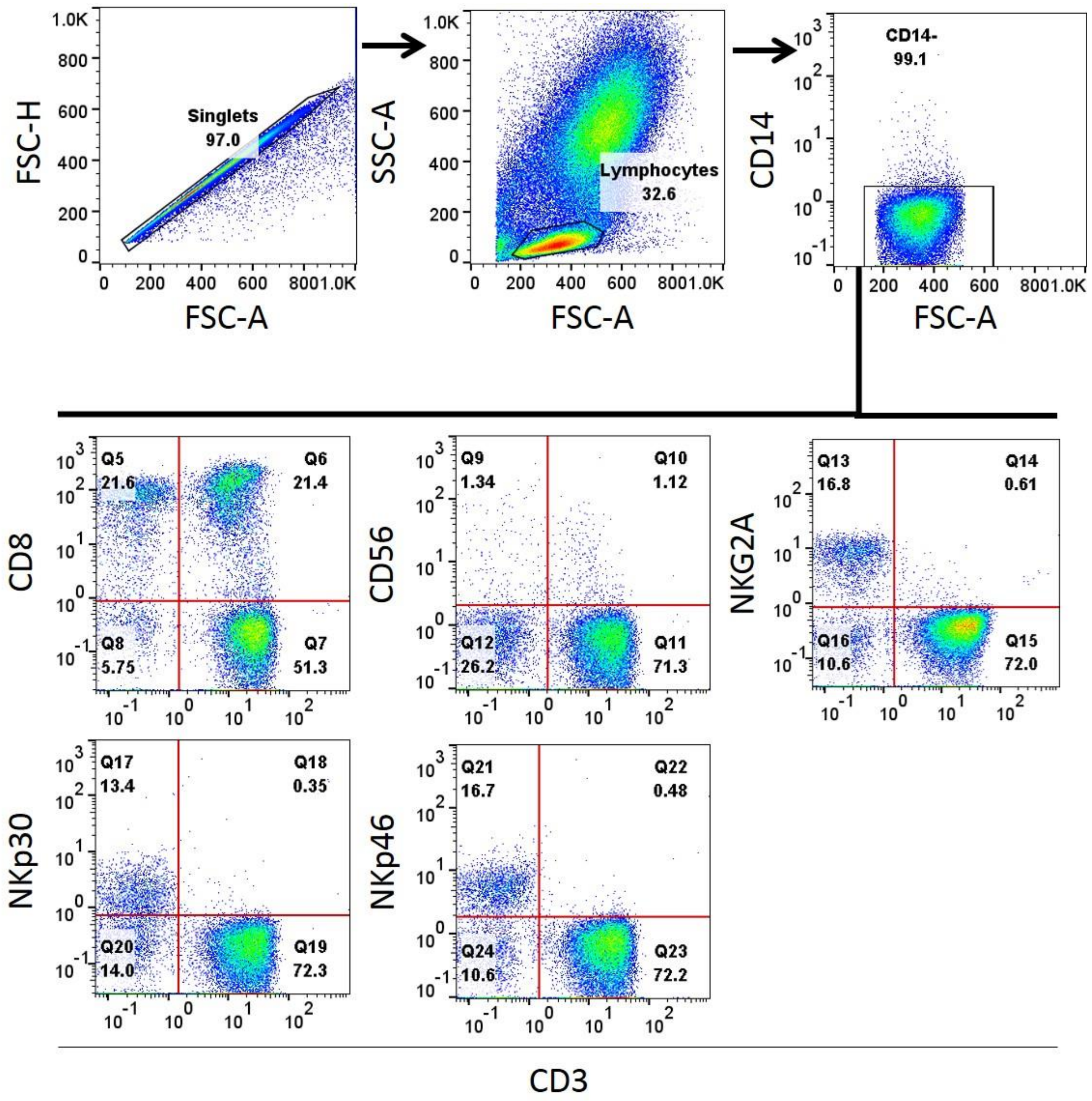


**Supplementary Figure 9 | Gating strategy for Natural killer cell subpopulations.** Single cells (singlets) were selected by their FSC area (FSC-A) and height (FSC-H) patterns. Lymphocytes (LYM) were gated based on their characteristic forward and side scatter pattern (FSC, SSC). Natural killer (NK) cells were defined as $CD3^{-}CD20^{-}CD14^{-}$ and analyzed by the double positive expression of the following NK cell markers: CD8, CD56, NKG2A, NKp30, and NKp46.

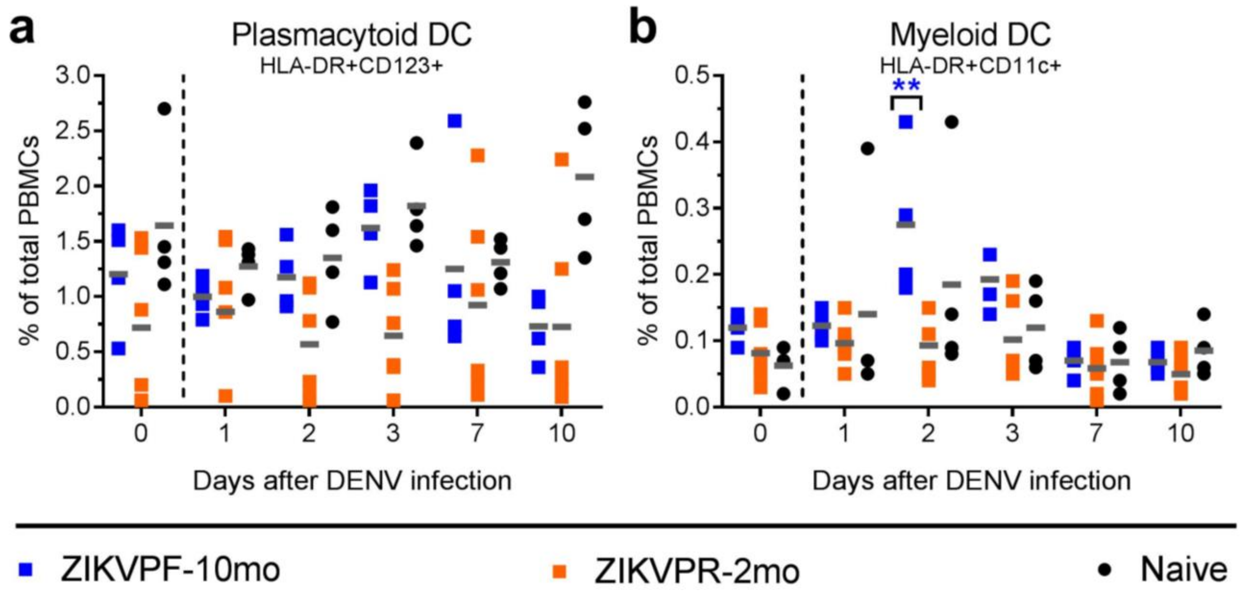


**Supplementary Figure 10 | Dendritic cells subsets modulation induced by DENV infection.** The frequency (% of total PBMCs) of dendritic cells (DCs) subsets including (**a**) plasmacytoid (pDCs: Lin-HLA-$DR^{+}CD123^{+}$) and (**b**) myeloid (mDCs: Lin-HLA-$DR^{+}CD11c^{+}$) was assessed before and up to 10 days after DENV infection by immunophenotyping using flow cytometry analysis. Symbols represent individual animals per group for each timepoint: blue squares (ZIKVPF-10mo), orange squares (ZIKVPR-2mo) and black circles (Naïve). Short gray lines depict mean value for each group detected overtime. Cutted line divide % of DCs quantified before and after DENV infection. Statistically significant differences within groups were determined using Two-Way Anova Dunnett's multiple comparisons test (comparison of each group response at each timepoint versus baseline of the same group) including 3 families, and 5 comparisons per family. Significant differences are reported as multiplicity adjusted *p* values (* <0.05, ** <0.01, *** <0.001, **** <0.0001). Asterisks represent significant difference between the corresponded timepoint and baseline within the same group.

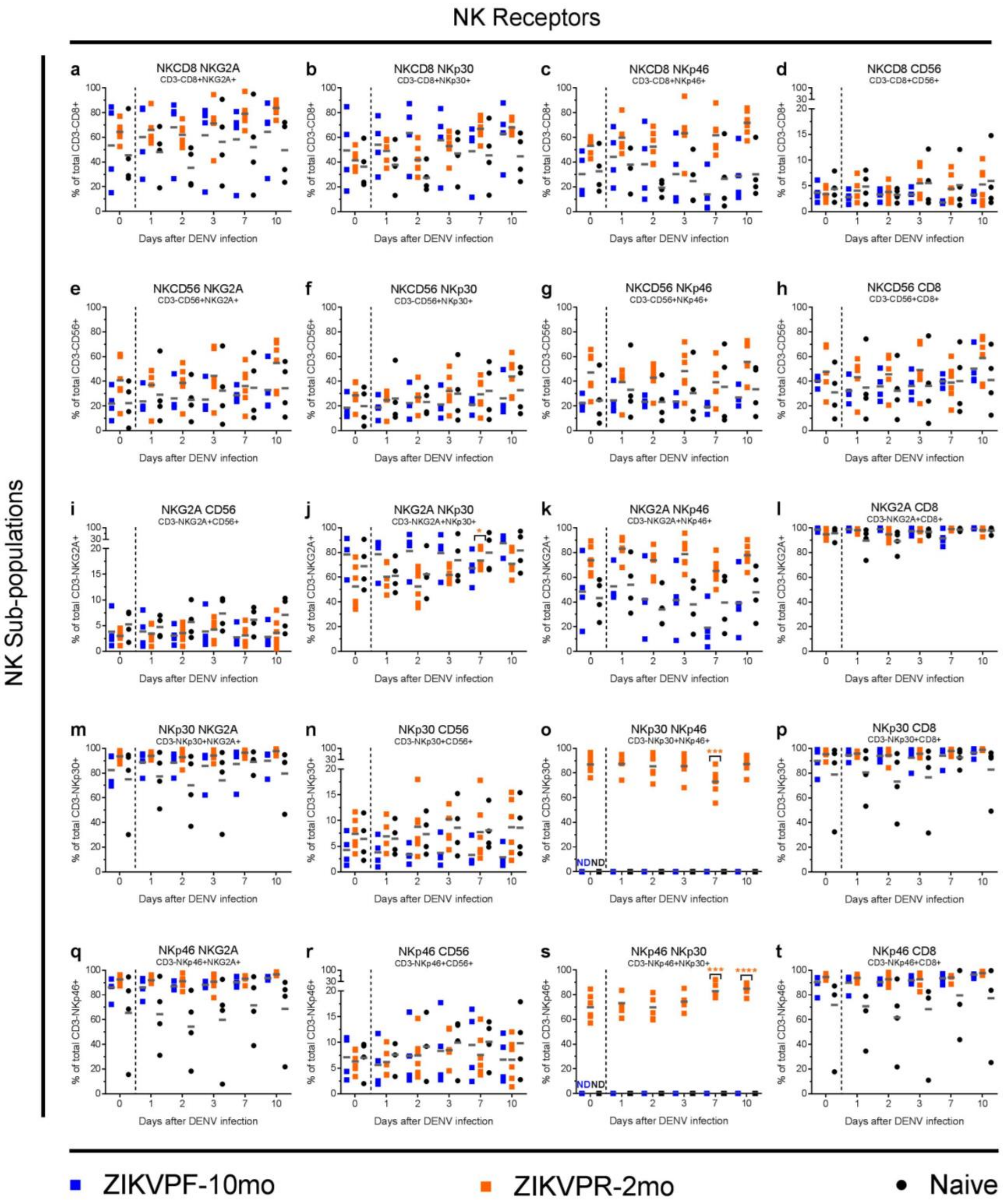


**Supplementary Figure 11 | Natural killer cell subpopulations and their differential receptors expression.** Natural killer (NK) cell subpopulations and the relative expression of multiple NK receptors within each subpopulation: (**a-d**) NKCD8, (**e-h**) NKCD56, (**i-l**) NKG2A, (**m-p**) NKp30 and (**q-t**) NKp46 were quantified by immunophenotyping using flow cytometry analysis before and up to 10 days after DENV infection. Individual symbols represent each animal per group over time: blue squares (ZIKVPF-10mo), orange squares (ZIKVPR-2mo) and black circles (Naïve). Short gray lines mark mean value for each group. Cutted line divide % of NK cells quantified before and after DENV infection. Statistically significant differences within groups were determined using Two-Way Anova Dunnett's multiple comparisons test (comparison of each group response at each timepoint versus baseline of the same group) including 3 families, and 5 comparisons per family. Significant differences are reported as multiplicity adjusted *p* values (* <0.05, ** <0.01, *** <0.001, **** <0.0001). Asterisks represent significant difference between the corresponded timepoint and baseline within the same group. ND (Not Done) in panels *8o* and *8s* refers that for ZIKVPF-10mo and Naïve groups the $NKp30^{+}NKp46^{+}$ and $NKp46^{+}NKp30^{+}$ subpopulations were not measured.

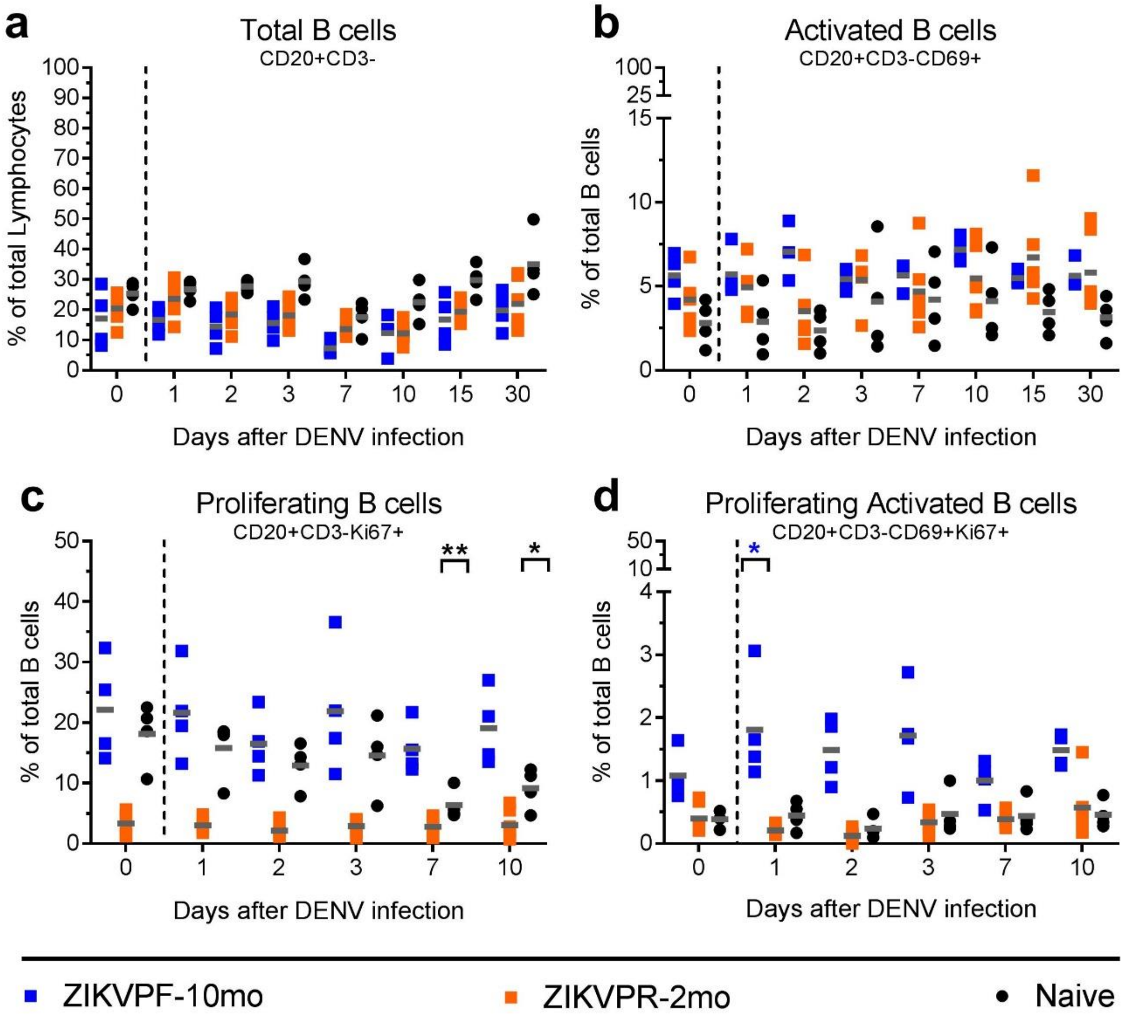


**Supplementary Figure 12 | B cells proliferation and activation higher in ZIKV middle-convalescent macaques.** The (**a**) total (% of total Lymphocytes), (**b**) activated, (**c**) proliferating, and (**d**) proliferating/activated B cells (% of total B cells) were determined at baseline and following DENV infection by immunophenotyping using flow cytometry analysis. B cells proliferation and activation were monitored since baseline up to 10 and 30 dpi, respectively. Symbols represent individual animals per group for each timepoint: blue squares (ZIKVPF-10mo), orange squares (ZIKVPR-2mo) and black circles (Naïve). Short gray lines depict mean value of B cells percent in each group of animals per timepoint. Cutted line divide % of B cells quantified before and after DENV infection. Statistically significant differences within groups were determined using Two-Way Anova Dunnett's multiple comparisons test (comparison of each group response at each timepoint versus baseline of the same group) including 3 families, and 7 and 5 comparisons per family in panels a-b and c-d, respectively. Significant differences are reported as multiplicity adjusted *p* values (* <0.05, ** <0.01). Asterisks represent significant difference between the corresponded timepoint and baseline within the same group.

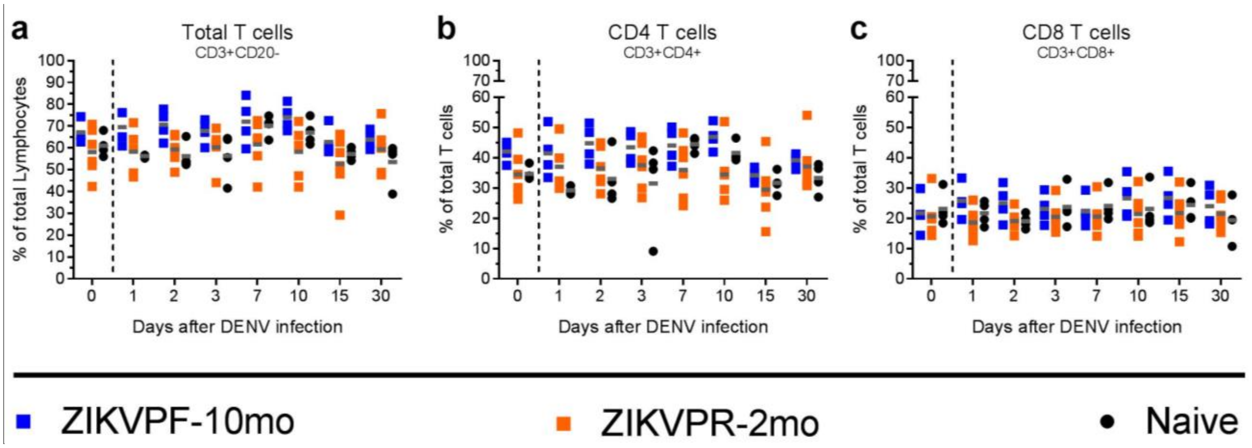


**Supplementary Figure 13 | Comparable T cells frequency between groups.** The (**a**) total T cells (% of total Lymphocytes), (**b**) $CD4^{+}$ and (**c**) $CD8^{+}$ T cell compartments (% of total T cells) frequencies were quantified at baseline and following DENV infection up to 30 dpi by immunophenotyping using flow cytometry. Symbols represent individual animals per group for each timepoint: blue squares (ZIKVPF-10mo), orange squares (ZIKVPR-2mo) and black circles (Naïve). Short gray lines mark mean value of T cells percent in each cohort per timepoint. Cutted line divide % of T cells quantified before and after DENV infection. Statistically significant differences within groups were determined using Two-Way Anova Dunnett's multiple comparisons test (comparison of each group response at each timepoint versus baseline of the same group) including 3 families, and 7 comparisons per family.

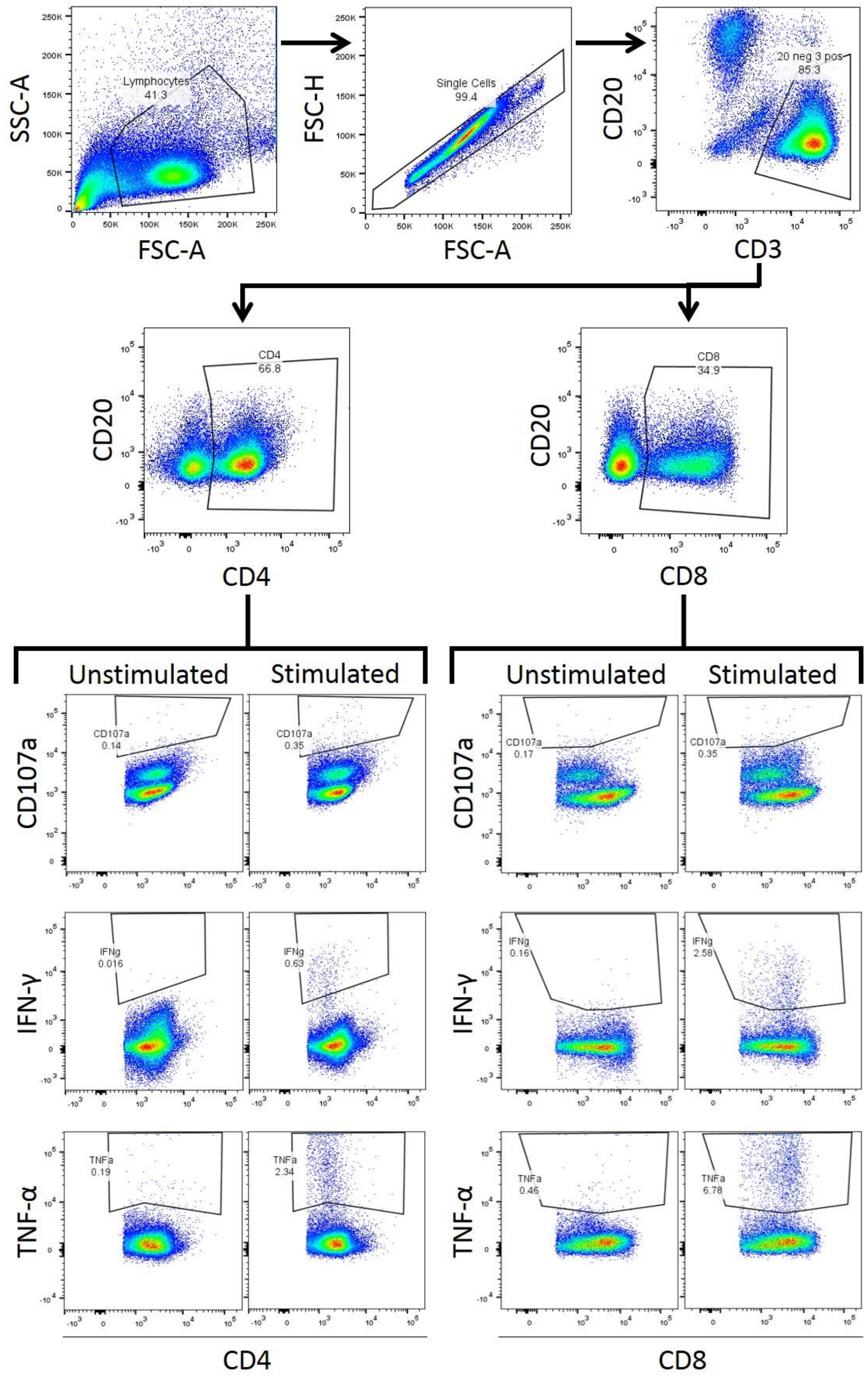
SSC-A
FSC-A
Lymphocytes
41.3
FSC-H
FSC-A
Single Cells
99.4
CD20
CD3
20 neg 3 pos
85.3
CD20
CD4
CD4
66.8
CD20
CD8
CD8
34.9
Unstimulated
Stimulated
Unstimulated
Stimulated
CD107a
CD107a
0.14
CD107a
0.35
CD107a
CD107a
0.17
CD107a
0.35
IFN-γ
IFNg
0.016
IFNg
0.63
IFN-γ
IFNg
0.16
IFNg
2.58
TNF-α
TNFa
0.19
TNFa
2.34
TNF-α
TNFa
0.46
TNFa
6.78
CD4
CD8

**Supplementary Figure 14 | Gating strategy for CD4+ and CD8+ T cell functional response.** After stimulation, lymphocytes (LYM) were gated based on their characteristic forward and side scatter pattern (FSC, SSC). Single cells (singlets) were selected by their FSC area (FSC-A) and height (FSC-H) patterns. Cells were stained for the following markers: CD3, CD4, CD8, CD20 (excluded), CD107a (functional cytotoxicity). Levels of IFN-γ and TNF-α also were measured in gated lymphocytes cell populations.

**Supplementary Table 1 | Sequence alignment and amino acid identity of ZIKV strains PRVABC59 and H/PF/2013.**

| | |
|---|---|
| Pairwise alignment of both ZIKV strains sequences.<br>(Sequences downloaded from ViPR database and global alignment was performed using Blosum62 in Genious Software). | >99.99% amino acid identity |
| Envelope (E) protein region of both ZIKV strains. | Identical |
| Amino acids residues changes between both ZIKV strains. Marked in red within sequences.<br><br>From ZIKV-PR → ZIKV-PF | $T_{80} \rightarrow$ I (Capsid)<br>$G_{892} \rightarrow$ W (NS1)<br>$V_{2611} \rightarrow$ A (NS5)<br>$V_{2634} \rightarrow$ M (NS5) |
| ZIKV-PRVABC59<br><br>Accession number: KX377337 | MKNPKKKSGGFRIVNMLKRGVARVSPFGGLKRLPAGLLLGHGPIRMVLAI<br>LAFLRFTAIKPSLGLINRWGSVGKKEAME**T**IKKFKKDLAAMLRIINARKE<br>KKRRGADTSVGIVGLLLTTAMAAEVTRRGSAYYMYLDRNDAGEAISFPTT<br>LGMNKCYIQIMDLGHMCDATMSYECPMLDEGVEPDDVDCWCNTTSTWVVY<br>GTCHHKKGEARRSRRAVTLPSHSTRKLQTRSQTWLESREYTKHLIRVENW<br>IFRNPGFALAAAAIAWLLGSSTSQKVIYLVMILLIAPAYSIRCIGVSNRD<br>FVEGMSGGTWVDVVLEHGGCVTVMAQDKPTVDIELVTTTVSNMAEVRSYC<br>YEASISDMASDSRCPTQGEAYLDKQSDTQYVCKRTLVDRGWGNGCGLFGK<br>GSLVTCAKFACSKKMTGKSIQPENLEYRIMLSVHGSQHSGMIVNDTGHET<br>DENRAKVEITPNSPRAEATLGGFGSLGLDCEPRTGLDFSDLYYLTMNNKH<br>WLVHKEWFHDIPLPWHAGADTGTPHWNNKEALVEFKDAHAKRQTVVVLGS<br>QEGAVHTALAGALEAEMDGAKGRLSSGHLKCRLKMDKLRLKGVSYSLCTA<br>AFTFTKIPAETLHGTVTVEVQYAGTDGPCKVPAQMAVDMQTLTPVGRLIT<br>ANPVITESTENSKMMLELDPPFGDSYIVIGVGEKKITHHWHRSGSTIGKA<br>FEATVRGAKRMAVLGDTAWDFGSVGGALNSLGKGIHQIFGAAFKSLFGGM<br>SWFSQILIGTLLMWLGLNTKNGSISLMCLALGGVLIFLSTAVSADVGCSV<br>DFSKKETRCGTGVFVYNDVEAWRDRYKYHPDSPRRLAAAVKQAWEDGICG<br>ISSVSRMENIMWRSVEGELNAILEENGVQLTVVVGSVKNPM**G**RGPQRLPV<br>PVNELPHGWKAWGKSYFVRAAKTNNSFVVDGDTLKECPLKHRAWNSFLVE<br>DHGFGVFHTSVWLKVREDYSLECDPAVIGTAVKGKEAVHSDLGYWIESEK<br>NDTWRLKRAHLIEMKTCEWPKSHTLWTDGIEESDLIIPKSLAGPLSHHNT<br>REGYRTQMKGPWHSEELEIRFEECPGTKVHVEETCGTRGPSLRSTTASGR<br>VIEEWCCRECTMPPLSFRAKDGCWYGMEIRPRKEPESNLVRSMVTAGSTD<br>HMDHFSLGVLVILLMVQEGLKKRMTTKIIISTSMAVLVAMILGGFSMSDL<br>AKLAILMGATFAEMNTGGDVAHLALIAAFKVRPALLVSFIFRANWTPRES<br>MLLALASCLLQTAISALEGDLMVLINGFALAWLAIRAMVVPRTDNITLAI<br>LAALTPLARGTLLVAWRAGLATCGGFMLLSLKGKGSVKKNLPFVMALGLT<br>AVRLVDPINVVGLLLLTRSGKRSWPPSEVLTAVGLICALAGGFAKADIEM<br>AGPMAAVGLLIVSYVVSGKSVDMYIERAGDITWEKDAEVTGNSPRLDVAL |

| | |
|---|---|
| | DESGDFSLVEDDGPPMREIILKVVLMTICGMNPIAIPFAAGAWYVYVKTG<br>KRSGALWDVPAPKEVKKGETTDGVYRVMTRRLLGSTQVGVGVMQEGVFHT<br>MWHVTKGSALRSGEGRLDPYWGDVKQDLVSYCGPWKLDAAWDGHSEVQLL<br>AVPPGERARNIQTLPGIFKTKDGDIGAVALDYPAGTSGSPILDKCGRVIG<br>LYGNGVVIKNGSYVSAITQGRREEETPVECFEPSMLKKKQLTVLDLHPGA<br>GKTRRVLPEIVREAIKTRLRTVILAPTRVVAAEMEEALRGLPVRYMTTAV<br>NVTHSGTEIVDLMCHATFTSRLLQPIRVPNYNLYIMDEAHFTDPSSIAAR<br>GYISTRVEMGEAAAIFMTATPPGTRDAFPDSNSPIMDTEVEVPERAWSSG<br>FDWVTDHSGKTVWFVPSVRNGNEIAACLTKAGKRVIQLSRKTFETEFQKT<br>KHQEWDFVVTTDISEMGANFKADRVIDSRRCLKPVILDGERVILAGPMPV<br>THASAAQRRGRIGRNPNKPGDEYLYGGGCAETDEDHAHWLEARMLLDNIY<br>LQDGLIASLYRPEADKVAAIEGEFKLRTEQRKTFVELMKRGDLPVWLAYQ<br>VASAGITYTDRRWCFDGTTNNTIMEDSVPAEVWTRHGEKRVLKPRWMDAR<br>VCSDHAALKSFKEFAAGKRGAAFGVMEALGTLPGHMTERFQEAIDNLAVL<br>MRAETGSRPYKAAAAQLPETLETIMLLGLLGTVSLGIFFVLMRNKGIGKM<br>GFGMVTLGASAWLMWLSEIEPARIACVLIVVFLLLVVLIPEPEKQRSPQD<br>NQMAIIIMVAVGLLGLITANELGWLERTKSDLSHLMGRREEGATIGFSMD<br>IDLRPASAWAIYAALTTFITPAVQHAVTTSYNNYSLMAMATQAGVLFGMG<br>KGMPFYAWDFGVPLLMIGCYSQLTPLTLIVAIILLVAHYMYLIPGLQAAA<br>ARAAQKRTAAGIMKNPVVDGIVVTDIDTMTIDPQVEKKMGQVLLIAVAVS<br>SAILSRTAWGWGEAGALITAATSTLWEGSPNKYWNSSTATSLCNIFRGSY<br>LAGASLIYTVTRNAGLVKRRGGGTGETLGEKWKARLNQMSALEFYSYKKS<br>GITEVCREEARRALKDGVATGGHAVSRGSAKLRWLVERGYLQPYGKVIDL<br>GCGRGGWSYY**V**ATIRKVQEVKGYTKGGPGHEEP**V**LVQSYGWNIVRLKSGV<br>DVFHMAAEPCDTLLCDIGESSSSPEVEEARTLRVLSMVGDWLEKRPGAFC<br>IKVLCPYTSTMMETLERLQRRYGGGLVRVPLSRNSTHEMYWVSGAKSNTI<br>KSVSTTSQLLLGRMDGPRRPVKYEEDVNLGSGTRAVVSCAEAPNMKIIGN<br>RIERIRSEHAETWFFDENHPYRTWAYHGSYEAPTQGSASSLINGVVRLLS<br>KPWDVVTGVTGIAMTDTTPYGQQRVFKEKVDTRVPDPQEGTRQVMSMVSS<br>WLWKELGKHKRPRVCTKEEFINKVRSNAALGAIFEEEKEWKTAVEAVNDP<br>RFWALVDKEREHHLRGECQSCVYNMMGKREKKQGEFGKAKGSRAIWYMWL<br>GARFLEFEALGFLNEDHWMGRENSGGGVEGLGLQRLGYVLEEMSRIPGGR<br>MYADDTAGWDTRISRFDLENEALITNQMEKGHRALALAIIKYTYQNKVVK<br>VLRPAEKGKTVMDIISRQDQRGSGQVVTYALNTFTNLVVQLIRNMEAEEV<br>LEMQDLWLLRRSEKVTNWLQSNGWDRLKRMAVSGDDCVVKPIDDRFAHAL<br>RFLNDMGKVRKDTQEWKPSTGWDNWEEVPFCSHHFNKLHLKDGRSIVVPC<br>RHQDELIGRARVSPGAGWSIRETACLAKSYAQMWQLLYFHRRDLRLMANA<br>ICSSVPVDWVPTGRTTWSIHGKGEWMTTEDMLVVWNRVWIEENDHMEDKT<br>PVTKWTDIPYLGKREDLWCGSLIGHRPRTTWAENIKNTVNMVRRIIGDEE<br>KYMDYLSTQVRYLGEEGSTPGVL |
| ZIKV H/PF2013<br><br>Accession number:<br>KJ776791 | MKNPKKKSGGFRIVNMLKRGVARVSPFGGLKRLPAGLLLGHGPIRMVLAI<br>LAFLRFTAIKPSLGLINRWGSVGKKEAME**I**IKKFKKDLAAMLRIINARKE<br>KKRRGADTSVGIVGLLLTTAMAAEVTRRGSAYYMYLDRNDAGEAISFPTT<br>LGMNKCYIQIMDLGHMCDATMSYECPMLDEGVEPDDVDCWCNTTSTWVVY<br>GTCHHKKGEARRSRRAVTLPSHSTRKLQTRSQTWLESREYTKHLIRVENW<br>IFRNPGFALAAAAIAWLLGSSTSQKVIYLVMILLIAPAYSIRCIGVSNRD<br>FVEGMSGGTWVDVVLEHGGCVTVMAQDKPTVDIELVTTTVSNMAEVRSYC<br>YEASISDMASDSRCPTQGEAYLDKQSDTQYVCKRTLVDRGWGNGCGLFGK<br>GSLVTCAKFACSKKMTGKSIQPENLEYRIMLSVHGSQHSGMIVNDTGHET<br>DENRAKVEITPNSPRAEATLGGFGSLGLDCEPRTGLDFSDLYYLTMNNKH<br>WLVHKEWFHDIPLPWHAGADTGTPHWNNKEALVEFKDAHAKRQTVVVLGS<br>QEGAVHTALAGALEAEMDGAKGRLSSGHLKCRLKMDKLRLKGVSYSLCTA<br>AFTFTKIPAETLHGTVTVEVQYAGTDGPCKVPAQMAVDMQTLTPVGRLIT<br>ANPVITESTENSKMMLELDPPFGDSYIVIGVGEKKITHHWHRSGSTIGKA<br>FEATVRGAKRMAVLGDTAWDFGSVGGALNSLGKGIHQIFGAAFKSLFGGM<br>SWFSQILIGTLLMWLGLNTKNGSISLMCLALGGVLIFLSTAVSADVGCSV<br>DFSKKETRCGTGVFVYNDVEAWRDRYKYHPDSPRRLAAAVKQAWEDGICG |

| | |
|---|---|
| | ISSVSRMENIMWRSVEGELNAILEENGVQLTVVVGSVKNPM**W**RGPQRLPV<br>PVNELPHGWKAWGKSYFVRAAKTNNSFVVDGDTLKECPLKHRAWNSFLVE<br>DHGFGVFHTSVWLKVREDYSLECDPAVIGTAVKGKEAVHSDLGYWIESEK<br>NDTWRLKRAHLIEMKTCEWPKSHTLWTDGIEESDLIIPKSLAGPLSHHNT<br>REGYRTQMKGPWHSEELEIRFEECPGTKVHVEETCGTRGPSLRSTTASGR<br>VIEEWCCRECTMPPLSFRAKDGCWYGMEIRPRKEPESNLVRSMVTAGSTD<br>HMDHFSLGVLVILLMVQEGLKKRMTTKIIISTSMAVLVAMILGGFSMSDL<br>AKLAILMGATFAEMNTGGDVAHLALIAAFKVRPALLVSFIFRANWTPRES<br>MLLALASCLLQTAISALEGDLMVLINGFALAWLAIRAMVVPRTDNITLAI<br>LAALTPLARGTLLVAWRAGLATCGGFMLLSLKGKGSVKKNLPFVMALGLT<br>AVRLVDPINVVGLLLLTRSGKRSWPPSEVLTAVGLICALAGGFAKADIEM<br>AGPMAAVGLLIVSYVVSGKSVDMYIERAGDITWEKDAEVTGNSPRLDVAL<br>DESGDFSLVEDDGPPMREIILKVVLMTICGMNPIAIPFAAGAWYVYVKTG<br>KRSGALWDVPAPKEVKKGETTDGVYRVMTRRLLGSTQVGVGVMQEGVFHT<br>MWHVTKGSALRSGEGRLDPYWGDVKQDLVSYCGPWKLDAAWDGHSEVQLL<br>AVPPGERARNIQTLPGIFKTKDGDIGAVALDYPAGTSGSPILDKCGRVIG<br>LYGNGVVIKNGSYVSAITQGRREEETPVECFEPSMLKKKQLTVLDLHPGA<br>GKTRRVLPEIVREAIKTRLRTVILAPTRVVAAEMEEALRGLPVRYMTTAV<br>NVTHSGTEIVDLMCHATFTSRLLQPIRVPNYNLYIMDEAHFTDPSSIAAR<br>GYISTRVEMGEAAAIFMTATPPGTRDAFPDSNSPIMDTEVEVPERAWSSG<br>FDWVTDHSGKTVWFVPSVRNGNEIAACLTKAGKRVIQLSRKTFETEFQKT<br>KHQEWDFVVTTDISEMGANFKADRVIDSRRCLKPVILDGERVILAGPMPV<br>THASAAQRRGRIGRNPNKPGDEYLYGGGCAETDEDHAHWLEARMLLDNIY<br>LQDGLIASLYRPEADKVAAIEGEFKLRTEQRKTFVELMKRGDLPVWLAYQ<br>VASAGITYTDRRWCFDGTTNNTIMEDSVPAEVWTRHGEKRVLKPRWMDAR<br>VCSDHAALKSFKEFAAGKRGAAFGVMEALGTLPGHMTERFQEAIDNLAVL<br>MRAETGSRPYKAAAAQLPETLETIMLLGLLGTVSLGIFFVLMRNKGIGKM<br>GFGMVTLGASAWLMWLSEIEPARIACVLIVVFLLLVVLIPEPEKQRSPQD<br>NQMAIIIMVAVGLLGLITANELGWLERTKSDLSHLMGRREEGATIGFSMD<br>IDLRPASAWAIYAALTTFITPAVQHAVTTSYNNYSLMAMATQAGVLFGMG<br>KGMPFYAWDFGVPLLMIGCYSQLTPLTLIVAIILLVAHYMYLIPGLQAAA<br>ARAAQKRTAAGIMKNPVVDGIVVTDIDTMTIDPQVEKKMGQVLLIAVAVS<br>SAILSRTAWGWGEAGALITAATSTLWEGSPNKYWNSSTATSLCNIFRGSY<br>LAGASLIYTVTRNAGLVKRRGGGTGETLGEKWKARLNQMSALEFYSYKKS<br>GITEVCREEARRALKDGVATGGHAVSRGSAKLRWLVERGYLQPYGKVIDL<br>GCGRGGWSYY**A**ATIRKVQEVKGYTKGGPGHEEP**M**LVQSYGWNIVRLKSGV<br>DVFHMAAEPCDTLLCDIGESSSSPEVEEARTLRVLSMVGDWLEKRPGAFC<br>IKVLCPYTSTMMETLERLQRRYGGGLVRVPLSRNSTHEMYWVSGAKSNTI<br>KSVSTTSQLLLGRMDGPRRPVKYEEDVNLGSGTRAVVSCAEAPNMKIIGN<br>RIERIRSEHAETWFFDENHPYRTWAYHGSYEAPTQGSASSLINGVVRLLS<br>KPWDVVTGVTGIAMTDTTPYGQQRVFKEKVDTRVPDPQEGTRQVMSMVSS<br>WLWKELGKHKRPRVCTKEEFINKVRSNAALGAIFEEEKEWKTAVEAVNDP<br>RFWALVDKEREHHLRGECQSCVYNMMGKREKKQGEFGKAKGSRAIWYMWL<br>GARFLEFEALGFLNEDHWMGRENSGGGVEGLGLQRLGYVLEEMSRIPGGR<br>MYADDTAGWDTRISRFDLENEALITNQMEKGHRALALAIIKYTYQNKVVK<br>VLRPAEKGKTVMDIISRQDQRGSGQVVTYALNTFTNLVVQLIRNMEAEEV<br>LEMQDLWLLRRSEKVTNWLQSNGWDRLKRMAVSGDDCVVKPIDDRFAHAL<br>RFLNDMGKVRKDTQEWKPSTGWDNWEEVPFCSHHFNKLHLKDGRSIVVPC<br>RHQDELIGRARVSPGAGWSIRETACLAKSYAQMWQLLYFHRRDLRLMANA<br>ICSSVPVDWVPTGRTTWSIHGKGEWMTTEDMLVVWNRVWIEENDHMEDKT<br>PVTKWTDIPYLGKREDLWCGSLIGHRPRTTWAENIKNTVNMVRRIIGDEE<br>KYMDYLSTQVRYLGEEGSTPGVL |

**Supplementary Table 2 | Antibody panel for Immunophenotyping.**

| Cell Subset | Ab | Clone | Dye | Company | Cat. # |
|---|---|---|---|---|---|
| B / T cells | CD20 | 2H7 | PacificBlue | BioLegend | 302328 |
| | CD3 | 10D12 | PE-Vio770 | Miltenyi | 130-104-202 |
| | CD4 | M-T466 | PerCP | Miltenyi | 130-101-147 |
| | CD8 | BW135/80 | VioGreen | Miltenyi | 130-096-902 |
| | CD28 | 15E8 | APC-Vio770 | Miltenyi | 130-104-278 |
| | CD69 | FN50 | PE | BD | 557050 |
| | CD95 | DX2 | APC | Miltenyi | 130-092-417 |
| | Ki67 | B56 | Alexa 488 | BD | 558616 |
| NK | CD3 | 10D12 | APC | Miltenyi | 130-091-998 |
| | CD16 | VEP13 | APC-Vio770 | Miltenyi | 130-096-655 |
| | CD56 | AF12-7H3 | PE | Miltenyi | 130-090-755 |
| | CD14 | M5E2 | V500 | BD | 561391 |
| | CD8 | SK1 | BV421 | BioLegend | 344748 |
| | NKp30 | AF29-4D12 | PE-Vio770 | Miltenyi | 130-104-116 |
| | NKp46 | BAB281 | PC5 | Beck-Coulter | A66902 |
| | NK2GA | REA110 | FITC | Miltenyi | 130-098-818 |
| DC | CD20 | 2H7 | FITC | BD | 555622 |
| | CD3 | SP34 | | BD | 556611 |
| | CD14 | M5E2 | | BD | 555397 |
| | CD16 | 3G8 | | BD | 555406 |
| | NKG2A | REA110 | | Miltenyi | 130-098-818 |
| | CD8 | SK1 | | BioLegend | 344704 |
| | HLA DR | REA 805 | VioGreen | Miltenyi | 130-111-795 |
| | CD123 | 7G3 | APC | BD | 560087 |
| | CD11c | 3.9 | PE/Cy7 | BioLegend | 301608 |

#### Supplementary Table 3 | Antibody panel for T cell functional response assessment.

| Marker | Stain | Clone | Catalog Number | Vendor | Dilution |
|---|---|---|---|---|---|
| CD4 | PerCP-Cy-5.5 | SK3 | 566316 | BD Biosciences | 1:25 |
| CD8β | PE | ECD | 6607123 | Beckman-Coulter | 1:20 |
| CD3 | Pacific Blue | SP34-2 | 558124 | BD Biosciences | 1:30 |
| CD20 | BV605 | 2H7 | 563783 | BD Biosciences | 1:30 |
| CD107a | FITC | H4A3 | 555800 | BD Biosciences | 1:10 |
| CD28 | PE-Cy-5 | CD28.2 | 555730 | BD Biosciences | 1:10 |
| CD95 | BV510 | DX2 | 305640 | Biolegend | 1:30 |
| IFN-γ | APC | B27 | 554702 | BD Biosciences | 1:30 |
| TNF-α | PE-Cy-7 | Mab11 | 557647 | BD Biosciences | 1:30 |

**Supplementary Table 4 | Peptide sequences for stimulation of T cell functional response.**

#### Dengue Virus Type 2 Peptides

| Peptide | Amino Acid Sequence | Peptide | Amino Acid Sequence | Peptide | Amino Acid Sequence |
|---|---|---|---|---|---|
| 1 | MRCIGISNRDFVEGV | 29 | AWLVHRQWFLDLPLPWL | 57 | MRGAKRMAILGDTAWDF |
| 2 | ISNRDFVEGVSGGSWVDI | 30 | WFLDLPLPWLPGADTQGSNW | 58 | AILGDTAWDFGSLGGVF |
| 3 | GVSGGSWVDIVLEHGSCV | 31 | PGADTQGSNWIQKETLV | 59 | WDFGSLGGVFTSIGKALH |
| 4 | DIVLEHGSCVTTMAKNK | 32 | SNWIQKETLVTFKNPHAK | 60 | VFTSIGKALHQVFGAIY |
| 5 | SCVTTMAKNKPTLDFELI | 33 | LVTFKNPHAKKQDVVVL | 61 | ALHQVFGAIYGAAFSGV |
| 6 | NKPTLDFELIETEAKQPA | 34 | HAKKQDVVVLGSQEGAMH | 62 | AIYGAAFSGVSWIMKILI |
| 7 | LIETEAKQPATLRKYCI | 35 | VLGSQEGAMHTALTGA | 63 | GVSWIMKILIGVIITWI |
| 8 | KQPATLRKYCIEAKL | 36 | GAMHTALTGATEIQM | 64 | ILIGVIITWIGMNSR |
| 9 | LRKYCIEAKLTNTTTDSR | 37 | ALTGATEIQMSSGNLLF | 65 | IITWIGMNSRSTSLSVSL |
| 10 | KLTNTTTDSRCPTQGEPSL | 38 | IQMSSGNLLFTGHLKCRL | 66 | SRSTSLSVSLVLVGVVTL |
| 11 | RCPTQGEPSLNEEQDKRF | 39 | LFTGHLKCRLRMDKLQLK | 67 | SLVLVGVVTLYLGVMVQA |
| 12 | SLNEEQDKRFVCKHSMV | 40 | RLRMDKLQLKGMSYSM | | |
| 13 | KRFVCKHSMVDRGWGNGCG L | 41 | LQLKGMSYSMCTGKFKVV | | |
| 14 | DRGWGNGCGLFGKGGIV | 42 | SMCTGKFKVVKEIAETQH | | |
| 15 | CGLFGKGGIVTCAMFTCK | 43 | VVKEIAETQHGTIVIRV | | |
| 16 | IVTCAMFTCKKNMKGKVV | 44 | TQHGTIVIRVQYEGDGSPCK | | |
| 17 | CKKNMKGKVVQPENLEY | 45 | VQYEGDGSPCKIPFEIM | | |
| 18 | KVVQPENLEYTIVITPH | 46 | SPCKIPFEIMDLEKRHVL | | |
| 19 | LEYTIVITPHSGEEHAV | 47 | IMDLEKRHVLGRLITV | | |
| 20 | TPHSGEEHAVGNDTGKH | 48 | RHVLGRLITVNPIVTEK | | |
| 21 | HAVGNDTGKHGKEIKI | 49 | ITVNPIVTEKDSPVNIEA | | |
| 22 | TGKHGKEIKITPQSSI | 50 | EKDSPVNIEAEPPFGDSY | | |
| 23 | EIKITPQSSITEAELTGY | 51 | EAEPPFGDSYIIIGV | | |
| 24 | SITEAELTGYGTVTM | 52 | FGDSYIIIGVEPGQLKL | | |
| 25 | ELTGYGTVTMECSPRTGL | 53 | IGVEPGQLKLNWFKK | | |
| 26 | TMECSPRTGLDFNEMVLL | 54 | GQLKLNWFKKGSSIGQMI | | |
| 27 | GLDFNEMVLLQMENKAWL | 55 | KKGSSIGQMIETTMRGAK | | |
| 28 | LLQMENKAWLVHRQWFL | 56 | MIETTMRGAKRMAIL | | |

#### Supplementary Table 4 | Continuation

#### Zika Virus Envelope Peptides

| Peptide | Amino Acid Sequence | Peptide | Amino Acid Sequence | Peptide | Amino Acid Sequence |
|---|---|---|---|---|---|
| ZIKV59 | IRCIGVSNRDFVEGM | ZIKV87 | LSVHGSQHSGMIVND | ZIKV115 | KGRLSSGHLKCRLKM |
| ZIKV60 | VSNRDFVEGMSGGTW | ZIKV88 | SQHSGMIVNDTGHET | ZIKV116 | SGHLKCRLKMDKLRL |
| ZIKV61 | FVEGMSGGTWVDVVL | ZIKV89 | MIVNDTGHETDENRA | ZIKV117 | CRLKMDKLRLKGVSY |
| ZIKV62 | SGGTWVDVVLEHGGC | ZIKV90 | TGHETDENRAKVEIT | ZIKV118 | DKLRLKGVSYSLCTA |
| ZIKV63 | VDVVLEHGGCVTVMA | ZIKV91 | DENRAKVEITPNSPR | ZIKV119 | KGVSYSLCTAAFTFT |
| ZIKV64 | EHGGCVTVMAQDKPT | ZIKV92 | KVEITPNSPRAEATL | ZIKV120 | SLCTAAFTFTKIPAE |
| ZIKV65 | VTVMAQDKPTVDIEL | ZIKV93 | PNSPRAEATLGGFGS | ZIKV121 | AFTFTKIPAETLHGT |
| ZIKV66 | QDKPTVDIELVTTTV | ZIKV94 | AEATLGGFGSLGLDC | ZIKV122 | KIPAETLHGTVTVEV |
| ZIKV67 | VDIELVTTTVSNMAE | ZIKV95 | GGFGSLGLDCEPRTG | ZIKV123 | TLHGTVTVEVQYAGT |
| ZIKV68 | VTTTVSNMAEVRSYC | ZIKV96 | LGLDCEPRTGLDFSD | ZIKV124 | VTVEVQYAGTDGPCK |
| ZIKV69 | SNMAEVRSYCYEASI | ZIKV97 | EPRTGLDFSDLYYLT | ZIKV125 | QYAGTDGPCKVPAQM |
| ZIKV70 | VRSYCYEASISDMAS | ZIKV98 | LDFSDLYYLTMNNKH | ZIKV126 | DGPCKVPAQMAVDMQ |
| ZIKV71 | YEASISDMASDSRCP | ZIKV99 | LYYLTMNNKHWLVHK | ZIKV127 | VPAQMAVDMQTLTPV |
| ZIKV72 | SDMASDSRCPTQGEA | ZIKV100 | MNNKHWLVHKEWFHD | ZIKV128 | AVDMQTLTPVGRLIT |
| ZIKV73 | DSRCPTQGEAYLDKQ | ZIKV101 | WLVHKEWFHDIPLPW | ZIKV129 | TLTPVGRLITANPVI |
| ZIKV74 | TQGEAYLDKQSDTQY | ZIKV102 | EWFHDIPLPWHAGAD | ZIKV130 | GRLITANPVITESTE |
| ZIKV75 | YLDKQSDTQYVCKRT | ZIKV103 | IPLPWHAGADTGTPH | ZIKV131 | ANPVITESTENSKMM |
| ZIKV76 | SDTQYVCKRTLVDRG | ZIKV104 | HAGADTGTPHWNNKE | ZIKV132 | TESTENSKMMLELDP |
| ZIKV77 | VCKRTLVDRGWGNGC | ZIKV105 | TGTPHWNNKEALVEF | ZIKV133 | NSKMMLELDPPFGDS |
| ZIKV78 | LVDRGWGNGCGLFGK | ZIKV106 | WNNKEALVEFKDAHA | ZIKV134 | LELDPPFGDSYIVIG |
| ZIKV79 | WGNGCGLFGKGSLVT | ZIKV107 | ALVEFKDAHAKRQTV | ZIKV135 | PFGDSYIVIGVGEKK |
| ZIKV80 | GLFGKGSLVTCAKFA | ZIKV108 | KDAHAKRQTVVVLGS | ZIKV136 | YIVIGVGEKKITHHW |
| ZIKV81 | GSLVTCAKFACSKKM | ZIKV109 | KRQTVVVLGSQEGAV | ZIKV137 | VGEKKITHHWHRSGS |
| ZIKV82 | CAKFACSKKMTGKSI | ZIKV110 | VVLGSQEGAVHTALA | ZIKV138 | ITHHWHRSGSTIGKA |
| ZIKV83 | CSKKMTGKSIQPENL | ZIKV111 | QEGAVHTALAGALEA | ZIKV139 | HRSGSTIGKAFEATV |
| ZIKV84 | TGKSIQPENLEYRIM | ZIKV112 | HTALAGALEAEMDGA | ZIKV140 | TIGKAFEATVRGAKR |
| ZIKV85 | QPENLEYRIMLSVHG | ZIKV113 | GALEAEMDGAKGRLS | ZIKV141 | FEATVRGAKRMAVLG |
| ZIKV86 | EYRIMLSVHGSQHSG | ZIKV114 | EMDGAKGRLSSGHLK | ZIKV142 | RGAKRMAVLGDTAWD |

#### Supplementary Table 4 | Continuation

| Peptide | Amino Acid Sequence |
| --- | --- |
| ZIKV143 | MAVLGDTAWDFGSVG |
| ZIKV144 | DTAWDFGSVGGALNS |
| ZIKV145 | FGSVGGALNSLGKGI |
| ZIKV146 | GALNSLGKGIHQIFG |
| ZIKV147 | LGKGIHQIFGAAFKS |
| ZIKV148 | HQIFGAAFKSLFGGM |
| ZIKV149 | AAFKSLFGGMSWFSQ |
| ZIKV150 | LFGGMSWFSQILIGT |
| ZIKV151 | SWFSQILIGTLLMWL |
| ZIKV152 | ILIGTLLMWLGLNTK |
| ZIKV153 | LLMWLGLNTKNGSIS |
| ZIKV154 | GLNTKNGSISLMCLA |
| ZIKV155 | NGSISLMCLALGGVL |
| ZIKV156 | LMCLALGGVLIFLST |
| ZIKV157 | LGGVLIFLSTAVSAD |
| ZIKV158 | IFLSTAVSADVGCSV |
| ZIKV159 | AVSADVGCSVDFSKK |

#### Supplementary Table 4 | Continuation

##### Zika Virus Non-Structural Peptides

| Peptide | Amino Acid Sequence | Peptide | Amino Acid Sequence | Peptide | Amino Acid Sequence |
|---|---|---|---|---|---|
| ZIKV160 | VGCSVDFSKKETRCG | ZIKV188 | ECPLKHRAWNSFLVE | ZIKV216 | GTKVHVEETCGTRGP |
| ZIKV161 | DFSKKETRCGTGVFV | ZIKV189 | HRAWNSFLVEDHGFG | ZIKV217 | VEETCGTRGPSLRST |
| ZIKV162 | ETRCGTGVFVYNDVE | ZIKV190 | SFLVEDHGFGVFHTS | ZIKV218 | GTRGPSLRSTTASGR |
| ZIKV163 | TGVFVYNDVEAWRDR | ZIKV191 | DHGFGVFHTSVWLKV | ZIKV219 | SLRSTTASGRVIEEW |
| ZIKV164 | YNDVEAWRDRYKYHP | ZIKV192 | VFHTSVWLKVREDYS | ZIKV220 | TASGRVIEEWCCREC |
| ZIKV165 | AWRDRYKYHPDSPRR | ZIKV193 | VWLKVREDYSLECDP | ZIKV221 | VIEEWCCRECTMPPL |
| ZIKV166 | YKYHPDSPRRLAAAV | ZIKV194 | REDYSLECDPAVIGT | ZIKV222 | CCRECTMPPLSFRAK |
| ZIKV167 | DSPRRLAAAVKQAWE | ZIKV195 | LECDPAVIGTAVKGK | ZIKV223 | TMPPLSFRAKDGCWY |
| ZIKV168 | LAAAVKQAWEDGICG | ZIKV196 | AVIGTAVKGKEAVHS | ZIKV224 | SFRAKDGCWYGMEIR |
| ZIKV169 | KQAWEDGICGISSVS | ZIKV197 | AVKGKEAVHSDLGYW | ZIKV225 | DGCWYGMEIRPRKEP |
| ZIKV170 | DGICGISSVSRMENI | ZIKV198 | EAVHSDLGYWIESEK | ZIKV226 | GMEIRPRKEPESNLV |
| ZIKV171 | ISSVSRMENIMWRSV | ZIKV199 | DLGYWIESEKNDTWR | ZIKV227 | PRKEPESNLVRSMVT |
| ZIKV172 | RMENIMWRSVEGELN | ZIKV200 | IESEKNDTWRLKRAH | ZIKV228 | ESNLVRSMVTAGSTD |
| ZIKV173 | MWRSVEGELNAILEE | ZIKV201 | NDTWRLKRAHLIEMK | ZIKV229 | RSMVTAGSTDHMDHF |
| ZIKV174 | EGELNAILEENGVQL | ZIKV202 | LKRAHLIEMKTCEWP | ZIKV230 | AGSTDHMDHFSLGVL |
| ZIKV175 | AILEENGVQLTVVVG | ZIKV203 | LIEMKTCEWPKSHTL | ZIKV231 | HMDHFSLGVLVILLM |
| ZIKV176 | NGVQLTVVVGSVKNP | ZIKV204 | TCEWPKSHTLWTDGI | ZIKV232 | SLGVLVILLMVQEGL |
| ZIKV177 | TVVVGSVKNPMWRGP | ZIKV205 | KSHTLWTDGIEESDL | ZIKV233 | VILLMVQEGLKKRMT |
| ZIKV178 | SVKNPMWRGPQRLPV | ZIKV206 | WTDGIEESDLIIPKS | ZIKV234 | VQEGLKKRMTTKIII |
| ZIKV179 | MWRGPQRLPVPVNEL | ZIKV207 | EESDLIIPKSLAGPL | ZIKV235 | KKRMTTKIIISTSMA |
| ZIKV180 | QRLPVPVNELPHGWK | ZIKV208 | IIPKSLAGPLSHHNT | ZIKV236 | TKIIISTSMAVLVAM |
| ZIKV181 | PVNELPHGWKAWGKS | ZIKV209 | LAGPLSHHNTREGYR | ZIKV237 | STSMAVLVAMILGGF |
| ZIKV182 | PHGWKAWGKSYFVRA | ZIKV210 | SHHNTREGYRTQMKG | ZIKV238 | VLVAMILGGFSMSDL |
| ZIKV183 | AWGKSYFVRAAKTNN | ZIKV211 | REGYRTQMKGPWHSE | ZIKV239 | ILGGFSMSDLAKLAI |
| ZIKV184 | YFVRAAKTNNSFVVD | ZIKV212 | TQMKGPWHSEELEIR | ZIKV240 | SMSDLAKLAILMGAT |
| ZIKV185 | AKTNNSFVVDGDTLK | ZIKV213 | PWHSEELEIRFEECP | ZIKV241 | AKLAILMGATFAEMN |
| ZIKV186 | SFVVDGDTLKECPLK | ZIKV214 | ELEIRFEECPGTKVH | ZIKV242 | LMGATFAEMNTGGDV |
| ZIKV187 | GDTLKECPLKHRAWN | ZIKV215 | FEECPGTKVHVEETC | ZIKV243 | FAEMNTGGDVAHLAL |

| Peptide | Amino Acid Sequence | Peptide | Amino Acid Sequence | Peptide | Amino Acid Sequence |
|---|---|---|---|---|---|
| ZIKV244 | TGGDVAHLALIAAFK | ZIKV272 | DPINVVGLLLLTRSG | ZIKV300 | YVKTGKRSGALWDVP |
| ZIKV245 | AHLALIAAFKVRPAL | ZIKV273 | VGLLLLTRSGKRSWP | ZIKV301 | KRSGALWDVPAPKEV |
| ZIKV246 | IAAFKVRPALLVSFI | ZIKV274 | LTRSGKRSWPPSEVL | ZIKV302 | LWDVPAPKEVKKGET |
| ZIKV247 | VRPALLVSFIFRANW | ZIKV275 | KRSWPPSEVLTAVGL | ZIKV303 | APKEVKKGETTDGVY |
| ZIKV248 | LVSFIFRANWTPRES | ZIKV276 | PSEVLTAVGLICALA | ZIKV304 | KKGETTDGVYRVMTR |
| ZIKV249 | FRANWTPRESMLLAL | ZIKV277 | TAVGLICALAGGFAK | ZIKV305 | TDGVYRVMTRRLLGS |
| ZIKV250 | TPRESMLLALASCLL | ZIKV278 | ICALAGGFAKADIEM | ZIKV306 | RVMTRRLLGSTQVGV |
| ZIKV251 | MLLALASCLLQTAIS | ZIKV279 | GGFAKADIEMAGPMA | ZIKV307 | RLLGSTQVGVGVMQE |
| ZIKV252 | ASCLLQTAISALEGD | ZIKV280 | ADIEMAGPMAAVGLL | ZIKV308 | TQVGVGVMQEGVFHT |
| ZIKV253 | QTAISALEGDLMVLI | ZIKV281 | AGPMAAVGLLIVSYV | ZIKV309 | GVMQEGVFHTMWHVT |
| ZIKV254 | ALEGDLMVLINGFAL | ZIKV282 | AVGLLIVSYVVSGKS | ZIKV310 | GVFHTMWHVTKGSAL |
| ZIKV255 | LMVLINGFALAWLAI | ZIKV283 | IVSYVVSGKSVDMYI | ZIKV311 | MWHVTKGSALRSGEG |
| ZIKV256 | NGFALAWLAIRAMVV | ZIKV284 | VSGKSVDMYIERAGD | ZIKV312 | KGSALRSGEGRLDPY |
| ZIKV257 | AWLAIRAMVVPRTDN | ZIKV285 | VDMYIERAGDITWEK | ZIKV313 | RSGEGRLDPYWGDVK |
| ZIKV258 | RAMVVPRTDNITLAI | ZIKV286 | ERAGDITWEKDAEVT | ZIKV314 | RLDPYWGDVKQDLVS |
| ZIKV259 | PRTDNITLAILAALT | ZIKV287 | ITWEKDAEVTGNSPR | ZIKV315 | WGDVKQDLVSYCGPW |
| ZIKV260 | ITLAILAALTPLARG | ZIKV288 | DAEVTGNSPRLDVAL | ZIKV316 | QDLVSYCGPWKLDAA |
| ZIKV261 | LAALTPLARGTLLVA | ZIKV289 | GNSPRLDVALDESGD | ZIKV317 | YCGPWKLDAAWDGHS |
| ZIKV262 | PLARGTLLVAWRAGL | ZIKV290 | LDVALDESGDFSLVE | ZIKV318 | KLDAAWDGHSEVQLL |
| ZIKV263 | TLLVAWRAGLATCGG | ZIKV291 | DESGDFSLVEDDGPP | ZIKV319 | WDGHSEVQLLAVPPG |
| ZIKV264 | WRAGLATCGGFMLLS | ZIKV292 | FSLVEDDGPPMREII | ZIKV320 | EVQLLAVPPGERARN |
| ZIKV265 | ATCGGFMLLSLKGKG | ZIKV293 | DDGPPMREIILKVVL | ZIKV321 | AVPPGERARNIQTLP |
| ZIKV266 | FMLLSLKGKGSVKKN | ZIKV294 | MREIILKVVLMTICG | ZIKV322 | ERARNIQTLPGIFKT |
| ZIKV267 | LKGKGSVKKNLPFVM | ZIKV295 | LKVVLMTICGMNPIA | ZIKV323 | IQTLPGIFKTKDGDI |
| ZIKV268 | SVKKNLPFVMALGLT | ZIKV296 | MTICGMNPIAIPFAA | ZIKV324 | GIFKTKDGDIGAVAL |
| ZIKV269 | LPFVMALGLTAVRLV | ZIKV297 | MNPIAIPFAAGAWYV | ZIKV325 | KDGDIGAVALDYPAG |
| ZIKV270 | ALGLTAVRLVDPINV | ZIKV298 | IPFAAGAWYVYVKTG | ZIKV326 | GAVALDYPAGTSGSP |
| ZIKV271 | AVRLVDPINVVGLLL | ZIKV299 | GAWYVYVKTGKRSGA | ZIKV327 | DYPAGTSGSPILDKC |

| Peptide | Amino Acid Sequence | Peptide | Amino Acid Sequence | Peptide | Amino Acid Sequence |
|---|---|---|---|---|---|
| ZIKV328 | TSGSPILDKCGRVIG | ZIKV356 | IRVPNYNLYIMDEAH | ZIKV384 | MGANFKADRVIDSRR |
| ZIKV329 | ILDKCGRVIGLYGNG | ZIKV357 | YNLYIMDEAHFTDPS | ZIKV385 | KADRVIDSRRCLKPV |
| ZIKV330 | GRVIGLYGNGVVIKN | ZIKV358 | MDEAHFTDPSSIAAR | ZIKV386 | IDSRRCLKPVILDGE |
| ZIKV331 | LYGNGVVIKNGSYVS | ZIKV359 | FTDPSSIAARGYIST | ZIKV387 | CLKPVILDGERVILA |
| ZIKV332 | VVIKNGSYVSAITQG | ZIKV360 | SIAARGYISTRVEMG | ZIKV388 | ILDGERVILAGPMPV |
| ZIKV333 | GSYVSAITQGRREEE | ZIKV361 | GYISTRVEMGEAAAI | ZIKV389 | RVILAGPMPVTHASA |
| ZIKV334 | AITQGRREEETPVEC | ZIKV362 | RVEMGEAAAIFMTAT | ZIKV390 | GPMPVTHASAAQRRG |
| ZIKV335 | RREEETPVECFEPSM | ZIKV363 | EAAAIFMTATPPGTR | ZIKV391 | THASAAQRRGRIGRN |
| ZIKV336 | TPVECFEPSMLKKKQ | ZIKV364 | FMTATPPGTRDAFPD | ZIKV392 | AQRRGRIGRNPNKPG |
| ZIKV337 | FEPSMLKKKQLTVLD | ZIKV365 | PPGTRDAFPDSNSPI | ZIKV393 | RIGRNPNKPGDEYLY |
| ZIKV338 | LKKKQLTVLDLHPGA | ZIKV366 | DAFPDSNSPIMDTEV | ZIKV394 | PNKPGDEYLYGGGCA |
| ZIKV339 | LTVLDLHPGAGKTRR | ZIKV367 | SNSPIMDTEVEVPER | ZIKV395 | DEYLYGGGCAETDED |
| ZIKV340 | LHPGAGKTRRVLPEI | ZIKV368 | MDTEVEVPERAWSSG | ZIKV396 | GGGCAETDEDHAHWL |
| ZIKV341 | GKTRRVLPEIVREAI | ZIKV369 | EVPERAWSSGFDWVT | ZIKV397 | ETDEDHAHWLEARML |
| ZIKV342 | VLPEIVREAIKTRLR | ZIKV370 | AWSSGFDWVTDHSGK | ZIKV398 | HAHWLEARMLLDNIY |
| ZIKV343 | VREAIKTRLRTVILA | ZIKV371 | FDWVTDHSGKTVWFV | ZIKV399 | EARMLLDNIYLQDGL |
| ZIKV344 | KTRLRTVILAPTRVV | ZIKV372 | DHSGKTVWFVPSVRN | ZIKV400 | LDNIYLQDGLIASLY |
| ZIKV345 | TVILAPTRVVAAEME | ZIKV373 | TVWFVPSVRNGNEIA | ZIKV401 | LQDGLIASLYRPEAD |
| ZIKV346 | PTRVVAAEMEEALRG | ZIKV374 | PSVRNGNEIAACLTK | ZIKV402 | IASLYRPEADKVAAI |
| ZIKV347 | AAEMEEALRGLPVRY | ZIKV375 | GNEIAACLTKAGKRV | ZIKV403 | RPEADKVAAIEGEFK |
| ZIKV348 | EALRGLPVRYMTTAV | ZIKV376 | ACLTKAGKRVIQLSR | ZIKV404 | KVAAIEGEFKLRTEQ |
| ZIKV349 | LPVRYMTTAVNVTHS | ZIKV377 | AGKRVIQLSRKTFET | ZIKV405 | EGEFKLRTEQRKTFV |
| ZIKV350 | MTTAVNVTHSGTEIV | ZIKV378 | IQLSRKTFETEFQKT | ZIKV406 | LRTEQRKTFVELMKR |
| ZIKV351 | NVTHSGTEIVDLMCH | ZIKV379 | KTFETEFQKTKHQEW | ZIKV407 | RKTFVELMKRGDLPV |
| ZIKV352 | GTEIVDLMCHATFTS | ZIKV380 | EFQKTKHQEWDFVVT | ZIKV408 | ELMKRGDLPVWLAYQ |
| ZIKV353 | DLMCHATFTSRLLQP | ZIKV381 | KHQEWDFVVTTDISE | ZIKV409 | GDLPVWLAYQVASAG |
| ZIKV354 | ATFTSRLLQPIRVPN | ZIKV382 | DFVVTTDISEMGANF | ZIKV410 | WLAYQVASAGITYTD |
| ZIKV355 | RLLQPIRVPNYNLYI | ZIKV383 | TDISEMGANFKADRV | ZIKV411 | VASAGITYTDRRWCF |

| Peptide | Amino Acid Sequence | Peptide | Amino Acid Sequence | Peptide | Amino Acid Sequence |
|---|---|---|---|---|---|
| ZIKV412 | ITYTDRRWCFDGTTN | ZIKV440 | GIGKMGFGMVTLGAS | ZIKV468 | LMAMATQAGVLFGMG |
| ZIKV413 | RRWCFDGTTNNTIME | ZIKV441 | GFGMVTLGASAWLMW | ZIKV469 | TQAGVLFGMGKGMPF |
| ZIKV414 | DGTTNNTIMEDSVPA | ZIKV442 | TLGASAWLMWLSEIE | ZIKV470 | LFGMGKGMPFYAWDF |
| ZIKV415 | NTIMEDSVPAEVWTR | ZIKV443 | AWLMWLSEIEPARIA | ZIKV471 | KGMPFYAWDFGVPLL |
| ZIKV416 | DSVPAEVWTRHGEKR | ZIKV444 | LSEIEPARIACVLIV | ZIKV472 | YAWDFGVPLLMIGCY |
| ZIKV417 | EVWTRHGEKRVLKPR | ZIKV445 | PARIACVLIVVFLLL | ZIKV473 | GVPLLMIGCYSQLTP |
| ZIKV418 | HGEKRVLKPRWMDAR | ZIKV446 | CVLIVVFLLLVVLIP | ZIKV474 | MIGCYSQLTPLTLIV |
| ZIKV419 | VLKPRWMDARVCSDH | ZIKV447 | VFLLLVVLIPEPEKQ | ZIKV475 | SQLTPLTLIVAIILL |
| ZIKV420 | WMDARVCSDHAALKS | ZIKV448 | VVLIPEPEKQRSPQD | ZIKV476 | LTLIVAIILLVAHYM |
| ZIKV421 | VCSDHAALKSFKEFA | ZIKV449 | EPEKQRSPQDNQMAI | ZIKV477 | AIILLVAHYMYLIPG |
| ZIKV422 | AALKSFKEFAAGKRG | ZIKV450 | RSPQDNQMAIIIMVA | ZIKV478 | VAHYMYLIPGLQAAA |
| ZIKV423 | FKEFAAGKRGAAFGV | ZIKV451 | NQMAIIIMVAVGLLG | ZIKV479 | YLIPGLQAAAARAAQ |
| ZIKV424 | AGKRGAAFGVMEALG | ZIKV452 | IIMVAVGLLGLITAN | ZIKV480 | LQAAAARAAQKRTAA |
| ZIKV425 | AAFGVMEALGTLPGH | ZIKV453 | VGLLGLITANELGWL | ZIKV481 | ARAAQKRTAAGIMKN |
| ZIKV426 | MEALGTLPGHMTERF | ZIKV454 | LITANELGWLERTKS | ZIKV482 | KRTAAGIMKNPVVDG |
| ZIKV427 | TLPGHMTERFQEAID | ZIKV455 | ELGWLERTKSDLSHL | ZIKV483 | GIMKNPVVDGIVVTD |
| ZIKV428 | MTERFQEAIDNLAVL | ZIKV456 | ERTKSDLSHLMGRRE | ZIKV484 | PVVDGIVVTDIDTMT |
| ZIKV429 | QEAIDNLAVLMRAET | ZIKV457 | DLSHLMGRREEGATI | ZIKV485 | IVVTDIDTMTIDPQV |
| ZIKV430 | NLAVLMRAETGSRPY | ZIKV458 | MGRREEGATIGFSMD | ZIKV486 | IDTMTIDPQVEKKMG |
| ZIKV431 | MRAETGSRPYKAAAA | ZIKV459 | EGATIGFSMDIDLRP | ZIKV487 | IDPQVEKKMGQVLLI |
| ZIKV432 | GSRPYKAAAAQLPET | ZIKV460 | GFSMDIDLRPASAWA | ZIKV488 | EKKMGQVLLIAVAVS |
| ZIKV433 | KAAAAQLPETLETIM | ZIKV461 | IDLRPASAWAIYAAL | ZIKV489 | QVLLIAVAVSSAILS |
| ZIKV434 | QLPETLETIMLLGLL | ZIKV462 | ASAWAIYAALTTFIT | ZIKV490 | AVAVSSAILSRTAWG |
| ZIKV435 | LETIMLLGLLGTVSL | ZIKV463 | IYAALTTFITPAVQH | ZIKV491 | SAILSRTAWGWGEAG |
| ZIKV436 | LLGLLGTVSLGIFFV | ZIKV464 | TTFITPAVQHAVTTS | ZIKV492 | RTAWGWGEAGALITA |
| ZIKV437 | GTVSLGIFFVLMRNK | ZIKV465 | PAVQHAVTTSYNNYS | ZIKV493 | WGEAGALITAATSTL |
| ZIKV438 | GIFFVLMRNKGIGKM | ZIKV466 | AVTTSYNNYSLMAMA | ZIKV494 | ALITAATSTLWEGSP |
| ZIKV439 | LMRNKGIGKMGFGMV | ZIKV467 | YNNYSLMAMATQAGV | ZIKV495 | ATSTLWEGSPNKYWN |

| Peptide | Amino Acid Sequence | Peptide | Amino Acid Sequence | Peptide | Amino Acid Sequence |
|---|---|---|---|---|---|
| ZIKV496 | WEGSPNKYWNSSTAT | ZIKV524 | KVQEVKGYTKGGPGH | ZIKV552 | TSQLLLGRMDGPRRP |
| ZIKV497 | NKYWNSSTATSLCNI | ZIKV525 | KGYTKGGPGHEEPVL | ZIKV553 | LGRMDGPRRPVKYEE |
| ZIKV498 | SSTATSLCNIFRGSY | ZIKV526 | GGPGHEEPVLVQSYG | ZIKV554 | GPRRPVKYEEDVNLG |
| ZIKV499 | SLCNIFRGSYLAGAS | ZIKV527 | EEPVLVQSYGWNIVR | ZIKV555 | VKYEEDVNLGSGTRA |
| ZIKV500 | FRGSYLAGASLIYTV | ZIKV528 | VQSYGWNIVRLKSGV | ZIKV556 | DVNLGSGTRAVVSCA |
| ZIKV501 | LAGASLIYTVTRNAG | ZIKV529 | WNIVRLKSGVDVFHM | ZIKV557 | SGTRAVVSCAEAPNM |
| ZIKV502 | LIYTVTRNAGLVKRR | ZIKV530 | LKSGVDVFHMAAEPC | ZIKV558 | VVSCAEAPNMKIIGN |
| ZIKV503 | TRNAGLVKRRGGGTG | ZIKV531 | DVFHMAAEPCDTLLC | ZIKV559 | EAPNMKIIGNRIERI |
| ZIKV504 | LVKRRGGGTGETLGE | ZIKV532 | AAEPCDTLLCDIGES | ZIKV560 | KIIGNRIERIRSEHA |
| ZIKV505 | GGGTGETLGEKWKAR | ZIKV533 | DTLLCDIGESSSSPE | ZIKV561 | RIERIRSEHAETWFF |
| ZIKV506 | ETLGEKWKARLNQMS | ZIKV534 | DIGESSSSPEVEEAR | ZIKV562 | RSEHAETWFFDENHP |
| ZIKV507 | KWKARLNQMSALEFY | ZIKV535 | SSSPEVEEARTLRVL | ZIKV563 | ETWFFDENHPYRTWA |
| ZIKV508 | LNQMSALEFYSYKKS | ZIKV536 | VEEARTLRVLSMVGD | ZIKV564 | DENHPYRTWAYHGSY |
| ZIKV509 | ALEFYSYKKSGITEV | ZIKV537 | TLRVLSMVGDWLEKR | ZIKV565 | YRTWAYHGSYEAPTQ |
| ZIKV510 | SYKKSGITEVCREEA | ZIKV538 | SMVGDWLEKRPGAFC | ZIKV566 | YHGSYEAPTQGSASS |
| ZIKV511 | GITEVCREEARRALK | ZIKV539 | WLEKRPGAFCIKVLC | ZIKV567 | EAPTQGSASSLINGV |
| ZIKV512 | CREEARRALKDGVAT | ZIKV540 | PGAFCIKVLCPYTST | ZIKV568 | GSASSLINGVVRLLS |
| ZIKV513 | RRALKDGVATGGHAV | ZIKV541 | IKVLCPYTSTMMETL | ZIKV569 | LINGVVRLLSKPWDV |
| ZIKV514 | DGVATGGHAVSRGSA | ZIKV542 | PYTSTMMETLERLQR | ZIKV570 | VRLLSKPWDVVTGVT |
| ZIKV515 | GGHAVSRGSAKLRWL | ZIKV543 | MMETLERLQRRYGGG | ZIKV571 | KPWDVVTGVTGIAMT |
| ZIKV516 | SRGSAKLRWLVERGY | ZIKV544 | ERLQRRYGGGLVRVP | ZIKV572 | VTGVTGIAMTDTTPY |
| ZIKV517 | KLRWLVERGYLQPYG | ZIKV545 | RYGGGLVRVPLSRNS | ZIKV573 | GIAMTDTTPYGQQRV |
| ZIKV518 | VERGYLQPYGKVIDL | ZIKV546 | LVRVPLSRNSTHEMY | ZIKV574 | DTTPYGQQRVFKEKV |
| ZIKV519 | LQPYGKVIDLGCGRG | ZIKV547 | LSRNSTHEMYWVSGA | ZIKV575 | GQQRVFKEKVDTRVP |
| ZIKV520 | KVIDLGCGRGGWSYY | ZIKV548 | THEMYWVSGAKSNTI | ZIKV576 | FKEKVDTRVPDPQEG |
| ZIKV521 | GCGRGGWSYYVATIR | ZIKV549 | WVSGAKSNTIKSVST | ZIKV577 | DTRVPDPQEGTRQVM |
| ZIKV522 | GWSYYVATIRKVQEV | ZIKV550 | KSNTIKSVSTTSQLL | ZIKV578 | DPQEGTRQVMSMVSS |
| ZIKV523 | VATIRKVQEVKGYTK | ZIKV551 | KSVSTTSQLLLGRMD | ZIKV579 | TRQVMSMVSSWLWKE |

| Peptide | Amino Acid Sequence | Peptide | Amino Acid Sequence | Peptide | Amino Acid Sequence |
|---|---|---|---|---|---|
| ZIKV580 | SMVSSWLWKELGKHK | ZIKV608 | LGYVLEEMSRIPGGR | ZIKV636 | RLKRMAVSGDDCVVK |
| ZIKV581 | WLWKELGKHKRPRVC | ZIKV609 | EEMSRIPGGRMYADD | ZIKV637 | AVSGDDCVVKPIDDR |
| ZIKV582 | LGKHKRPRVCTKEEF | ZIKV610 | IPGGRMYADDTAGWD | ZIKV638 | DCVVKPIDDRFAHAL |
| ZIKV583 | RPRVCTKEEFINKVR | ZIKV611 | MYADDTAGWDTRISR | ZIKV639 | PIDDRFAHALRFLND |
| ZIKV584 | TKEEFINKVRSNAAL | ZIKV612 | TAGWDTRISRFDLEN | ZIKV640 | FAHALRFLNDMGKVR |
| ZIKV585 | INKVRSNAALGAIFE | ZIKV613 | TRISRFDLENEALIT | ZIKV641 | RFLNDMGKVRKDTQE |
| ZIKV586 | SNAALGAIFEEEKEW | ZIKV614 | FDLENEALITNQMEK | ZIKV642 | MGKVRKDTQEWKPST |
| ZIKV587 | GAIFEEEKEWKTAVE | ZIKV615 | EALITNQMEKGHRAL | ZIKV643 | KDTQEWKPSTGWDNW |
| ZIKV588 | EEKEWKTAVEAVNDP | ZIKV616 | NQMEKGHRALALAII | ZIKV644 | WKPSTGWDNWEEVPF |
| ZIKV589 | KTAVEAVNDPRFWAL | ZIKV617 | GHRALALAIIKYTYQ | ZIKV645 | GWDNWEEVPFCSHHF |
| ZIKV590 | AVNDPRFWALVDKER | ZIKV618 | ALAIIKYTYQNKVVK | ZIKV646 | EEVPFCSHHFNKLHL |
| ZIKV591 | RFWALVDKEREHHLR | ZIKV619 | KYTYQNKVVKVLRPA | ZIKV647 | CSHHFNKLHLKDGRS |
| ZIKV592 | VDKEREHHLRGECQS | ZIKV620 | NKVVKVLRPAEKGKT | ZIKV648 | NKLHLKDGRSIVVPC |
| ZIKV593 | EHHLRGECQSCVYNM | ZIKV621 | VLRPAEKGKTVMDII | ZIKV649 | KDGRSIVVPCRHQDE |
| ZIKV594 | GECQSCVYNMMGKRE | ZIKV622 | EKGKTVMDIISRQDQ | ZIKV650 | IVVPCRHQDELIGRA |
| ZIKV595 | CVYNMMGKREKKQGE | ZIKV623 | VMDIISRQDQRGSGQ | ZIKV651 | RHQDELIGRARVSPG |
| ZIKV596 | MGKREKKQGEFGKAK | ZIKV624 | SRQDQRGSGQVVTYA | ZIKV652 | LIGRARVSPGAGWSI |
| ZIKV597 | KKQGEFGKAKGSRAI | ZIKV625 | RGSGQVVTYALNTFT | ZIKV653 | RVSPGAGWSIRETAC |
| ZIKV598 | FGKAKGSRAIWYMWL | ZIKV626 | VVTYALNTFTNLVVQ | ZIKV654 | AGWSIRETACLAKSY |
| ZIKV599 | GSRAIWYMWLGARFL | ZIKV627 | LNTFTNLVVQLIRNM | ZIKV655 | RETACLAKSYAQMWQ |
| ZIKV600 | WYMWLGARFLEFEAL | ZIKV628 | NLVVQLIRNMEAEEV | ZIKV656 | LAKSYAQMWQLLYFH |
| ZIKV601 | GARFLEFEALGFLNE | ZIKV629 | LIRNMEAEEVLEMQD | ZIKV657 | AQMWQLLYFHRRDLR |
| ZIKV602 | EFEALGFLNEDHWMG | ZIKV630 | EAEEVLEMQDLWLLR | ZIKV658 | LLYFHRRDLRLMANA |
| ZIKV603 | GFLNEDHWMGRENSG | ZIKV631 | LEMQDLWLLRRSEKV | ZIKV659 | RRDLRLMANAICSSV |
| ZIKV604 | DHWMGRENSGGGVEG | ZIKV632 | LWLLRRSEKVTNWLQ | ZIKV660 | LMANAICSSVPVDWV |
| ZIKV605 | RENSGGGVEGLGLQR | ZIKV633 | RSEKVTNWLQSNGWD | ZIKV661 | ICSSVPVDWVPTGRT |
| ZIKV606 | GGVEGLGLQRLGYVL | ZIKV634 | TNWLQSNGWDRLKRM | ZIKV662 | PVDWVPTGRTTWSIH |
| ZIKV607 | LGLQRLGYVLEEMSR | ZIKV635 | SNGWDRLKRMAVSGD | ZIKV663 | PTGRTTWSIHGKGEW |

| Peptide | Amino Acid Sequence |
| --- | --- |
| ZIKV664 | TWSIHGKGEWMTTED |
| ZIKV665 | GKGEWMTTEDMLVVW |
| ZIKV666 | MTTEDMLVVWNRVWI |
| ZIKV667 | MLVVWNRVWIEENDH |
| ZIKV668 | NRVWIEENDHMEDKT |
| ZIKV669 | EENDHMEDKTPVTKW |
| ZIKV670 | MEDKTPVTKWTDIPY |
| ZIKV671 | PVTKWTDIPYLGKRE |
| ZIKV672 | TDIPYLGKREDLWCG |
| ZIKV673 | LGKREDLWCGSLIGH |
| ZIKV674 | DLWCGSLIGHRPRTT |
| ZIKV675 | SLIGHRPRTTWAENI |
| ZIKV676 | RPRTTWAENIKNTVN |
| ZIKV677 | WAENIKNTVNMVRRI |
| ZIKV678 | KNTVNMVRRIIGDEE |
| ZIKV679 | MVRRIIGDEEKYMDY |
| ZIKV680 | IGDEEKYMDYLSTQV |
| ZIKV681 | KYMDYLSTQVRYLGE |
| ZIKV682 | LSTQVRYLGEEGSTP |
| ZIKV683 | RYLGEEGSTPGVL |
